# VTAM: A robust pipeline for validating metabarcoding data using internal controls

**DOI:** 10.1101/2020.11.06.371187

**Authors:** Aitor González, Vincent Dubut, Emmanuel Corse, Reda Mekdad, Thomas Dechatre, Emese Meglécz

## Abstract

1. Metabarcoding studies should be carefully designed to minimize false positives and false negative occurrences. The use of internal controls, replicates, and several overlapping markers is expected to improve the bioinformatics data analysis.
2. VTAM is a tool to perform all steps of data curation from raw fastq data to taxonomically assigned ASV (Amplicon Sequence Variant or simply variant) table. It addresses all known technical error types and includes other features rarely present in existing pipelines for validating metabarcoding data: Filtering parameters are obtained from internal control samples; cross-sample contamination and tag-jump are controlled; technical replicates are used to ensure repeatability; it handles data obtained from several overlapping markers.
3. Two datasets were analysed by VTAM and the results were compared to those obtained with a pipeline based on DADA2. The false positive occurrences in samples were considerably higher when curated by DADA2, which is likely due to the lack of control for tag-jump and cross-sample contamination.
4. VTAM is a robust tool to validate metabarcoding data and improve traceability, reproducibility, and comparability between runs and datasets.

## 1 Introduction

Metabarcoding has become a powerful approach to study biodiversity from environmental samples (including gut content or faecal samples). Metabarcoding, however, is prone to some pitfalls, and consequently, every metabarcoding study should be designed in a from-benchtop-to-desktop way (from sampling to data analysis) to minimize the bias of each step on the outcome (Alberdi, Aizpurua, Gilbert, & Bohmann, 2018; Cristescu & Hebert, 2018; Zinger et al., 2019). Several papers have called for good practice in study design, data production and analyses to ensure repeatability and comparability between studies. Notably, the importance of mock community samples, negative controls, and replicates is frequently highlighted (Alberdi et al., 2018; Bakker, 2018; Cristescu & Hebert, 2018; O’ Rourke, Bokulich, Jusino, MacManes, & Foster, 2020). However, their use in bioinformatics pipelines is often limited to the verification of expectations. In this study, we present the bioinformatics pipeline, VTAM (Validation and Taxonomic Assignation of Metabarcoding data) that effectively integrates negative controls, mock communities and technical replicates to control experimental fluctuations (e. g. sequencing depth, PCR stochasticity) and validate metabarcoding data.

A recent study on the effect of different steps of data curation on spatial partitioning of biodiversity listed the following potential problems: Sequencing and PCR errors, presence of highly spurious sequences, chimeras, internal or external contamination and dysfunctional PCRs (Calderón-Sanou, Münkemüller, Boyer, Zinger, & Thuiller, 2020). They showed that exhaustive curation and ensuring repeatability by technical replicates are necessary, especially for biodiversity measurements. Ideally, a metabarcoding workflow should address all of these technical errors. Existing tools, however, are either highly flexible but too complex or they do not include the curation of all potential biases (Mahé, Rognes, Quince, de Vargas, & Dunthorn, 2014; Boyer et al., 2016; Callahan et al., 2016; Edgar, 2016b; Rognes, Flouri, Nichols, Quince, & Mahé, 2016; Bolyen et al., 2019). The filtering steps of VTAM aim to address these points and include several additional features that are unique or rarely found in existing pipelines: (i) the use of internal controls and (ii) replicates to optimize filtering parameter values; (iii) the integration of multiple overlapping markers and (iv) filtration to address cross-sample contamination, including tag-jumps. Finally, VTAM is a variant-based filtering pipeline (such as other denoising methods: Callahan et al., 2016; Edgar, 2016b) that deals with amplicon sequence variants (ASVs).

## 2 Features

### 2.1 Implementation

VTAM is based on the method described in Corse et al. 2017. It is a command-line application that runs on Linux, MacOS or Windows Subsystem for Linux (WSL). VTAM is implemented in Python 3, using a Conda environment to ensure repeatability and easy installation of VTAM and these third-party applications: Wop Mars (https://wopmars.readthedocs.io), NCBI BLAST, Vsearch (Rognes et al., 2016), Cutadapt (Martin, 2011). Data is stored in an SQLite database that ensures traceability.

### 2.2 Workflow

Table 1 summarizes the different commands and steps of VTAM, their purpose and the related error types.

**Table 1.** List of VTAM commands and their roles.

| VTAM command | VTAM step<br>(Name in Corse et al. 2017) | Role | Error Type |
| --- | --- | --- | --- |
| merge |  | Merges paired-end reads and quality filtering | Sequencing errors |
| sortreads |  | Assigns reads to samples | Sequencing errors |
| filter | Dereplicate | Dereplicates |  |
| filter | Delete singletons | Deletes singletons | Sequencing errors, highly spurious sequences |
| filter | LFN_variant filter (LFNtag) | Deletes low frequency errors | Tug jump, inter sample contamination |
| filter | LFN_read_count filter (LFNneg) | Deletes low frequency errors | Sequencing error, light contamination |
| filter | LFN_sample_replicate filter (LFNpos) | Deletes low frequency errors | Sequencing error, light contamination |
| filter | FilterMinReplicateNumber | Ensures consistency between replicates | PCR heterogeneity |
| filter | FilterPCRError (Obliclean) | Eliminates PCR errors (even if frequent) | PCR errors |
| filter | FilterChimera | Eliminates chimeras | Chimeras |
| filter | FilterRenkonen | Eliminates aberrant replicates | Dysfunctional PCRs |
| filter | FilterIndel (Pseudogene filter) | Eliminates pseudogenes | Pseudogenes, spurious sequences |
| filter | FilterCondonStop (Pseudogene filter) | Eliminates pseudogenes | Pseudogenes, spurious sequences |
| taxassign | (LTG) | Assigns variants to taxa | Highly spurious sequences |
| optimize | OptimizeLFNsampleReplicate | Finds the optimal parameter for the LFN-sample-replicate filter |  |
| optimize | OptimizePCRError | Finds the optimal parameter for FilterPCRError |  |
| optimize | OptimizeLFNreadCountAndLFN-variant | Finds the optimal value for LFN-read-count and LFN-variant filters |  |
| pool |  | Pools the results from different runs/markers |  |

#### 2.2.1 Pre-processing (optional)

An example of the data structure is illustrated in Fig. 1.

**Figure 1.**
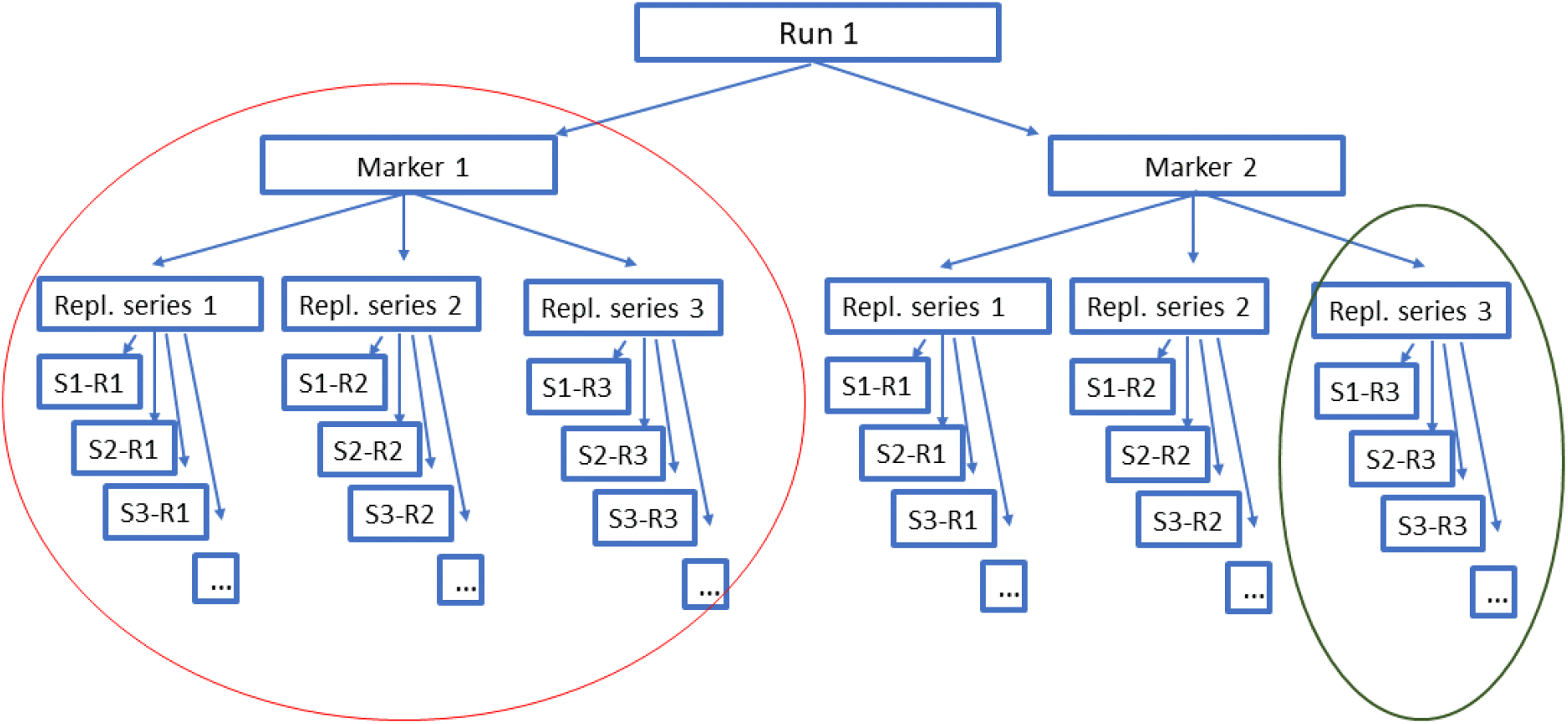
An example of a data structure with one run, two markers and three replicates for each sample. S1-R1: Replicate1 of Sample1. Replicates are not essential but strongly recommended. Samples should include at least one mock sample and one negative control.

Paired-end FASTQ files are merged, reads are trimmed and demultiplexed according to forward and reverse tag combinations.

#### 2.2.2 Filtering

Demultiplexed reads are dereplicated and ASVs are stored in an SQLite database. All occurrences are characterized by their read count.

*Filter LFN:* eliminates occurrences likely due to Low Frequency Noise. Occurrences are filtered out if they have low read counts (i) in absolute terms (N_ijk_ is small, where N_ijk_ is the read count of variant i in sample j and replicate k), (ii) compared to the total number of reads of the sample-replicate (N_ijk_/N_jk_) or (iii) compared to the total number of reads of the variant (N_ijk_/N_i_).
*Filter Min Replicate Number*: Occurrences are retained only if the ASV is pre sent in at least a user-defined number of replicates.
*Filter PCRerror*: ASVs with one difference from another ASV of the same sample are filtered out if the proportion of their read counts is below a user-defined threshold value.
*Filter Chimera* runs the uchime3_ denovo chimera filtering implemented in *vsearch*.
*Filter Renkonen* removes whole replicates that are too different compared to other replicates in the same sample.
*Filter Indel* and *Filter Codon Stop* are intended to detect pseudogenes and should only be used for coding markers. *Filter Indel* eliminates all variants, with aberrant length, where the modulo three of the length is different from the majority.
*Filter Codon Stop* eliminates all variants that have codon STOP in all reading frames of the direct strand.

The output of the filters is an ASV table with validated variants in lines, samples in columns and the sum of read counts over replicates in the cells.

#### 2.2.3 Taxonomic assignation

Taxonomic assignation is based on the Lowest Taxonomic Group method described in detail in Supporting Information 1. The taxonomic reference database has a BLAST format with taxonomic identifiers so that custom databases or the complete NCBI nucleotide database can be used by VTAM. A custom taxonomic reference database of COI sequences mined from NCBI nucleotide and BOLD (https://www.boldsystems.org/) databases is available with the program.

#### 2.2.4 Parameter optimization

Users should first identify expected and unexpected occurrences based on the first filtration with default parameters. The optimization step will guide users to choose parameter values that maximize the number of expected occurrences in the dataset and minimize the number of unexpected occurrences (false positives). Parameters are optimized for the three LFN filters and the Filter PCRerror. Optimized parameters can then be used to repeat the filtering steps.

#### 2.2.5 Pool runs/markers

A run is FASTQ data from a sequencing run and a marker is a region of a locus amplified by a primer pair. The pool command produces an ASV table with any number of run-marker combinations. When more than one overlapping marker is used, ASVs identical to their overlapping parts are pooled to the same line.

## 3 Benchmarking

VTAM was tested with two published metabarcoding datasets: a fish dataset obtained from fish faecal samples (Corse et al., 2017), and a bat dataset obtained from bat guano samples (Galan et al., 2018). Both datasets included negative controls, mock samples and three PCR replicates. A fragment of the COI gene was amplified using two overlapping markers in the fish dataset, and one in the bat dataset (See details in the original studies).

Both datasets were analysed by VTAM. The fish dataset was analysed separately for the two markers and the results of both markers were pooled together.

Both datasets were also analysed with the DADA2 denoising algorithm (Callahan et al., 2016), one of the most widely used methods for metabarcoding data curation.

The output of DADA2 was filtered by LULU (Frøslev et al., 2017) to further eliminate probable false positive occurrences. Then the three replicates of each sample were pooled (as in VTAM), only accepting the occurrence if it was present in at least two replicates (Supporting information 2).

We compared the α-diversity and β-diversity obtained for the environmental samples to address the effect of the curation pipelines on diversity estimations. α-diversity was estimated using both ASV richness and cluster richness (clusters aggregate ASVs with < 3% divergence), and β-diversity was summarized using the Bray-Curtis pairwise dissimilarity index. (Supporting information 3).

In the fish dataset, all expected variants in the mock samples were validated by both pipelines. However, in the bat dataset, two expected variants had very low read abundance (2-18 reads/ replicate), which were in the range of the number of reads in the negative controls (ten out of the 19 negative controls had at least one read count greater than 18). Therefore, we ignored these two expected variants in the Bulk France mock sample, and we optimized the VTAM parameters to retain all other expected occurrences.

After filtering with VTAM, the number of false positives in the mock samples was markedly lower than with DADA2 (Table 2). Similarly, ASV and cluster richness were on average two times lower with VTAM than with DADA 2 in environmental samples (Fig. 2A and B). In contrast, dissimilarities between samples were higher with VTAM (Fig. 2D). In both pipelines, most clusters contained a single ASV (Supporting information 3; Fig. 2C).

**Table 2.**
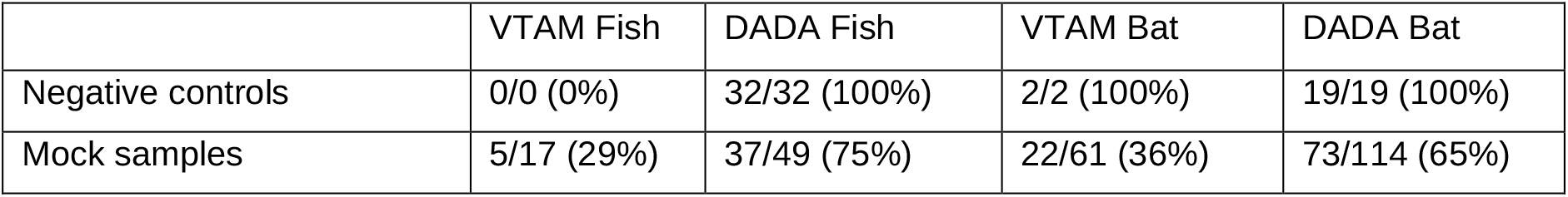
Number of false positive occurrences compared to the total number of occurrences. In negative control and mock samples, the count of false positives is precise, since the sample composition is known.

|  | VTAM Fish | DADA Fish | VTAM Bat | DADA Bat |
| --- | --- | --- | --- | --- |
| Negative controls | 0/0 (0%) | 32/32 (100%) | 2/2 (100%) | 19/19 (100%) |
| Mock samples | 5/17 (29%) | 37/49 (75%) | 22/61 (36%) | 73/114 (65%) |

**Figure 2.**
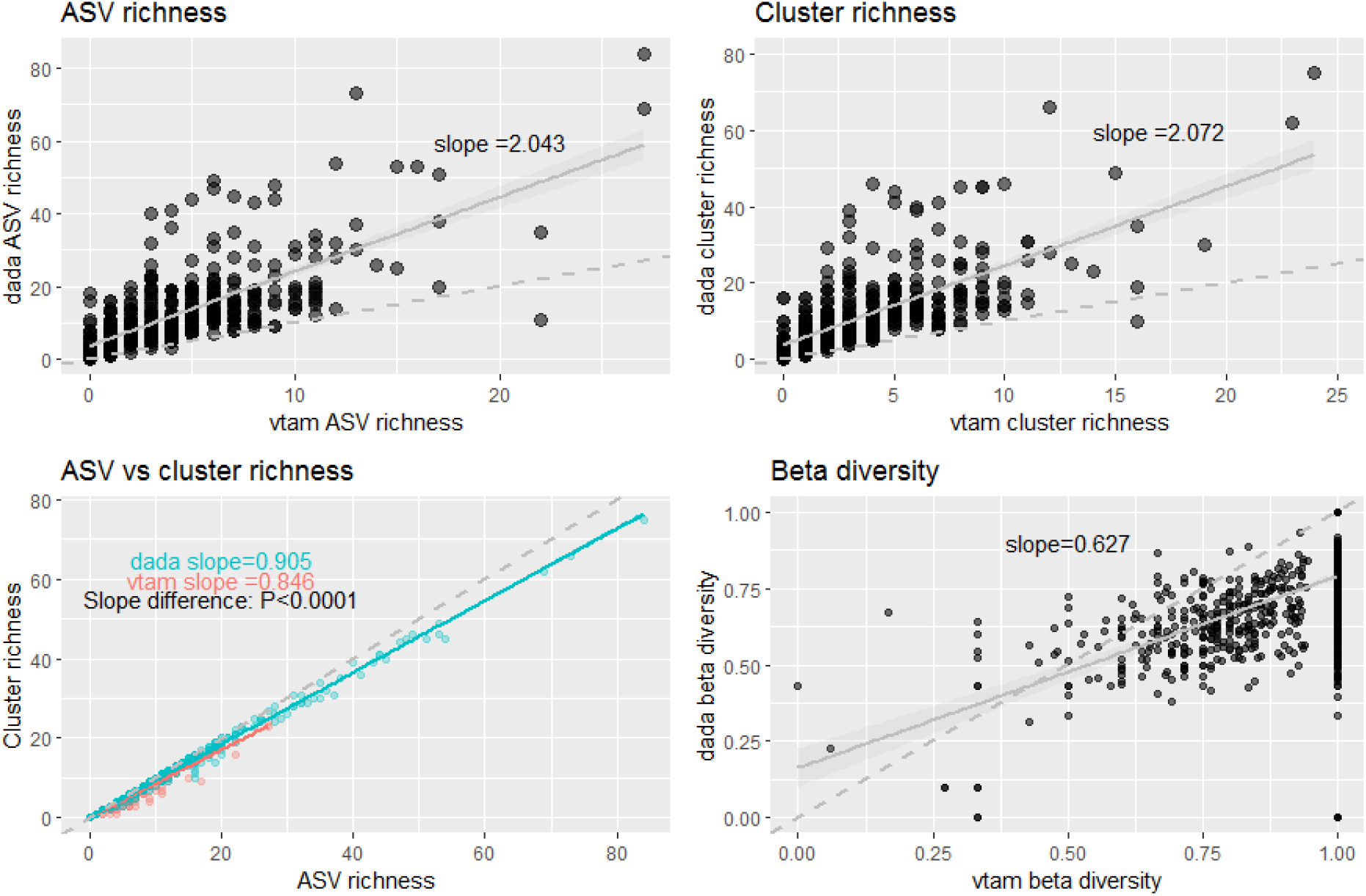
Diversity estimates from the fish and bat datasets, based on the VTAM and DADA2-based pipelines. A) ASV richness per sample B) cluster 316 richness per sample C) The correlation between ASV and cluster richness. *P-value* indicates a significant slope difference between the two pipelines. D) β-diversity was estimated using the Bray-Curtis dissimilarity index calculated for each pairwise sample comparison. Solid lines indicate linear regression lines, hatched lines are the 1:1 reference lines.

## 4 Discussion

Metabarcoding is known to be prone to two types of errors: false negatives and false positives. Based on controls (negative and mock samples), VTAM aims to find a compromise between these two error types by minimizing false positive occurrences while retaining expected variants in mock samples to avoid false negatives. Therefore, the mock samples should contain both well and weakly amplified taxa, where the abundance, i. e. the number of reads, of weakly amplified taxa is marginally higher than those found in negative samples. This should ensure finding filtering parameter values that simultaneously minimize false positives and false negatives. Additionally, in large-scale studies with more than one sequencing run, the use of identical mock samples in all runs can ensure comparability among runs if they consistently yield the same results.

The use of technical replicates is another important tool to limit false positives and false negatives (Alberdi et al. 2018, Corse et al. 2017). False positives can be strongly reduced by only accepting variants in a sample if they are present in at least a certain number of replicates. This strategy is strongly advised to reduce experimental stochasticity and validate ASV occurrences. Furthermore, removing replicates with radically different compositions (Renkonen filter) further reduces the effect of experimental stochasticity (De Barba et al., 2014). Additionally, false negatives can be further reduced by amplifying several markers (Corse et al., 2019). If the different markers overlap, VTAM can pool sequences that are identical in their overlapping regions. This integrates the results of different markers unambiguously.

While false positive occurrences due to sequencing and PCR errors are generally well detected by denoising pipelines such as DADA2, tag-jump and cross-sample contamination are rarely taken into account (but see Boyer et al., 2016; Edgar, 2016a). However, failing to filter out these artefacts is likely to inflate false positive occurrences and artificially increase inter-sample similarities. In fact, the DADA2 based pipeline produced ASV and cluster richness per sample that was on average twice as high as with VTAM and even higher for some samples (Fig. 2 A, B). On the other hand, dissimilarities between samples were lower after DADA2 filtration. Additionally, the near 1: 1 correlation between ASV and cluster richness in both pipelines indicated that most clusters contained just one ASV per sample. This supports the notion that diversity inflation in DADA2 resulted from unfiltered tag-jump contaminations rather than PCR or sequencing errors as this would have produced more ASVs that belong to the same cluster. Our VTAM pipeline, therefore, appears more appropriate for comparing the diversity between samples and for investigating the biological responses to environmental change.

## 5 Conclusions

The VTAM metabarcoding pipeline aims to address known technical errors during data analysis (Table 1) to validate metabarcoding data. It is a complete pipeline from raw FASTQ data to curated ASV tables with taxonomic assignments.

The implementation of VTAM provides several advantages such as using a Conda environment to facilitate the installation, data storage in SQLite database for traceability and the possibility to run one or several sequencing run -marker combinations using the same command. VTAM includes features rarely considered in most metabarcoding pipelines, and we believe it provides a useful tool for the analysis and validation of metabarcoding data for conducting robust analyses of biodiversity.

## Supporting information

SuppInfo1.pdf

SuppInfo2.pdf

SuppInfo3.pdf

## Acknowledgements

We thank Diane Zarzoso-Lacoste and Samanta Ortuno Miguel for valuable comments on the use of VTAM, Luc Giffon and Lionel Spinelli for the development of Wopmars and Kurt Villsen for English editing. Centre de Calcul Intensif d’ Aix - Marseille is acknowledged for granting access to its high performance computing resources. This work is a contribution to the European project SEAMo BB, fu nded by ERA-Net Mar-TERA and managed by ANR (number ANR_ 17_ MART-0001_ 01).

## Authors’ contributions

EM, EC, VD conceived the ideas and designed the methodology. EM and AG conceived the software architecture and tested the VTAM. AG, TD and RM developed the VTAM software; AG contributed to the Wop Mars software xsdevelopment. EM, AG, VD and EC wrote the manuscript. All authors contributed critically to the draft and approved the final version of the manuscript.

## Data Availability

VTAM is available at https://github.com/aitgon/vtam. A detailed user manual is found at https://vtam.readthedocs.io.

Empirical data used in this paper are available from the Dryad Digital Repository https://datadryad.org/stash/dataset/doi:10.5061/dryad.f40v5 and https://datadryad.org/stash/dataset/doi:10.5061/dryad.kv02g.

## Supporting Information

**SuppInfo1.pdf**

Description of the taxonomic assignation and its custom database.

**SuppInfo2.pdf**

Commands, user input files, and the final ASV tables produced by VTAM and the DADA based pipeline for the fish and the bat datasets.

**SuppInfo3.pdf**

Diversity estimation protocol

