## Supplementary material for "VTAM: A robust pipeline for validating metabarcoding data using internal controls": SuppInfo1.pdf

Description of the Taxonomic assignation and its custom database.

### Implementation of taxonomic assignation

Taxonomic assignation is based on the Lowest Taxonomic Group method, similar to the lowest common ancestor approach. Variants are BLASTed against the NCBI nt or a custom BLAST database. For a given identity level between the variant and the BLAST hits, the LTG will be the Lowest Taxonomic Group that contains at least a user-defined percentage (include_prop) of the hits. VTAM intends to establish LTG by using first a high %identity (100%). If LTG cannot be defined with this %identity, the %identity is decreased gradually (from 100% to 70%) till the LTG can be established. Bellow a user-defined identity level (ltg_rule_threshold, 97% by default) users can impose that the LTG should be inferred only if a minimal number of different taxa (min_number_of_taxa) are present among the hits. Users can also define a minimal coverage (min_query_coverage) between ASV and database sequences to take them into account in the taxonomic assignation.

### Custom taxonomic reference database of COI

A custom taxonomic reference database of COI sequences mined from NCBI nucleotide and BOLD (https://www.boldsystems.org/ ) databases is available with the program.

#### Seed reference dataset

The seed dataset (Appendix 1) was created by downloading all complete mitogenomes from NCBI and select the barcoding region of one species per taxonomic class.

#### Selection of NCBI COI sequences

The whole NCBI BLAST nt database was downloaded from ftp://ftp.ncbi.nlm.nih.gov/blast/db.

Sequences with COI, COXI, CO1 or COX1 in the sequence titles were extracted using the blastdbcmd command.

These sequences were aligned to the seed dataset using vsearch (Rognes, Flouri, Nichols, Quince, & Mahé, 2016) and trimmed to the target region. Sequences were retained only if they had at least 60% similarity to the most similar seed sequence, and the alignment covered at least 80 of the seed sequence.

##### Selection of BOLD COI sequences

COI-5P and COI-3P sequences were also downloaded from BOLD Public Data Portal https://v3.boldsystems.org/index.php/databases, together with their taxonomic information. As for NCBI sequences, they were aligned to the seed sequences to select the barcoding fragment. We associated each sequence with the NCBI Taxonomic identifier (TaxID) of the taxon of the lowest taxonomic level in their lineage present in NCBI Taxonomy database. This allowed us to use NCBI TaxIDs for all sequences, at the price of occasionally ignoring low-level taxa if they are not present in the NCBI Taxonomy database.

##### Creating a non-redundant database

The above selected NCBI and BOLD sequences were pooled and dereplicated within taxa: in case of identical sequences with different TaxIDs, the sequence is present in the final database once for each taxon.

#### Taxonomic reference database format

VTAM uses a simple BLAST database format, containing taxonomic identifiers. To produce this database only two files are necessary:

A fasta file with sequence identifiers, and a tsv file with sequence identifiers and their TaxIDs. The makeblastdb blast command can be used to create the indexed database files used by BLAST.

VTAM also need a taxonomy.tsv file that links each TaxID to the TaxID of its direct parent, its rank, and the name of the taxon. If all TaxIDs are present in the NCBI Taxonomy database, the Taxonomy file available in VTAM is sufficient. However, users can also complete this file with arbitrary TaxIDs, where each new TaxID should be linked to an existing parental TaxID.

Using the existing NCBI TaxID system avoids very frequent inconsistencies between lineages downloaded from different sources, and lineages that lack taxa at some main taxonomic levels. At the same time, the simplicity of the input files allows users to complete existing databases with custom sequences even from key taxa, not yet present in NCBI.

### Appendix 1

Seed Dataset to create COI reference Database

>NC_000892

TGTTCTTTATTTATTATTTGCTGGTTTTGCTGGTGTTTTAGCTGTAACTTTATCATTATTAAT
TAGATTACAATTAGTTGCTACTGGGTATGGATGATTAGCTTTGAATTATCAATTTTATAAC
ACTATTGTAACTGCTCATGGATTATTAATAGTATTTTTTCTCCTTATGCCTGCTTTAATAGG
TGGTTTTGGTAATTGAATAGTTCCTGTTCTAATTGGTTCTATTGATATGGCTTACCCTAGAT
TAAATAATATTAGTTTTTGATTATTGCCCCCTAGTTTATTATTATTAGTTGGTTCAATGTAC
ATTGAAATAGGTGCAGGTACTGGTTGAACAGTTTATCCTCCTTTAAGTTTAATCGAATTTC
ATTCAAGTGCTAGCGTTGATATGGCTATTTTTAGTCTTCATGTTTCTGGTTTGAGTAGTTTA
TTAGGTGCTATTAACTTTATTGTAACTATTTTTTGTATGAAAACTCGTGGTTTATCTTGAAG
AGCTTTACCTTTATTTGTTTGAAGTGTATTACTTACTGCTATTTTATTATTGCTAACTTTAC
CTGTTCTTGCCGGTTCTTTAACCATGTTATTAACTGATAGACATTTTAATACAAGTTTTTAT
TCAGTTTATGGTGGTGGAGATCCTGTTTTATATCAACATTTATTT

>NC_000895

AACTATGTACACAAATTTTAGTATATTGGCAGGGATAGTAGGAACTTTACTATCGTTAGTT
ATCAGAATGGAATTATCAACAGGAAACATGTTAGATGGAGACGGTCAACAATATAACGT
AATCGTAACCGCACATGGATTAATAATGATATTCTTCGTGGTTATGCCGGCAATGTTAGG
AGGATTTGCAAACTGGTTCATACCAATAATGGTAGGATCACCAGATGTAGCTTTTCCAAG
ATTAAACAACATTAGCTTATGGTTAATCATTGTATCATTTTTCTTATTACTTACATCATCAT
GTGTAGGAATAGGAGTAGGAACAGGTTGGACAGTTTATCCACCATTAAGTACGATGGAA
TATCACCCTGGACATGCAGTAGACGTAGGAATATTAAGTTTACATATAGCAGGAGCATCA
TCATTACTAGGAGCAATAAACTTTTTAACAACAGTATTTAATATGAAAATAGCAGGATTA
AGTTGGTCAAAAGTATCATTATTTGTATGGTCAATATTAATAACTGCAGTATTATTAGTAT
TATCTTTACCTGTATTAGCAGGAGGATTAACAATGTTAATTACAGATCGAAATTTTGAAA
CAACATTTTTCGATCCAATAGGAGGAGGAGATCCAATACTGTATCAGCACCTTTTT

>NC_000946

GATTTTATACTTAATTTTTGGAGCTTTTGCTGCAGTTATTGCAGTAGCACTCTCTTTATTAA
TCCGTATGGAATTAGCAGCACCTGGAGATCAAATTTTTCAAGGAAATTATCAATTATATA
ATGTAGCTATTACAGCGCATGGATTGATTATGCTTTTTTTTGTGGTTTTTCCTATTTTAGGA
GGAGGTTTTGGAAATTACTTCGTTCCTTTAATGTTAGGGGCACCTGATATGGCATTTCCTC
GTTTAAATAATATTAGTTTTTGACTTTTTCCTCCTGCTATCGCATTTTTATTATTATCTTCTT
TAGTAGAATCTGGGGCAGGTACTGGATGAACTGTGTATCCTCCTTTATCTTCTATTCAAGG
ACATTCAGGGCCTTCTGTAGATTTAGCTATTTTTAGTTTACATGTTTCTGGGGCTTCTTCTG

TTATTGGATCTATCAACTTTATTGTTACTATTTTCAATATGAGAGCACCTGGAATGTTTAT
GCATAAAATCCCTTTATTTGCTTGAGCTGTATTAATTACCGCTTTTTTACTTGTAATTTCTT
TACCTGTAGTTGCAGGGGCTATTACAATGCTTTTAACTGATCGTAATTTTAATACTACTTT
CTTTGATCCAGCTGGAGGAGGAGATCCTATTTTATATCAACATTTATTT

>NC_001131

CACCCTATATCTAATCTTTGGGGCCTGAGCAGGAATAGTGGGAACTGCTTTAAGCATCCT
AATTCGGGCAGAATTAAGTCAACCAGGCACTTTACTAGGAGACGATCAAATCTTTAACGT
TATCGTAACCGCCCATGCTTTCGTTATAATCTTTTTTATAGTTATACCAATTATAATCGGA
GGCTTCGGAAACTGACTTGTACCAATAATACTTAGCGCCCCAGATATAGCCTTCCCACGT
ATAAACAACATAAGCTTTTGACTACTTCCACCCTCACTCCTTCTACTTTTAGCTTCCGCAG
GAGTTGAAGCAGGGGCCGGAACGGGATGAACCGTATACCCACCCCTAGCAGGAAATTTA
GCCCACACAGGGGCCTCTGTTGACTTAACAATTTTCTCCCTTCACCTAGCTGGTATTTCAT
CAATTCTAGGGGCAGTCAACTTTATTACAACAATTTTTAATATAAAGCCCCCAACTATAA
CACAATACCAAATTCCTTTATTTGTTTGATCCGTTTTAATTACTGCAGTCCTCCTTCTTCTA
TCACTTCCTGTACTTGCAGCCGCCATTACTATACTTTTAACAGATCGTAATTTAAATACAT
CTTTCTTCGACCCTGCAGGGGGAGGAGACCCAATTCTTTACCAACACCTATTT

>NC_001326

ATATTTATTATTCGGATTGGTATCTGGGATAATTGGATCTGTATTCTCTTTTATAATTAGA
ATGGAACTATCAGCTCCAGGATCTCAATTCCTTTCTGGAAATGGTCAATTATACAATGTTG
CAATCTCAGCACATGGTATACTTATGATTTTTTTCTTCATTATTCCTGCTTTATTTGGTGCA
TTTGGTAATTATTTAGTACCTCTTATGATAGGTGCTCCAGATGTTGCTTACCCTAGAGTAA
ATAATTTCACATTCTGGTTACTACCTCCTGCTCTAATGCTATTATTAATTTCTGCATTAACA
GAAGAAGGACCTGGTGGTGGTTGGACGGTATATCCACCACTATCAAGTATCACTTCTCAT
AGTGGTCCAGCTATTGATCTAGCTATTTTAAGTTTACAATTAACTGGTATTTCATCAACAT
TAGGATCAGTAAATTTAATAGCGACAATGATTAATATGAGAGCTCCAGGATTAAGTTTAT
ATCAAATGCCATTATTTGCATGGGCTATAATGATTACATCAATTCTATTACTGCTTACTTT
ACCAGTTTTAGCAGGTGGATTATTTATGTTATTCTCAGATAGAAATTTAAATACTTCATTC
TATGCTCCTGAAGGTGGTGGTGACCCTGTACTTTACCAACATTTATTC

>NC_001637

TACGTTATATTTAATATTTGGTGGCTTTTCTGGTATTATAGGTACTATATTTTCTATGATTA
TAAGGTTAGAATTAGCTGCTCCGGGTTCTCAAATATTGAGTGGTAATAGCCAACTTTACA
ACGTTATTATTACGGCGCATGCCTTTGTTATGATTTCTTTTTTGTTATGCCGGTTATGATA
GGCGGTTTCGGAAATTGATTCGTTCCTTTAATGATTGGTGCTCCTGATATGGCTTTTCCAA
GGCTTAACAATATAAGTTTTTGATTACTACCGCCTTCCTTATTTCTATTATTATGTTCTTCA
TTAGTAGAATTTGGGGCAGGAACCGGCTGAACAGTTTACCCGCCGTTAAGTTCTATTGTA

GCTCACTCCGGCGGTTCCGTCGATTTAGCAATATTTAGTTTACATCTTGCAGGTATATCTT
CTCTTCTAGGCGCTATAAACTTTATTACTACGATATTTAATATGAGAGTACCTGGTTTATC
TATGCATAAATTACCATTATTTGTTTGGTCTGTGTTGATTACAGCCTTTTTGTTATTATTTT
CTCTACCTGTATTAGCAGGTGCTATTACTATGCTTTTAACTGATAGAAATTTTAATACTAG
CTTTTTTGATCCATCAGGCGGAGGCGATCCTATCCTTTATCAACATTTATTC

>NC_001660

TACTCTATATTTAATTTTCGGTGCCATTGCTGGAGTAATGGGTACATGCTTTTCAGTACTA
ATTCGTATGGAATTAGCACAACCCGGCAACCAAATTCTTGGTGGAAATCATCAACTTTAT
AATGTGTTAATAACAGCTCACGCTTTTTTAATGATCTTCTTTATGGTTATGCCGGCGATGA
TAGGTGGTTTTGGTAATTGGTTTGTTCCTATTCTTATAGGAAGTCCGGATATGGCATTCCC
TAGATTAAATAATATTTCATTTTGGCTTTTGCCACCGTCATTGTTACTTCTTCTAAGCTCAG
CCTTAGTAGAGGTGGGTTGCGGTTCAGGGTGGACGGTCTATCCACCCTTAAGTGGTATAA
CCAGTCATTCTGGAGGATCTGTTGATTTAGCAATTTTTAGTCTTCATTTATCAGGTGTTTCC
TCTATTTTAGGTTCTATTAATTTTATAACAACTATCTTCAATATGAGGGCCCCTGGATTGA
CCATGCATAGATTACCTCTATTTGTGTGGTCTGTTTTAGTAACAGCTTTCCTACTTTTATTA
TCCCTTCCAGTACTGGCAGGTGCAATTACCATGTTATTAACTGATAGAAACTTTAATACAA
CCTTTTTTGATCCTGCTGGTGGCGGGGATCCCATTTTATACCAGCATCTTTTT

>NC_001715

TACTTTATATTTAGTATTCTCAATTTTTGCTGGTATGATTGGTACTGCTTTTTCAGTTTTAA
TTAGATTCGAATTGGCTGGTCCAGGTGTACAATATTTATATGGTGATCACCAATTGTATAA
TGTTATCATCACTGCACACGCATTTATTATGATTTTCTTCCTTGTAATGCCGGCAATGCTTG
GTGGTTTCGGGAATTATTTCGTACCTATCATGATTGGTGCGCCAGATATGGCATTCCCAAG
ATTAAATAACATTAGCTTCTGGTTATTACCGCCATCATTAATTTTATTAGTTGGTTCAGCTT
TTGTTGAGCAAGGAGCCGGTACAGGATGGACAGTATACCCACCTTTATCGTCAATTGGAT
TCCATTCAGGTGGATCAGTTGATTTAGCTATCTTCAGTTTACACTTAGCTGGTATTAGTAG
TATGCTTGGAAGTATCAACTTCATCACTACTATCCTTAATATGAGGGCACCGGGGATGAC
TATGCATAAATTACCATTATTTGTATGGTCAATATTAATTACAGCTATCTTATTATTATTAT
CATTACCTGTGTTAGCTGGTGCTATTACTATGTTATTAACGGATAGAAACCTTAATACTAC
ATTCTATGATCCAGCCGGAGGTGGTGACCCAGTGCTTTATCAGCATCTTTTT

>NC_001804

TACCCTATACATGATCTTCGGTGCCTGAGCTGGAATAGTTGGAACCGCCCTAAGCCTGCTT
ATTCGAGCTGAACTCAGCCAACCTGGGGCTCTCCTGGGCGATGACCAAATTTATAATGTA
GTCGTTACAGCACATGCATTCGTGATAATCTTCTTTATAGTAATACCGATCATAATCGGCG
GGTTTGGCAACTGATTAATTCCCCTGATGATTGGGGCACCCGACATAGCATTTCCACGTAT
AAACAACATAAGCTTCTGACTACTACCACCCTCACTCCTACTCCTACTAGCATCTTCTGGA

GTAGAAGCAGGAGCAGGCACAGGATGGACAGTATACCCTCCACTAGCGGGCAACCTCGC
CCATGCAGGAGCATCCGTAGATTTAACAATTTTCTCCTTACATCTAGCCGGTGTATCCTCA
ATCTTAGGGGCCATCAACTTCATCACAACAGTAATCAACATAAAACCCCCAACAATAACA
CAGTATCAGACACCACTATTTATCTGATCAGTCTTAGTGACCGCCGTACTACTCCTACTCT
CGCTACCGGTGCTAGCTGCCGGAATTACCATACTACTGACAGATCGAAATCTAAACACAA
CATTCTTTGACCCTGCTGGAGGAGGAGACCCTATTCTATACCAACACCTATTC

>NC_001823

TGCGTTATACATAATGTTTGGTACATTTGCAGGTATTACAGCAACAACAATATCAGTAGT
AATGCGTTTAGAGCTTGGTCTTCCTGGAAATCAAATTCTTCAAGGAAATCATCAATTATAT
AATGTTTTAATAACTGCTCATGGATTACTTATGTTATTCATGGTTGTTATGCCTATCCTTTT
AGGAGGTTTTGGTAATTTCTTTGTACCATTATTAATTGGTGCACCAGATATGGCTTTCCCA
AGATTAAATAATATTAGTTTCTGGTTATTACCACCTGCATTATTACTTCTTTTCTTTTCAGC
ACTTGTAGAAGTTGGTGCAGGAACAGGTTGGACAGCATATCCACCATTATCAGGTATTCA
ATCACATTCAGGTGCTTCTGTAGATTTAGCAATATTTAGTCTTCATTTATCAGGTGCATCA
TCAGTTTTAGCTTCTATAAATTTTATTACTACTATATTCAATATGCGTGCTCCTGGTATGAC
AATGCATAGAATGCCATTGTTTGTTTGGTCAATTCTTGTAACATCATTCTTACTTGTATTTG
CATTACCTGTTCTTGCTGGTGGTATAACTATGTTATTAACAGATCGTAATTTTAATACTAC
ATTCTTTGATCCTGCAGGTGGTGGTGATCCAGTGTTATTTCAACATTTATTC

>NC_002174

AATTTTATATTTAATTTTTGGAGCTTTTTCAGGTGTTTTAGGAACAATGATGTCAATTTTAA
TTCGTATGGAGTTATCACAACCAGGAAGTCAAATTTTATCTGGTAATTATCAACTTTATAA
TGTTTTAATAACAGGTCATGCTATTTTAATGATATTTTTTATGGTAATGCCAATTATTATTG
GTGGTTTTGGTAATTTTTTAGTACCTATAATGATTGGTGCACCAGATATGGCGTTCCCTCG
TTTAAACAATCTTAGCTTTTGGTTACTTCCACCTTCATTACTATTATTATTATCATCTGCTC
TTGTTGAAGCTGGAGTTGGTACAGGATGGACAGTTTATCCTCCATTGTCTAGTGCTCAAGC
GCATTCAGGACCTTCTGTCGATTTAGCGATATTTAGTTTACATTTAGCTGGAGCGGCATCA
ATTTTAGGATCTATCAACTTTATTACTACTATTTTTAATATGAGAGCACCCGGTATGACTA
TGCATAGATTACCATTATATGTGTGGTCAGTTTTAGTTACATCTTTTTTATTAGTAATCTCT
TTACCTGTTTTAGGGGGTGCTATAACTATGCTTTTAACGGATAGAAATTTTAATACAACTT
TCTTTGATCCAGCAGGTGGGGGGGATCCTGTATTATTTCAACATTTATTT

>NC_002196

CACCTTATACCTAATCTTCGGCGCATGAGCTGGCATAGTTGGAACCGCCCTTAGCCTTCTT
ATTCGCGCAGAACTTGGTCAACCAGGAACCCTCCTAGGAGACGACCAAATCTACAACGTA
ATCGTCACTGCCCATGCTTTCGTAATAATCTTCTTCATAGTTATACCAATCATAATTGGAG
GGTTCGGAAACTGATTAGTCCCCCTTATAATCGGAGCCCCAGACATAGCATTCCCACGCA

TAAACAACATAAGCTTCTGACTACTCCCTCCGTCCTTCCTACTTCTACTAGCCTCCTCCAC
AGTAGAAGCGGGAGCAGGCACAGGATGAACCGTATACCCACCCCTAGCCGGCAATTTAG
CCCACGCTGGAGCCTCAGTAGACCTAGCCATCTTCTCCCTCCACCTAGCAGGTGTCTCCTC
AATCCTAGGAGCAATCAACTTCATCACAACCGCCATCAACATAAAACCTCCTGCCCTATC
ACAGTACCAAACCCCCCTATTCGTATGATCCGTCCTTATCACTGCCGTCCTACTACTACTA
TCTCTTCCAGTCCTCGCTGCCGGCATTACTATACTACTTACAGACCGAAACCTAAACACCA
CATTCTTTGACCCTGCTGGAGGAGGAGATCCTGTCCTATACCAACACCTCTTC

>NC_002322

GACTTTATATTTTATTTTTGGTGTGTGGTCTGGGTTGGTGGGCTTAGCTCTAAGGTTACTA
GTGCGGGCAGAATTGGGACAGCCCGGGAGAATGCTTGGAAATGACCAGCTTTATAATGTT
ATCGTTACAGCTCATGCTTTAGTTATAATTTTTTTTCTGGTTATGCCGGTTATAATCGGGG
GATTCGGGAATTGACTCATTCCTCTGATGATTGGGGCTCCAGACATGGCGTACCCTCGGA
TAAATAACATAAGGTTTTGGCTTTTGCCTCCGGCTCTTCTTTTGCTTCTTGGGTCTGCTGCA
ATGGGGGCTGGTGCAGGAACTGGTTGAACGGTGTACCCCCCGCTATCTAGGGTGGGGGCC
CATGGGGGCCCAGCTGTAGACTTGGCTATTTTTTCTTTGCATTTGGCAGGGGCTTCTTCGA
TTTTGGGGGCTATTAATTTTATTGGGTCTGTTATAAACATGAAGCCCGGCGGGTTTAAAAT
GGAGAATGTTCCGTTAATCGTGTGATCAGTTCTTAATACAGTTGGTCTTCTGTTGCTCTCT
CTTCCGGTAATGGCTGGGGCTATTACTATGCTTTTATTAGACCGAAATTTTAATACTTCTT
TTTTCGATCCGGCGGGGGGAGGAGACCCGGTTCTTTTTCAGCACTTATTT

>NC_002511

GACTCTCTATTTCATCTTCGGTGCCATTGCTGGAGTGATGGGCACATGCTTTTCAGTACTG
ATTCGTATGGAATTAGCACGACCCGGCGACCAAATTCTTGGTGGAAATCATCAACTTTAT
AATGTTTTAATAACGGCTCACGCTTTTTTAATGATCTTTTTTATGGTTATGCCTGCGATGAT
AGGTGGATTTGGGAATTGGTTTGTTCCAATTCTGATAGGTGCACCTGACATGGCATTTCCA
CGATTAAATAATATTTCATTCTGGTTGTTGCCCCCAAGTCTCTTGCTCCTATTAAGCTCAG
CCTTGGTAGAAGTGGGTAGCGGGACTGGGTGGACGGTCTATCCGCCCTTAAGTGGTATTA
CCAGCCATTCTGGAGGAGCAGTTGATTTAGCAATTTTTAGTCTTCATCTATCTGGTGTTTC
ATCCATTTTAGGTTCAATCAATTTTATAACAACTATCTTCAACATGCGTGGACCTGGAATG
ACTATGCATAGATTACCCTTATTTGTGTGGTCCGTTCTAGTAACAGCATTCCTACTTTTATT
ATCACTTCCGGTACTGGCGGGGGCAATTACCATGTTATTAACCGATCGAAACTTTAATAC
AACCTTTTTTGATCCCGCTGGGGGGGGAGACCCCATATTATATCAGCATCTTTTT

>NC_002553

TACATTATATTTAATTTTTGGTATTTTTTTCAGGTGTCATAGGTACTGTATTCTCAATAATAA
TACGTCTAGAACTTGCATTCCCTGGAAATCAAATATTAAATGGTAATCATCAATTATACA
ATGTTATTATAACAGCACACGGATTATTAATGGTATTTTTTTCAGTAACGCCTGCATTGAT

TGGTGGTTTTGGTAATTGGTTAGTACCTATTTTAATTGGAGCTCCTGATATGGCTTCTCCA
CGATCAAATAATATAAGTTCCTGGTTATTACCACCTTCTTTATTACTCCTATTATCGTCATC
ATTAATTGAAGTTGGAGCAGGTACTGGATGGACAGTATATCCTCCATTATCATCAATTCA
ATCACATTCTGGTCCCTCTGTAGACTTAGCTATTTTCAGTTTACATTTATCAGGTGCAGGT
TCTATTTTAGGAGCTGTTAATTTTATTACAACAATTTTTAACATGAGAGCTCCTGGTTTAA
CTATGAATCGCTTACCTTTATTTGTTTGGGCATTATTAATCACTGCATTTTTAATTCTATTA
TCTTTACCTGTTTTTGCAGGGGCTATCACAATGTTACTAACAGATCGTAATTTTAATACTA
CTTTCTATGATCCTGCAGGAGGTGGTGATCCAGTATTATATCAGCATTTATTT

>NC_002571

AGTTTTATATTTAATTTTTGGTGCTTTTTCTGGTGTATTAGGTACTATGATGTCTATGTTAA
TTCGAATGGAGTTATCACAACCAGGAAGTTTAATTTTAAATGGAAATTATCAACTTTATA
ACGTATTAGTTACAGGGCATGCGTTTATAATGGTTTTTTTATGGTTATGCCGGTTTTAAT
GGGTGGTTTTGGTAACTTTTTTTACCAATTTTAATTGGAGCTCCTGACATGGCTTTTCCTA
GATTAAATAATATTTCTTTTTGGTTATTGCCTCCATCATTATTATTATTATTATCTTCAGCT
TTAGTTGAAGCTGGGGCAGGTACAGGATGGACTGTATATCCACCATTAGCTGGTATACAA
GCACATTCTGGTCCTTCTATTGATTTAGCTATTTTTAGTTTACATTTAACTGGGATTTCTTC
AATTTTAGGATCTATAAATTTTATAACTACAACTTTTAATATGAGAGCGCCAGGGTTACAT
ATGCACCGAATTCCTTTATTTGTTTGGTCAATGTTAATTACTTCTTTTTTATTAATAATTTC
GTTACCTGTGTTTGGAGGTTCTATAACTATGCTATTAACAGATAGAAATTTTAATACTTCT
TTCTATGATCCTGCAGGTGGTGGTGATCCTATTTTGTATCAACATTTATTC

>NC_002572

GGTTTTATATCTTGTTTTTGCAATTTTTTCTGGTGTCGTTGGAACAGCTTTATCTATTTTAA
TAAGGGCTGAACTTTCTGGTCCAGGTGTTCAGGTTTTAGGTGGGAATCATCAATTATATA
ATGTGATTGTTACTGGTCACGCCTTTATAATGATTTTCTTTATGGTGATGCCTGCTCTAATT
GGTGGCTATGGAAATTTTTTAGTTCCTATAATGATTGGTGCCGTTGATATGGCTTTCCCAA
GAATGAATAATGTTAGTTTTTGGTTATTACCTCCAGCCTTATTATTATTAATTAGTTCAAC
ACTAACTGAGGGTGGTGCTGGTACAGGTTGGACTGTTTACCCACCTTTAAGTAGTGTCGA
AGGTCACCCAAGTGCTGCTATTGATTTAGGTATTTTTAGTTTACACGTTGCGGGTGCTTCG
TCTATCTTAGGCGCTATCAACTTTATTACAACAATATTCAATATGAGATGCCCTGGTATGA
CGTTCCATAGACTACCTCTTTTTGTTTGGGCTGTTTTGATAACTGCTTTTTTATTACTTCTA
TCTTTACCTGTTTTCGCAGGTGCTATTACAATGCTTCTTACAGATAGAAACTTTAATACGA
CGTTTTTTAATCCAGCTGGTGGGGGAGATCCTGTTTTATATCAGCATCTTTTT

>NC_002573

TATTTTATATTTATTTTTTGGAGTATTTAATGGATTTTTAGCTGTTTTATTATCTATGTTAAT
GAGACTAGAATTAGCATTTCCAGGAGACCAAATTTTATTTGGAGAATATCACTTTTATAA

CATGATTACTACTGTTCACGGAGTATTAATGCTTTTCGTAGTAGTAATGCCAATCTTATTT
GGAGGATTTGGTAACTATTTTGTACCAATTTTAATTGGTGCACCAGATATGTCTTTTCCAA
GATTAAATAATTTTAGTTTTTGGTTATTGCCTGGTGCTATTTTGTTAGCTGTATTAGCTACT
TATTCAGAAGGAGGACCAGGTACAGGATGGACAGTATATCCACCATTGTCTTCTTTACAA
TCTCACTCAGGAGCAAGCGTAGATTTAATGATATTTAGCTTTCACTTAGTAGGTATTGGAT
CTATCGTAGCAGCTATTAACTTCATTTGTACAATTTTTTATTATAAAAATGAAGCAATGTT
TAATAAAGACCTACCATTATTTGTTTGGTCTGTAGCAGTTACTTCTTTTTTAGTAATTGTAG
CTATTCCAGTATTAGCAGCAGCTATTACTTTATTATTATTTGATCGTAATTTTAATACTTCT
TTCTATGACCCAGTAGGAGGAGGGGACGTAGTATTATATCAACATTTATTT

>NC_002639

TACCCTTTACCTTATTTTCGGGGCCTGAGCCGGAATAATTGGCACAGCCCTTAGCGTAATC
ATCCGAACAGAGTTGAGTCAGCCAGGATCCTTAATCAATAATGACCAACTCTATAATACA
ATTATCACAGCCCACGCATTTGTAATAATCTTTTTTATAGTAATACCTATCATAATCGGGG
GCTTTGGGAACTGACTAGTCCCAATAATAATCGGCGCCCCTGATATAGCATTTCCCCGAA
TAAACAATATAAGCTTCTGACTTTTGCCCCCATCACTTATACTATTACTCTCCTCCTCACTA
GTAAGCTCTGGAGCAGGAACTGGATGAACAGTTTATCCCCCTCTCTCTAATCACATTTCTC
ACATAGGCCCTTCAGTAGACCTAGCTATTTTCTCCCTTCATCTAGCAGGAGTTTCCTCAAT
CTTAGGGGCAATCAACTTTATCACAACAATTATTAACATAAAAACACGGTCTATAGAAAT
ATACCATATCCCATTATTTGTATGATCAATTCTAATCACCGCAATCCTACTTCTTTTATCCT
TACCTGTTTTAGCCGCAGCTATCACTATACTCCTCACAGACCGTAATCTCAACACCACCTT
CTTTGACCCCTCTGGTGGAGGGGATCCCATCCTCTATCAGCATTTATTT

>NC_003052

TACTTAGTATCTAATCTTTGGTGGATTTGCAGGATAGATAGGGTCAGCTCTATCTATTATT
ATCCGTTTGGAATTATCTGCTGGAGGTAAAGTCTATCTTATGGGTAACTATGATCAATATA
ATGTAGTAGTTACTGCACATGGGGTAGTCATGATTTTCTTCTTGGTTATGCCTGCTCTTAT
CGGTGGTTTCGGTAACTGGCTATTACCAGTAATGGTAGGTAGTCCAGATATGGCTTTCCC
AAGATAGAATAACATTAGTTTCTGGTTATTACCGCCTTCTTTAATTTTATAGACTATTAGT
CTATTTAGTGGTGGAATGGGTACCGGATGGACAATTTATCCACCATTATCGGATACTCCA
TATCATATGGGACCTGCTGTAGATCTTGGTATTCTAAGTCTACACATTGCAGGTGTCTCTT
CTCTAATGGGTGCAATCAATCTTATTACTACTACAATTAATTGTAGAGCTCCCGGTATGAG
TTTTGAAAAATTACCTCTATTTGTATGGAGTGTATTTATTACTGCATGGTTACTATTATTAT
CATTACCTGTTTTAGCTGGAGCTATTACGATGTTATTAACTGACCGTAATCTTAATACTAG
TTTCTATGATCCTAACGGCGGAGGTGATCCGTTATTATACCAACATTAGTTC

>NC_003127

CACCCTGTATATAATCTTTGGTGCATGAGCTGGCATAGTTGGCACAGCCCTGAGCCTGCT
AATTCGAGCAGAACTAAGCCAACCAGGCGCTCTACTAGGCGATGACCAAATTTACAACGT
CCTCGTTACTGCGCACGCATTCGTAATAATTTTCTTTATAGTAATACCAATTATAATTGGC
GGCTTTGGCAACTGGCTCATTCCCCTGATAATCGGGGCCCCTGACATAGCATTTCCACGA
ATAAACAACATAAGCTTCTGGCTTCTGCCTCCATCCTTTCTCCTACTACTGGCATCTTCCG
GGGTAGAAGCCGGAGCAGGCACCGGATGGACAGTATACCCTCCACTAGCCGGCAACTTG
GCCCACGCTGGGGCATCAGTTGACCTAACCATCTTCTCGCTACACCTGGCAGGTGTGTCA
TCTATCCTCGGCTCCATCAACTTTATTACGACTATTATTAACATGAAACCACCAGCAATCT
CACAATACCAAACACCCCTCTTTATCTGATCCGTGATAATCACTACAATCTTACTATTACT
ATCTCTCCCTGTACTAGCCGCCGGAATTACTATGCTACTAACCGACCGAAACTTGAACAC
AACATTCTTTGATCCGGCAGGAGGGGGAGACCCAATCTTGTACCAACACCTATTC

>NC_004021

CACTCTATATTTAGTATTTGGTGCTTGAGCAGGAATAGTTGGTACCGCCCTAAGCCTCTTA
ATTCGAGCTGAATTAAGCCAACCAGGTACACTTCTTGGAGATGACCAAATTTATAATGTT
ATTGTAACTGCTCACGCATTTGTAATAATTTTTTTTATAGTTATGCCTGTAATAATTGGCG
GTTTCGGAAATTGATTAGTCCCATTAATAATCGGAGCCCCTGATATAGCATTTCCACGAAT
AAATAATATAAGTTTCTGACTTCTACCCCCATCATTTTTACTACTTCTAGCATCATCTGGA
ATTGAAGCCGGAGCAGGTACCGGTTGAACTGTTTACCCACCATTAGCTGGTAACTTAGCA
CATGCTGGAGCCTCAGTTGATTTAACAATTTTTTCACTTCATTTGGCAGGAATTTCATCAA
TTCTTGGAGCAATTAATTTTATCACCACTTCTATTAATATAAAACCCCCATCAATATCACA
ATATCAAACACCTCTATTTGTATGATCAGTATTAATTACAGCAATTCTACTTCTACTATCA
CTGCCAGTTCTTGCTGCAGGAATTACTATGCTTTTAACGGATCGAAACCTAAATACTACAT
TTTTTGACCCTGCTGGGGGTGGAGACCCTGTACTTTACCAACATTTATTT

>NC_004069

CACCTTGTACTTACTATTTGGTGCCTGAGCAGGCATAGTAGGAACAGCCCTAAGCCTGTT
AATTCGTGCTGAACTGGGTCAACCAGGGACCCTACTTGGGGATGACCAAATTTATAACGT
AATTGTAACCGCACATGCATTCGTAATAATTTTCTTTATAGTAATACCCATCATAATTGGA
GGATTCGGCAACTGACTAGTTCCTTTAATAATTGGTGCTCCAGACATAGCATTCCCTCGAA
TAAATAACATAAGCTTCTGGCTTCTCCCACCCTCTTTTCTACTACTTCTAGCATCATCTATA
GTTGAAGCTGGCGCAGGAACAGGCTGAACCGTATATCCCCCTCTAGCTGGTAATCTAGCC
CATGCAGGAGCTTCAGTAGATCTGACTATTTTTTCTTTACACTTAGCAGGTGTCTCTTCAA
TTTTAGGAGCCATTAACTTTATTACAACAATTATTAATATAAAACCCCCTGCCATATCACA
ATATCAAACCCCCCTGTTCGTGTGATCCGTACTAATTACTGCCGTACTTCTCCTTCTCTCAC
TTCCCGTACTAGCAGCCGGAATTACAATACTACTAACAGACCGAAACTTAAACACAACCT
TCTTTGACCCAGCAGGAGGTGGAGACCCTATTCTGTATCAACACCTATTC

>NC_004118

TACTTTATATTTAATTTTTGGTGCTATTTCTGGTGTTATAGGCACATGTTTTTCTATTCTAA
TTCGAATGGAATTAGCACAACCTGGAAATCAGATTCTTGGCGGTAACCATCAATTATATA
ATGTATTGATCACAGCGCACGCTTTTCTAATGATTTTTTTCATGGTAATGCCTGCGTTAAT
AGGAGGATTTGGTAATTGGTTCGTTCCGATCTTAATCGGCGCGCCCGACATGGCATTTCC
GCGATTAAATAATATAAGTTTTTGGTTATTACCACCTTCTTTATTATTACTTTTAAGTTCAG
CACTTGTTGAAGTAGGAGCCGGAACTGGATGGACTGTTTATCCACCTTTATCTAGTATTAC
AAGTCATTCAGGAGGTGCCGTAGATTTAGCTATTTTTAGTTTACATTTATCAGGTATCTCT
TCTATTTTAGGCTCAATTAATTTTATTACTACCATTTTTAATATGCGTGCTCCTGGAATGAC
TATGCATCGCTTGCCTTTATTTGTATGGTCTGTTTTAGTTACAGCGTTTCTTCTGTTATTAT
CATTACCTGTTTTAGCCGGTGCAATAACTATGCTTCTAACAGATAGAAATTTTAATACAAC
ATTTTTTGACCCTGCAGGTGGTGGTGATCCAGTATTATTTCAGCATCTTTTC

>NC_004309

AGTATTATACTTTATCTTTGGTAGTTTTAGTGGATTTTTAGGGACAGCTATGTCTGTGATT
ATTAGAATGGAATTAAGTATTCCTGGTAGTCCCTTTTTAGCTGGAGATTCACACTTATATA
ATGTTATCGTAACTGCCCATGCTTTTTTGATGATTTTTTTTTTGGTGATGCCTTTCTTAATG
GGAGGGTTTGGTAATTTCTTTGTTCCTTTAATGATTGGAGCTCCTGATATGAGTTTCCCTA
GAATGAACAATATAAGTTTTTGATTATTACCACCATCATTAATTTTATTAGTAGCTTCTTC
ATTAGTTGAAGGAGGAGCAGGTACTGGTTGAACGGTTTACCCACCATTATCAAGTGTAGA
ATTCCATTCAGGAGGTTCTGTAGATTTAGCTATATTTAGTTTACATTTAGCTGGAGTTTCA
TCATTATTAGGTGCTTCAAATTTTATAACAACAATATTAAATATGAGAGCTCCTGGTATGA
CAATGCATAAATTACCCTTATTTGTTTGAGCGGTATTTATTACAGCTATTTTATTATTATTA
TCATTACCTGTATTAGCTGCAGGAATAACAATGTTATTAACTGATAGAAACTTTAATACTA
GCTTTTTTGATCCTGCTGGAGGAGGAGACCCTATATTATATCAACATTTATTT

>NC_004336

AACACTGTATCTGATCTTCGCTATCTTTACTGGTCTTCTAGGTACAGCTTTCTCTGTTCTTA
TCCGACTTGAACTGTCTGGAGCTGGTAACCAATTTCTAAACGGTGACCACCAACTATACA
ATGTAATCGCTACATCTCACGGTATTTTCATGATTTACTTCATGGTAGTACCTGCTATGGC
TGGATTTGGTAACTATATGGTTCCTGTTCTTATTGGTGCTCCTGATATGGCTTTTCCACGAC
TGAATAATGTATCATTCTGGATTCTTCCTCCTGCTATTGTTCTTATCGTTTCAAGTGTATTC
GTAGAACAAGGTATGGGTACAGGTTGGACTCTTTACATGCCTATTACAGGTGTACAATCT
CACTCAGGTGGATCAGTAGATCTAGCTATTTTCTCACTACACCTTTCAGGTATCTCATCAA
TGCTAGGTGCTATGAACATTATCACTACTATTATTAACATGCGAGCTCCTGGTATGACACT
ACACAAGATGCCTCTATTTGTATGGGCTATGCTATTCCAATCAATTATCATTATCCTATGT

ATCCCAGTTCTAGCTGGTGCACTAACAATGATTATTACAGATCGAAACTTCAACACATCA
TTCTACGACCCTGCTGGTGGTGGTGACCCAGTTCTATATGAACACCTCTTT

>NC_004760

TCTTTTATATATTGTTTATTCTGTTATTGCTGGTATGGTTGGTTCTGCTTTATCTATTTTAAT
GCGTTTAGAACTTTCTTCTCCAGGTAACGCTTTCCTTGGTGGTAATCACCACCTTTATAAC
GTTATTATTACTGCTCACGGTTTAATTATGATTTTCTTCATGGTAATGCCGGGTCTTATTGG
TGGTTTTGGTAACTGGCTTTTACCTATTATGATTGGTGCTCCCGATATGGCTTTCCCTCGTC
TTAATAATTTAAGTTTCTGGTTACTTCCATCTTCTTTAACTTTACTTACCCTTAGTTTATTTG
TTGGTAATGGTGCCGGTGTAGGTTGGACTGTTTATCCTCCTTTATCTGATAGCCTTTATTCT
GGTGGTCATTCTGTAGATTTAGCCATTTTAAGTCTTCACTTAGCCGGTATTAGTTCTATGG
TTAGTTCTATGAATATGATTACTACTGTTATTAACATGCGTGCTCCAGGTATGACTATGGA
ACGTGTCCCACTTTTTGTTTGGGGTACTTTCATTACTTCTTTCCTTTTAGTTTTATCTTTACC
TGTTTTAGCTGGTGGTATTACTATGTTATTAACCGACCGTAACTTTAACACTACTTTCTAT
GACCCTACTGGTGGTGGTGACCCAATTCTTTTCCAACACCTTTTC

>NC_004815

CACCCTATATTTACTATTTGGGGCCTGAGCAGGAATAGTAGGAACCTCCCTAAGTCTACT
AATCCGAGGAGAATTAAGCTATCCAGGAACACTTATAGGAAACGACCAAATCTACAACG
TAATCGTAACAGCCCATGCTTTCACCATAATTTTTTTTATGGTAATACCAGTAATAATCGG
GGGATTTGGAAACTGACTAATTCCATTAATAATTGGAGCCCCAGACATGGCCTTCCCCCG
AATAAACAACATAAGCTTTTGACTCCTCCCACCATCATTCCTTTTACTATTGACATCCGCC
TGAACAGAAACTGGAGCAGGGACAGGATGAACCGTATACCCGCCTCTAGCAGGAAACCT
GGCCCACGCAGGACCAAGCGTAGACCTAACCATTTTTTCCCTACACTTGGCTGGCGTCTCT
TCAATCTTAGGAGCAATTAACTTCATCACCACAATCATTAATATAAAACCCCCAAACCAA
TCTCAATACCAAATACCCTTATTCATTTGATCCGTCCTAGTCACAGCAGTATTACTACTCT
TATCCTTACCCGTACTAGCAGCAGGAATTACAATACTTCTAACTGACCGAAACCTAAATA
CTTCATTCTTTGACCCATCTGGAGGAGGAGACCCCATCCTGTACCAACACTTATTT

>NC_005255

TACTTTATATCTAATCTTTGGTGCAATTGCTGGAGTAATGGGCACATGCTTCTCTGTACTA
ATTCGCATGGAATTAGCACAGCCAGGAAATCAAATTCTTGGTGGAAATCATCAACTTTAT
AATGTGTTAATAACAGCTCACGCTTTCTTAATGATCTTCTTTATGGTAATGCCTGCTCTTAT
AGGTGGGTTTGGAAACTGGTTTGTTCCTATTTTCATTGGAGCGCCTGATATGGCCTTTCCA
CGATTAAACAATATCAGTTTTTGGTTATTACCACCATCATTATTACTTCTTCTAAGTTCTGC
TTTAGTGGAAGTAGGTGCTGGTACCGGGTGGACGGTCTACCCACCTTTAAGCGGTATAAC
CAGTCATTCTGGAGGAGCTGTGGATTTAGCTATTTTTAGTCTTCATTTATCGGGTGTTTCCT
CAATATTAGGTGCTATCAATTTCATTACGACTATTTTTAACATGCGTGGCCCAGGAATGAC

CATGCATAGATTACCTCTATTTGTTTGGTCTGTGCTAATTACAGCTTTTCTGCTTTTATTGT
CACTTCCAGTATTAGCAGGTGCTATTACTATGCTATTAACTGATAGAAATTTTAATACCAC
TTTTTTTGATCCTGCAGGAGGAGGAGATCCTGTTTTATTTCAACATCTGTTT

>NC_005306

AACTTTATATTTTATTTTTGGTGCCTGAGCTGCCATGATAGGTACCGCCCTTAGAGCAATC
ATCCGAATTGAACTAGGATCCCCAGGAACATACATCAATGATGGTCACCTCTACAATGCC
ATTGTAACCGCACATGCCTTTATAATGATTTTCTTTGCAGTAATGCCAATTTTAATCGGGG
GATTCGGCAACTGGCTCGTCCCCCTTCTATTAGGGTCCCCTGATATGGCCTTTCCTCGATT
AAATAACCTAAGATTCTGACTCCTCCCCCCTTCCCTACTTCTACTGTTAGTAAGAAGGCTC
GTAGACAAGGGTGTTGGAGCAGGCTGAACTGTATACCCTCCCCTATCTGGCCTTGTCGGC
CAAAATGGCCCAAGAATGGACCTAGCAATCTTTTCCCTCCACCTTGCAGGAGCATCCTCC
TTAGCTGGAGCTATTAATTTCCTTGTCACCATCTTCCAGGCCCGATCCCCAGGTATGACCC
TTGAACGGATCCCTCTTTACCCGTGAGCAGTCGCCGTCACCGCCCTTTTGTTAGTTCTCGC
CCTTCCAGTCTTAGCTGGTGCCATTACCATGCTACTCACCGATCGTAATTTCAATACTTCA
TTCTTCTCTCCAGACGGCGGAGGAGACATAATTCTCTTTCAACATTTATTT

>NC_005332

AACTTTATATTTTGTTTTCGGAGCTTTTTCGGGAGTTTTAGGTTTTATTGTTTCAGTAGTTA
TGCGTCTAGAATTATCGCAAGCGGGTGACGCTTTTTTAGCGGGAAACTATCAATTGTATA
ATGTATTAGTTACAGCCCACGCGTTCTTAATGATCTTTTTTTTTGTTATGCCAACAGCGATT
GGAGGGTTTGGTAACTGATTTGTTCCACTAATGATTGGAGCGTGTGATATGGCTTTTCCAC
GTTTAAATAACATTAGTTTTTGATTATTACCACCTTCATTAATTCTTCTAGTATCTTCTTCT
TTAGTTGAAGCTGGGGTTGGAACTGGATGAACAGTATACCCTCCTCTAGCAGGGATTCAA
GCTCATTCAGGAGGTTCTGTTGATTTAGCAATTTTCTCTCTTCATTTAGCCGGAGTTTCGTC
GATTCTTGGTGCAATTAATTTTATTGTAACAATTATGAATATGCGAGCTCCAGGAATGACT
GCTCACCGAACTCCTTTATTTGTATGAGCTGTATTTATTACAGCTTTCTTACTTTTATTGTC
TTTACCGGTTCTAGCAGGAGCTATTACAATGCTTTTAACAGATCGTAACTTTAATACGTCA
TTCTTCGACCCTAATGGTGGAGGTGACCCAGTATTGTATCAACACTTGTTT

>NC_005840

AACTTTATATTTTATTTTGGGTGCTTGAGCTGGGATGGTTGGAACTTCTATATCAGTTTTA
ATTCGCTTGGAATTAGGCCAACCAGGGCAATTGTTAGGTGACGATCATTTATACAATGTT
GTGGTTACTGCGCATGCTTTTGTTATAATTTTTTTTATAGTGATACCTATTTTTATTGGAGG
TTTTGGAAATTGGTTGACACCTTTAATATTAGGGGTACCAGATATGGCGTTCCCTCGTTTA
AACAACCTATCTTTTTGACTTCTTCCCCCTTCTTTAATGGTTCTTCTCTCTTCTGCTTTAATC
GAAAGAGGTGTCGGAGCCGGATGAACTGTTTATCCTCCCCTTTCAAGTAATTTAGCACAT
CCTGGTGCATCTATAGACTTGGCTATTTTTTCTCTTCATATAGCTGGTGCGGCTTCTATTTT

AGGGGCTATTAATTTTATTACAACTATTATTAATATACGAAGCAAAGCTATTCGGGCGGA
CCAGATACCTTTATTTGTATGGTCAATTTTAATTACGGCTATTTTGCTATTATTGTCACTTC
CAGTGTTGGCAGGGGCTATTACTATACTTTTATGTGACCGTAACTTTAACACATCATTTTT
TGACCCTAGAGGAGGGGGTGACCCCGTTTTATTTCAGCACTTGTTT

>NC_005937

TACTATATATTTAATTTTTGGGTTTTGAGCTGGTATAGCTGGGTTTTCTTTAAGAGTTTTAA
TTCGATCTGAGTTAGGTCAGGCGGGTAGTTATATTGCAGATGATCAAATTTTTAATGTAAT
TATTACTGCTCATGCTTTTTTAATAATTTTTTTTATGGTAATGCCTGTTACTATAGGGGGTT
TTGGTAATTGATTAGTTCCTTTGATATTAGGTGCTCCTGATATAGCTTTTCCTCGTATAAAT
AATATAAGTTTTTGGTTATTATTACCATCTCTTTCTTTATTATTGTTTAGAGGTTTTGTAGA
AGTTGGAGTGGGTACAGGTTGGACGGTTTATCCCCCTCTTTCGGGGGTTTTAGGACATCCT
GGTCCTTCTGTTGATTTAGCTATTTTTTCTTTACATTTAGCTGGTGTTTCTTCTATTTTAGCT
TCTGTGAATTTTATTACAACTATGATTAATATACGAGCTGGGGTATTATCTCTGGAACGTG
TTTCTTTATTTGTGTGATCTATTTTTATTACTGCTATTTTGTTGTTATTATCTTTGCCTATTT
TGGCTGGGGCTATTACTATGTTGTTAACTGATCGTAATATTAATACTAGTTTTTTTGATCCT
ATGGGGGGTGGTGATCCTGTTTTGTATCAACATTTGTTT

>NC_005938

AACACTATATATAATCCTAGGGTTCTGAAGAGGGTTCGTAGGATTAGGACTTAGTGTCAT
TATTCGCATAGAACTAGGATCACCGGGGACAGTAATTGGGGAAGACCACACCTATAACG
TAATCGTAACAGCACACGCCCTAATTATAATCTTCTTTATAGTAATACCAATACTTATCGG
AGGATTCGGAAATTGAATATTACCAATTATATTAGGATCACCAGATATAGCATTTCCACG
AATAAACAACCTAAGATTTTGACTTCTCCCGCCATCACTACTATTATTATTAATAAGTGGA
CTAGTAGAAAGAGGAGTTGGAACAGGATGAACAATTTACCCGCCACTATCATCAAATAT
ATCACATTCAGGAGCATCAGTAGATATAGCAATCTTCTCCCTGCACCTAGCAGGAGCATC
ATCAATTTTAGGAGCCATCAACTTCATAACAACAACAATTAACATACGACCAAATCACAT
AAAACTAGAGCAAATACCATTGTTCGTATGATCAGTCTTCATTACAACAACTCTACTATTA
CTAGCATTACCAGTACTAGCAGGAGCAATCACAATATTACTAACAGACCGAAATCTAAAC
ACATCATTTTTTGACCCTACCGGTGGCGGTGACCCAATCCTATTTCAACACCTATTC

>NC_006083

AACTTTATATTTTCTATTAGGAGTATGAAGAGCTTTCGTAGGAACAGCTCTTAGAGCTCTA
ATTCGTTTAGAATTAGGAATACCTGGTAGACTTTTAGGTGACGATCAATTATACAATGTA
GTAGTAACAGCTCATGCCTTAATTATAATTTTTTTTTTGTAATACCTATTATAATTGGGG
GATTTGGAAATTGAATAACACCTTTAATAATTAATGCCCCCGATATAGCTTTTCCACGAAT
AAATAACATAAGATTTTGACTTCTTCCTCCTTCATTAATTCTACTTCTTTCCTCAAGACTAA
TTGAGAGAGGTGTAGGAACAGGGTGAACTCTGTACCCACCATTAGCTGATTTAGGGCATT

CGGGAGCTGCAGTCGATCTTGGGATTTTTTCTCTCCACTTAGCAGGGGTAAGAAGAATTC
TGGGAGCAGCAAATTTTATTACCACAATTTTAAATATACGACCTAAAGCTATAAAAATAG
AACTAGTTCCCCTTTTCGTTTGAAGAATTTTTTAACAACAATTCTTTTATTATTATCTCTA
CCTGTATTAGCAGGAGCTATTACAATACTATTAACAGATCGAAATTTTAATACCTCTTTTT
TTGATCCAGCGGGGGGAGGGGATCCAATTTTGTTTCAGCATCTTTTT

>NC_006581

GACTCTTTATTTCATCTTCGGTGCCATTGCTGGAGTGATGGGCACATGCTTCTCAGTACTG
ATTCGTATGGAATTAGCACGACCCGGCGATCAAATTCTTGGTGGGAATCATCAACTTTAT
AATGTTTTAATAACGGCTCACGCTTTTTTAATGATCTTTTTTATGGTTATGCCGGCGATGA
TAGGTGGATCTGGTAATTGGTCTGTTCCGATTCTGATAGGTGCGCCTGACATGGCATTTCC
ACGATTAAATAATATTTCATTCTGGTTGTTGCCTCCAAGTCTCTTGCTCCTATTAAGCTCA
GCCTTAGTAGAAGTGGGTAGCGGCACTGGGTGGACGGTCTATCCGCCCTTAAGTGGTATT
ACCAGCCATTCTGGAGGAGCAGTTGATTTAGCAATTTCTAGTCTTCATCTATCTGGTGTTT
CATCCATTTTAGGTTCTATTAATTTTATAACAACTATCTTCAACATGCGTGGACCTGGAAT
GACTATGCATAGATCACCTCTATTTGTGTGGTCCGTTCTAGTGACAGCATTCCCACTTTTA
TTATCACTTCCGGTACTGGCAGGGGCAATTACCATGTTATTAACCGATCGAAACTTTAATA
CAACCTTTTCTGATCCCGCTGGAGGGGGAGACCCCATATTATACCAGCATCTCTTT

>NC_006837

TACATTATATTTAATTTTTGCATTATTTTCAGGATTATTAGGTACATTATTATCATTAGTAA
TAAGATTAGAATTAATGGGTCCTGGTATACAAATTTTACAAGGTAACCATCAATTCTTTA
ATGTAGTAGTAACAGCACATGCATTTTTAATGGTATTTTTTTTATTATGCCTGCTTTAATT
GGTGGATTTGGTAATTATTTTATACCTGTTATGATAGGTGCAGTAGATATGGCATTTCCTA
GATTAAATAATATTTCATTTTGGCTATTACCGCCATCATTATTATTTTTATTAGCATCAGCT
TTTATAGAAAATGGTCCAGGTACAGGTTGGACGGTTTACCCTCCATTAGCCGGTATACAA
TCACATTCAGGGGGATCAGTAGATATTGCTATTTTTTCATTACATTTAGCAGGTATTAGTT
CTATGTTAGGTGCTATTAATATAATAACAACTGTTATTAATATGAGAAGTCCTGGTTTAAG
TTGGCATAAAATACCATTATTTGTATGGGCTGTTTTTGTAACTTCATTTTTATTATTATTAG
CATTACCTGTATTAGCAGGTGCAATTACCATGTTATTATCAGATAGAAATTTTAATAGTTC
TTTCTTTGATCCAGCTGGTGGTGGTGATCCTATTTTATATCAACATCTTTTT

>NC_006838

TACTCTATATTTAATTTTTGCGTTATTCTCAGGTATGATCGGTACTGCATTTTCTATGTTAA
TTAGATTAGAATTAGCAGGACCAGGAATACAATACTTACAAGGTGATCATCAATTATATA
ATGTAATTGTTACTGCTCATGCTTTCATTATGATTTTCTTCTTGGTTATGCCAGCTTTAATT
GGTGGATTTGGTAATTTCTTAGTACCAGTAATGATTGGTGCTGTAGATATGGCATTTCCTC
GATTAAATAATATCTCATTCTGGTTATTACCTCCAGCGTTAATCTTATTATTAGCTAGTAG

TTTTGTAGAAAATGGAGCAGGTACAGGATGGACAGTTTATCCACCTTTAGCTAGCTTACA
ATCTCATTCTGGTGGTTCTGTTGATTTAGCTATCTTCAGTTTACATCTTGCTGGGATAAGTT
CAATGTTAGGTGCTTCTAACTTCATTACTACTATTATTAATATGAGAGCTCCTGGGTTAAC
AATGCATAAATTACCATTGTTTGCTTGGGCCGTATTAATTACAGCAGTATTATTATTATTA
TCATTACCTGTATTAGCCGGTGCAATTACGATGTTATTAACAGATAGAAACTTTAATACTT
CTTTCTACGAACCAGCTGGTGGAGGTGATCCATTATTATACCAACATTTATTC

>NC_006862

ACAATTATATTTAATTTTTAGTATTTTAGCAGGAATTATTGGAACATTATTATCTTTAATA
ATTAGATTAGAGTTATCTACAGGAACAATGTTAGATGGTGATAGTCAATTATTTAATGTA
ATAATAACATCCCATGGATTAATTATGATTTTCTTCATGGTTATGCCTGCAATGTTAGGAG
GATTTGGTAATTGGTTTGTACCTATTATGATAGGTGCTCCAGATATGGCATTTCCAAGATT
AAATAATATTAGTTTTTGGTTATTAGTAGTTTCATTTATTTTATTACTTACATCATCATTTT
TAGGAATTGGAGCAGGAACAGGATGGACACTTTACCCACCATTAAGTACAATGGAATAT
CATCCAGGAGTTTCTGTAGATGTTGGAATTTTAAGTTTACATATTGCAGGAGCATCTTCTT
TATTAGGAGCTATCAATTTCGTAACTACAATTATGAATATGAAAATTGCAGGATTAAAAT
ATAGAAAAATGTCTTTGTTCGTATGGTCTGTATTAATTACTGCAATTTTATTAATTTTATCT
TTACCTGTATTAGCAGGAGGATTAACTATGTTAATTACTGATAGAAATTTTGAGACTACTT
TCTTTGATCCAATCGGAGGAGGAGATCCAATTTTATATCAACATTTATTC

>NC_007405

AACTTTATATTTAATTTTTGGAGCAATTTCAGGAGTTGCTGGTACTGCTTTATCTTTATATA
TTCGAATAACACTAGCACAGCCTAACGGTAGTTTTTAGAGTATAATCATCATCTATATAA
CGTTATTGTTACGGGTCATGCTATATTAATGATTTTTTTCATGGTAATGCCAACTCTGATT
GGAGGATTCGGTAATTGGTTTGTTCCTTTAATGATTGGTGCACCTGATATGGCTTTCCCAA
GAATGAATAATATCAGTTTTTGGTTATTACCTCCATCATTATTATTATTATTTGCATCAATG
TTAACCGAAGCTGGAGTTGGTACTGGTTGAACTGTATACCCGCCGTTGTCCAGTGCAACA
GCTCACTCTGGCGGTTCTGTAGATTTGGCTATTTTTAGTTTACATTTATCTGGCGCTTCTTC
TATTTTAGGTGCTATCAACTTTATTTGTACTATTTTCAATATGCGAGTAAAAAGTTTATCTT
TTCATAATCTTCCTTTATTTGTATGGTCAGTTTTAATTACAGCATTTTTATTATTGTTATCG
TTACCTGTATTAGCTGGTGCAATTACTATGCTATTAACAGATAGAAATTTCAATACCACAT
TTTTTGATCCTGCAGGAGGAGGTGATCCTGTGTTATTTCAACATCTTTTT

>NC_007690

TACTTTGTATTTTCTTTTTGGTTCTTGAGCTGGGATGGTAGGTACTGCTTTAAGAATAATA
ATTCGGGCAGAATTAGCTCAACCTGGTTCTTTTTTAGGGGATGATCAGATTTATAAAGTTA
TTGTTACTTCTCATGCTTTGGTAATGATTTTTTTTATGGTTATGCCTGTGATGATTGGTGGT
TTTGGAAATTGGTTAATACCATTAATGATCGGAGCTCCTGATTTAGCTTTTCCTCGTGTTA

AAAAAATGAGTTTTTGGCTTTTGCCTCCTTCTTTTCTTCTTTTATTAGCTTCTGCTGGTGTT
GAAAGAGGTGTTGGTACTGGTTGGACTATTTATCCTCCTTTGTCTAGTGGTATAGCTCATG
CTGGTGGTTCTGTTGATTTTGCTATTTTTTCTTTACATATAGCTGGAGCTTCTTCTATTATA
GCTTCTATTAATTTTATAACTACTATAATAAAAATGCGTGCTCCTGGTGTAACTTTTGATC
GTCTTTCTCTTTTTGTTTGGTCTATTTTTATTACTACTTTTCTTCTTTTATTATCTTTGCCTGT
TTTAGCTGGGGCTATAACTATGCTTCTAACAGATCGGAATGTTAATACTACTTTTTTTGAT
CCAGCTGGAGGGGGTGATCCTATTTTATTTCAGCATTTATTT

>NC_007788

GACACTATATCTAATATTTGGGGCCTGAGCAGGAATGGTTGGAACAGCCATGAGAGTAAT
AATACGAACAGAACTAGCCCAACCAGGCTCACTCCTCCAAGACGACCAAATATACAACG
TAATAGTTACAGCTCACGCTTTAGTAATGATATTCTTCATGGTAATGCCAATTATGATCGG
AGGATTCAGTAACTGACTGATCCCTCTAATGATCGGAGCACCCGATATGGCCTTCCCCCG
AATGAACAACATGAGCTTTTGACTAGTACCCCCCTCTTTTCTACTACTCCTAGCATCAGCC
GGAGTAGAAAGAGGCGCTGGAACCGGATGAACCATCTACCCTCCATTATCTAGCGGCCTA
GCCCACGCCGGAGGATCAGTGGACCTTGCAATATTCTCACTCCACCTTGCAGGAGCATCC
TCTATCCTGGCCTCCATAAAATTCATCACTACTGTAATAAACATGCGGACCCCAGGAATC
TCGTTCGACCGTCTACCACTATTCGTCTGATCAGTATTCGTAACAGCATTTCTGCTACTCC
TTTCCCTTCCCGTTCTAGCTGGAGCTATAACAATGCTTCTCACCGACCGAAACGTAAACAC
AACCTTCTTTGACCCCGCGGGGGGAGGAGACCCTATTTTATTCCAGCACCTTTTC

>NC_007886

TACTCTATATTTCATCTTCGGTGCCATTGCAGGAGTGATGGGCACATGCTTCTCCGTACTG
ATTCGTATGGAATTAGCCCGACCCGGCGATCAAATTCTTGGTGGGAATCATCAACTTTAT
AATGTTTTAATAACGGCTCACGCTTTTTTAATGATCTTTTTTATGGTTATGCCGGCGATGA
TAGGTGGATTTGGTAATTGGTTTGTTCCGATTCTGATAGGTGCACCTGACATGGCATTTCC
ACGATTAAATAATATATCATTCTGGTTGTTGCCACCAAGTCTCTTGCTCCTATTAAGCTCA
GCCTTAGTAGAAGTGGGCAGCGGCACTGGGTGGACGGTCTATCCGCCCTTAAGTGGTATT
ACCAGCCATTCTGGAGGAGCAGTTGATTTAGCAATTTTTAGTCTTCATCTATCAGGTGTTT
CATCAATTTTAGGTTCTATCAATTTTATAACAACTATCTTCAACATGCGTGGACCTGGAAT
GACTATGCATAGATTACCACTTTTTGTGTGGTCCGTTCTAGTGACAGCATTCCTACTTTTA
TTATCACTTCCGGTACTGGCAGGGGCAATTACAATGTTATTAACCGATCGAAACTTTAAT
ACAACCTTTTTTGATCCTGCAGGAGGGGGAGACCCAATATTATACCAGCATCTCTTT

>NC_008239

AACATTATATCTTATTTTTGGTGCTTTTTCTGGTGTGCTAGGAACCGCTTTCTCTCTGATTA
TTCGTATGGAATTAGCTCAACCAGGGAATCAAATTCTTGCAGGAAATCACCAGTTATATA
ACGTAATTATTACAGCGCATGCATTTTTAATGATTTTCTTCATGGTAATGCCTGTATTAAT

TGGTGGTTTTGGAAACTGGTTTGTTCCTATAATGATTGGAGCTCCAGATATGGCTTTTCCA
CGATTAAATAATATTAGTTTTTGGTTATTACCACCTTCTTTATTATTACTTTTAAGTTCTGC
TCTAGTTGAAGTTGGTGCTGGTACTGGATGGACCGTATATCCTCCGTTAAGTAGTATTCAA
TATCATTCTGGTGGATCTGTAGATTTAGCTATTTTTAGCCTACACGTATCAGGAGCTAGTA
GTATTTTAGGGGCTATTAATTTTATTACAACTATTTTTAATATGCGTGGACCTGGGATGAC
AATGCATCGATTACCTTTATTCGTATGGGCGGTGCTAATTACTGCATTTTTACTATTATTG
AGTCTTCCTGTATTTGCTGGAGCTATTACCATGTTATTAACAGATCGAAACTTTAATACTA
CATTCTTCGATCCTGCTGGAGGAGGAGACCCAATCTTATATCAACACTTATTT

>NC_008240

AACATTATATTTAATTTTCGCAGCTTTCTCCGGTATTCTAGGAACTTGTTTTTCGGTTTTAA
TTCGTATGGAATTAGCAGCGCCGGGAAATCAAATATTTGAAGGAAATCATCAACTTTATA
ATGTGTTTATTACTGCTCATGCATTCCTTATGATTTTCTTTATGGTGATGCCTGCATTAATT
GGAGGATTTGGTAATTGGTTTCTTCCTTTAATGATCGCAGCACCAGATATGGCTTTTCCAA
GACTAAATAATATTAGTTTTTGGCTATTACCACCGTCTCTTTTACTTCTTTTAAGCTCTGCT
TTAGTTGAAGTAGGCGCAGGTACTGGTTGGACAGTATATCCACCATTGAGCCATATTACA
AGTCATTCAGGAGGATCCGTGGATTTAGCTATTTTTAGTTTACATCTTTCGGGTGCTAGTA
GTATTTTAGGTGCAATTAATTTTATTACAACTGTATTTAATATGCGAGGACCTGGAATTAC
CATGAATCGTTTACCCCTATTTGTATGGGCCGTATTAATTACAGCGTTTTTACTTCTTCTGT
CATTACCTGTTTTCGCAGGAGCAATTACAATGCTATTAACTGATAGAAATTTCAATACTTC
TTTTTTTGACCCAGCGGGAGGTGGTGACCCTGTATTATACCAACATTTATTT

>NC_008453

AACTATATATTTTATTTTTGGAGTATGATCAGCTATACTAGGAACTGCTCTTAGACTAATC
ATTCGAGCAGAACTAGGACAACCTGGTTCTATAATTGGGGATGACCAAACTTATAACGTT
ATTGTTACAGCACACGCCTTCATTATAATTTTCTTTATAGTTATACCTATTATAATAGGTG
GATTTGGAAATTGATTAGTCCCTATTATAATTGGTGCTCCAGATATAGCTTTTCCTCGAAT
AAACAATATAAGATTCTGACTTCTTCCACCATCTCTATCTCTTCTTCTTGCATCCGCAGCC
GTGGAAAGAGGAGCTGGAACAGGATGAACAGTTTATCCCCCTTTAGCAAGAGAGACAGC
CCATGCTGGTGCTTCAGTAGATCTAGCAATTTTCTCACTTCATTTAGCTGGAGCCTCCTCT
ATTCTTGGCTCAGCTAACTTTATTACAACAATTATTAATATACGATCTCCAGGCATATCTT
TTGAACGAATACCCTTATTTGTATGATCAGTTCTTCTTACTGCTATTCTTCTTCTCCTAAGT
CTTCCTGTTCTAGCGGGAGCTATTACTATATTATTGACCGATCGAAATTATAATACATCCT
TCTTTAACCCAGTAGGAGGGGGGGATCCCATTCTTTATCAACATCTGTTC

>NC_008557

AACAATATATTTAATTTTTGGATTTTGATCTGCCATAGTTGGTTCAGGGTTAAGTTTAATT
ATTCGAGCTGAACTCGGGCAAGCAGGCTCCCTTCTAGGAGACGATCACCTATATAATGTG

ATTGTTACTGCCCACGCATTTGTAATGATTTTTTTATAGTAATGCCTATTTTAATTGGGGG
TTTTGGTAACTGAATGCTTCCATTAATATTAGGAGCACCAGACATAGCCTTTCCACGATTA
AACAATTTAAGTTTCTGATTATTACCACCCTCATTAACGCTTCTTTTAGCCTCTTCTTTAGT
AGAAGCAGGTGCCGGGACTGGTTGGACTGTTTACCCCCCTCTAGCTAGAAATATTGCTCA
CGCCGGTGGCTCTGTAGATTTAGCAATTTTTTCTTTACATTTAGCAGGTGCTTCTTCCATTT
TAGGGGCAGTAAATTTTATTACAACAGTGATTAATATACGTACTGAGGGGATAGTTTTTG
AGCGAATCCCGCTTTTTGTTTGAGCAGTCAAAATCACAGTTGTTTTACTTCTTCTTTCACTC
CCAGTCCTTGCTGGGGCAATCACTATATTATTGACAGATCGAAATCTAAATACTTCTTTTT
TTGATCCTGCAGGAGGAGGAGACCCTATTCTGTACCAACATCTTTTT

>NC_009458

TACTATATATTTCATTTTAGGTATTTGAGCATCCATAGTAGGAACTGCCCTGAGACTTTTA
ATTCGATTAGAATTAAGACAACCAGGAAGACTAATTGGAGATGACCAAACATACAATGT
TATTGTTACAGCACACGCTTTCGTTATAATTTTCTTTATAATTATACCAATCATAATTGGA
GGATTCGGAAATTGACTTGTTCCATTAATACTAGGAGCCCCAGACATAGCATTTCCCCGA
CTCAACAACATAAGATTTTGACTTCTTCCGCCTTCCTTAACACTCCTTTTATGCTCCGCTGC
TGTAGAAAGAGGAGCAGGAACCGGATGAACCGTATACCCTCCTCTCTCATCAAACATCTC
ACATAGAGGAGCATCAGTAGATATAACAATCTTTTCTCTACATTTAGCCGGAGTATCATC
AATTCTTGGAGCAATCAATTTCATCTCAACTATCATTAATATACGATCTAGCGGAATATCA
TTTGAACGAGTTCCATTATTTGTATGATCAGTAAAAATTACCGCAATCCTTCTTCTGCTTT
CTTTACCAGTACTAGCTGGAGCAATTACCATATTATTAACAGATCGAAATATAAATACTA
GCTTCTTTGACCCTATAGGAGGAGGAGATCCCATTTTATATCAACATTTATTC

>NC_009630

TACTTTGTATTTAATATTTGGTGGTATTGCAGGAATAATGGGTACTTGTTTTTCAATGCTA
ATTCGTATGGAGTTGTCACAGCCGGGAAATCAAATATTAGGTGGAAACTATCAGCTTTAT
AACGTGTTGATAACAGCTCATGCCTTTTTAATGATCTTCTTTATGGTTATGCCTGCTTTAAT
TGGTGGTTTTGGAAACTGGTTTGTTCCTATTTTGATAGGTGCGCCGGATATGGCCTTTCCA
CGGTTGAATAATATAAGCTTTTGGTTATTGCCACCATCCCTATTCCTTCTTTTAAGTTCAGC
CTTGGTAGAAGTTGGTGCTGGAACCGGTTGGACCGTTTATCCACCATTAAGTGGTATTACT
AGTCACTCTGGCCCTTCAGTAGATTTGGCTATTTTTAGCCTTCATTTATCTGGAGCAAGTA
GTATTTTGGGTGCTATAAACTTTATTACCACAATATTTAACATGCGCGCTCCTGGAATGAC
AATGAATCGCCTACCATTGTTTGTATGGTCTGTACTTATAACCGCATTCTTACTATTATTGT
CATTGCCTGTATTAGCCGGAGCTATTACCATGTTGTTAACAGATCGAAATTTTAATACTAC
ATTTTTTGATCCAGCTGGGGGTGGAGATCCTGTATTGTTTCAACACCTTTTC

>NC_010197

TACATTATACTTTTTATTTGGACTATGAGCTGGGATAATTGGTAGAGGTTTAAGAGCTCTT
ATTCGAGTAGAATTAAGACAACCAGGCAGCTTATTAGGTAACGACCAACTTTATAATGTA
ATTGTCACAGCTCATGCATTTTTAATAATTTTTTTATAGTCATACCAGTTATAATTGGGG
GGTTTGGAAATTGATTGGTTCCCTTGATACTGGGGGCTCCTGATATGGCATTTCCCCGCTT
AAACAACATAAGATTTTGGCTTCTCCCTCCAGCATTATTACTTCTATTAATATCTTCTTTAG
TAGAGAGGGGAGCTGGAACTGGGTGAACCGTATACCCTCCCTTATCATCGGGCTTAGGTC
ACAGAGGAGCAGCTGTGGATTTAGCCATTTTATCTTTACACCTTGCCGGTGTATCCTCAAT
TTTAGGAGCAATTAACTTTATTACTACAACAATTAATATACGAAGAAACTCTATAAAAAT
AATTCAAATCCCTTTATTTATTTGGGCAGTTTTTATTACAGCCATTTTGTTACTTTTATCTT
TACCAGTGCTAGCTGGAGCTATTACTATACTACTAACCGATCGTAATGTTAATACATCTTT
TTTTGACCCTGCTGGAGGGGGGGATCCTATTTTATACCAACACTTATTT

>NC_010202

GACTCTTTATTTATTATTTGGAGCATTTGCAGGCATGATAGGTACAGCATTTAGTATGCTT
ATAAGATTAGAGCTATCAGCCCCTGGGTCAATGTTAGGGGATGATCAATTATATAATGTT
ATAGTTACAGCCCATGCTTTTCTAATGATATTTTTCTTAGTTATGCCAGTAATGATTGGGG
GATTTGGAAATTGATTCGTGCCATTATATATTGGTGCACCCGATATGGCTTTTCCAAGATT
AAACAATATTAGTTTTTGATTATTACCTCCGGCTTTAACTCTATTATTAGGATCTGCTTTTG
TAGAGCAAGGGGTTGGTACAGGATGGACAGTATATCCCCCTTTAGCAGGCATACAAGCG
CATTCTGGGGGATCGGTTGATATGGCAATATTTAGTCTTCACTTGGCGGGTATTTCTTCGA
TATTAGGGGCTATGAATTTTATCACAACAATCTTTAATATGAGAGCGCCCGGTATTACAA
TGGATAGACTGCCATTATTTGTATGATCTATTTTAATAACAGCCTTTTTATTATTATTATCT
TTACCTGTATTAGCTGGTGGTATAACAATGCTTTTAACAGATAGAAATTTTAATACAACAT
TCTTTGATCCTGCTGGAGGAGGAGACCCAATACTATTTCAACATTTATTT

>NC_010303

GACTCTATATTCAATCTTTGGTGCCATTGCTGGAGTGATGGGCACATGCTTCTCAGTACCA
ATTCGTATGGAATTGGCACAACCCGGCGATCAAATTCTTGGTGGAAATCATCAACTTCAT
AATGTGTTAATAACGGCTCACGCTTCTTTAATGATCTCTTTTATGGTCATGCCGGCGATGA
TAGGTGGATTTGGTAATTGGTTCGTTCCGATTCTTATAGGTGCACCTGACATGGCATTTCC
ACGATTAAATAATATCTCATTCCGGTTGCTGCCACCTTCGCTGTTGCTCCTATTAAGCTCA
GCCCTGGTAGAAGTGGGTAGCGGCACTGGGTGGACGGTCCATCCGCCCTTAAGTGGTATT
ACCAGTCATTCTGGAGGAGCTGCTGATTCAGCAATTTCTAGTCTTCATTTATCAGGTGTTT
CATCCATTTCAGGTTCTATCAATTCTATAACTACTATCCCCAACATGCGCGGACCTGGAAT
GACTATGCATAGATCACCCCTATTCGTGCGGTCCGTTCCAGTGACAGCATTCCTACTTCTA
TTATCACTTCCGGTACCGGCAGGGGCAATTACCATGTTATTGACCGATCGAAACTTTAAT
ACAACCTTTTCTGATCCTGCTGGAGGGGGAGACCCGATATTATACCAGCATCTCTTT

>NC_010431

AACTCTTTATTTAATTTTTGGGGTTTGAGCAGGAATAGTTGGAACAGCTTTAAGTGTTATA
ATTCGTTTAGAATTAGGTCAACCTGGAGCTCTTCTAGGTGATGACCAACTTTATAATGTAA
TTGTTACTGCTCATGCTTTTGTAATAATTTTTTTTTTAGTAATGCCTGTTATAATTGGGGGT
TTTGGTAATTGATTAGTACCTTTAATATTAGGTGCTCCTGATATAGCATTTCCTCGTATAA
ATAATATAAGTTTCTGGTTATTACCACCTGCTTTAACTCTTCTTTTGAGAAGTTCTTTGGTT
GAGATAGGTGCAGGTACAGGTTGGACAGTCTATCCTCCTTTAGCAGGGAATCTAGCTCAC
TCTGGGGGTTCTGTTGATCTGGCTATTTTTTCTTTACATTTAGCTGGTGTTTCTTCTATTTTA
GCATCAATCAACTTTATTACTACTACTATTAATATACGTTGATATGGTTACCAGTTTGAGC
ATGTTCCTTTATTTGTATGATCAGTTAAACTAACAGCAATTTTATTGCTTTTATCTTTACCT
GTTTTAGCTGGTGGTATTACAATACTACTAACTGACCGTAATTTCAATACCTCTTTTTTTG
ACCCTTGTGGAGGAGGGGATCCAATTTTATATCAACATTTGTTT

>NC_010484

AACTTTATATTTTATTTTTGGAATCTGAGCTGGGCTCATTGGTCTTAGAATAAGATTCCTT
ATTCGTCTTGAGTTAGGGGTCGTCGGCTCTTACCTAGGTGACGAGCACCTTTATAATGTCT
TAGTGACTGCACACGCCTTTGTGATGATTTTCTTCATAGTTATGCCTGTGTCAATGGGAGG
ATTTGGTAATTGGCTGATTCCTCTGATGCTAGGCGTTGCTGACATGGCTTTCCCCCGTATA
AATAATCTATCCTTTTGGCTGTTAGTGCCAGCCTTTATGTTTCTTCTGCTGTCCTCCGCTAT
TGATGCAGGGGCCGGTACCGGTTGAACAGTGTACCCACCTTTATCGGATTCTACTTACCA
CGCAGGTGTCTCTGTTGATTTAGCGATTTTTAGCCTCCACCTATCCGGTATTTCTTCTATTC
TAGGTAGAATCAACTTCCTTACTACTATTATTTGTTCTCGTACTACAAAGAGGGTCTCTTT
AGACCGTCTGCCCTTGATGCTTTGAGCTATCGCTGTTACGGCTGTTCTCCTAATTACTAGA
CTTCCAGTGCTAGCTGGGGCTATCACAATGCTCTTAACAGATCGTAATTTTAATACTTCCT
TCTTTGATCCAGCCGGTGGAGGTAACCCTGTGTTATACCAACATCTCTTT

>NC_010637

TACATTATATCTAATTTTTGCAATTTTTGCGGGAGTTGTAGGTACTTTTTTATCGGTTTTAA
TTCGATTGGAATTAGCTGGGCCTGGCGTTCAAATATTAGGGGGTAACCACCAATTATATA
ACGTAATTATTACAGCTCATGCCTTTGTGATGATTTTTTTTATGGTGATGCCTGCACTAATT
GGAGGTTTCGCAAACTGGTTTGTTCCTATTATGATAGGTGCTCCAGATATGGCTTTTCCTC
GTTTAAATAACATTTCTTTCTGGTTATTAATACCTGCCTTCGTTTTATTATTAAGTTCATCA
TTCGTAGAAACTGGTGCGGGTACTGGCTGGACAGTGTACCCACCGTTAAGTAGTATAAGT
GGGCACCCTGGTGGATCTGTGGATTTAGCTATATTTAGCCTTCACGTTGCAGGGGCCTCA
AGTATTTTAGGTGCTTGTAATTTTATTACAACAATTCTTAATATGCGAGCACCAGGGATGA
CATTACACCGATTGCCACTTTTTTGCTGGGCAGTATTAATTACTGCGGTTTTATTAGTACT

ATCACTACCAGTATTTGCAGGGGCGATAACGATGTTGCTTACAGATAGAAATTTTAATAC
GGCATTTTTTGATGCTAGTCTTGGCGGTGACCCAGTTCTTTATGAACATCTTTTT

>NC_010651

TACTCTTTACCTTATTTTTAGAGTATTTGCAGGAATGATAGGAACAGCGTTTTCTATGCTA
ATTCGTATGGAACTAGCTGCGCCAGGTGTTCAATACCTGAATGGTGACCATCAACTTTAT
AATGTTATTATTACAGCACATGCTCTAATAATGATATTTTTTATGGTTATGCCCGCTATGG
TAGGAGGTTTTGGTAACTACCTCCTTCCAGTAATGGTTGGTAGACCAGATATGGCGTTTCC
TCGACTAAACAACATATCATTTTGGCTTCTACCCCCTTCTCTTATACTACTACTTGCTAGTG
CTTTTGTAGAACAAGGAGCAGGTACAGGGTGGACGGTTTATCCTCCACTTTCTGGTCTAC
AAAGCCATTCAGGTGGTTCGGTAGATCTTGCTATCTTTAGCCTACACCTAGCAGGGATAT
CTTCAATGCTGGGTGCTATGAACTTTATAACTACAGTTCTAAATATGCGAAATCCTGGGAT
GGACATGCATAAACTACCACTATTTGTATGGGCAATTTTTGTAACTGCCATACTACTACTT
CTTTCACTGCCAGTGCTAAGAGGTAGAATCACAATGCTACTAACAGATCGTAATTTTAAC
ACTTCATTTTATGATCCAGCAGGAGGTGGAGACCCAATACTATATCAACACCTATTC

>NC_011221

AACTTTATATATAATTTTTGGCGCATTTTCTGGAATGATAGGAACTGCTTTAAGTATGTTA
ATTAGAATAGAACTATCAACACCAGGGAGAGTAATAGGGGACGATCATTTATATAATGTT
ATAGTCACAGCTCATGCTTTTGTAATGATATTTTTTTTAGTTATGCCAGTTTTAATAGGAG
GTTATGGGAATTGATTTGTACCTATATATATAGGAGCCCCAGATATGGCTTTTCCTAGATT
AAATAATTTGAGTTTTTGATTACTTCCTCCAGCATTAATTTTACTTTTAACTTCATCCTTAG
TTGAACAAGGAGCAGGAACAGGTTGAACTGTTTACCCCCCATTATCTGGGCCTTTAGCTC
ATTCAGGAGGATCTGTAGATTTAGCTATTTTTAGTCTTCATTGTGCTGGATTTTCTTCCATT
GCAGGAGCAATTAATTTTATTACAACTATTTTTAATATGAGAACACCGGGTTTAACATTTG
ACAAACTTCCTTTGTTTGTGTGATCAGTGTTAATAACAGCATTTTTATTACTCCTTTCTTTA
CCTGTTTTAGCAGGAGCAATAACTATGCTACTAACTGATAGAAATTTTAATACTACTTTTT
TTGATCCAGCTGGAGGAGGAGACCCTGTTTTATATCAACATTTATTT

>NC_012056

GACTCTCTACCTTATTTTCGCCTTATTTTCCGGTATGATCGGAACAGCTTTTTCAATGGCCA
TACGGCTAGAGCTAGCTGCTCCAGGGATACAATACCTACAGGCTGATCATCAACTATACA
ATGTAATAATTACGGCACATGCCTTCGTTATGATCTTCTTCCTGGTCATGCCTGCCTTGAT
AGGCGGTTTTGGTAACATTTTTGTGCCACTACTTATAGGTGCCGTAGACATGGCATTTCCT
CGTCTGAATAATATTAGCTTTTGGTTACTACCTCCTGCATTAGGATTACTGCTTGCCAGTA
GCTTTGTTGAGCAAGGCGCTGGGACGGGTTGGACGGTATATCCCCCACTAGCAGGTATCA
ACTCTCACTCTGGAGGATCTGTAGACCTAGCTATCTTCAGTCTACACCTTGCTGGTATCTC
TTCTATGCTTGGTGCCATAAACTTCATAACAACCATCTTAAATATGCGAGCTCCGGGTATT

TCATTACACCGTATGCCTCTGTTTGCATGGGCTGTATTAGTGACAGCCGTTCTATTATTAT
TATCTTTACCAGTTCTAGCAGGTGGAATAACAATGTTGCTTACAGATCGAAACTTTAACA
CATCCTTCTATGAGCCTGCTGGTGGTGGCGACTTATTATTATATCAACACCTCTTT

>NC_012119

GACTCTATATTTCATCTTAGGTGCCATTGCTGGAGTGATGGGCACATGCTTCTCAGTACTG
ATTCGTATGGAATTAGCACGACCCGGCGATCAAATTCTTGGTGGGAATCATCAACTTTAT
AATGTTTTAATAACGGCTCACGCTTTTTTAATGATCTTTTTTATGGTTATGCCGGCGATGA
TAGGTGGATCTGGTAATTGGTCTGTTCCCATTCTGATAGGTGCACCTGACATGGCATTTCC
ACGATTAAATAATATTTCATTCTGGTTGTTGCCACCAAGTCTCTTGCTCCTATTAAGCCCA
GCCTTAGTAGAAGTGGGTAGCGGCACTGGGTGGACGGTCTATCCGCCCTTAAGTGGTATT
ACCAGCCATTCTGGAGGAGCAGTTGATTCAGCAATTTCTAGTCTTCATCTATCTGGTGTTT
CATCCATTTTAGGTTCTATCAATTTTATAACAACTATCTCCAACATGCGTGGACCTGGAAT
GACTATGCATAGATCACCCCTTTTTGTGTGGTCCGTTCCAGTGACAGCATTCCCACTTTTA
TTATCACTTCCGGTACTGGCAGGGGCAATTACCATGTTATTAACCGATCGAAACTTTAATA
CAACCTTTTCTGATCCCGCTGGAGGGGGAGACCCCATATTATACCAGCATCTCTTT

>NC_012651

TACTCCACATTCAATTTTCGGTGCTATTGCTGGAGTAATGGGTACATGCTTCTTAGTATCA
ATTCGTATGGAATTAGCACAACCCGGCAATCAAATTCTCAGTGGAAATCATCAACTCTAT
AATGTGTTAATAACAGCTCACGCTTTTTTAATGATATCTTTTATGGTTACGCCTGCCATGA
TAGGTGGATTTGGTAATTGGTTTGTTCCGATTCTCATAGGTGCACCTGACATGGCATTTCC
ACGATTGAATAATATCAGTTCTTGGCTATTACCGCCGTCACTGCTACTTCTTTTAAGTCCT
GCTTTGGTAGAAGTTGGCGCAGGTACCGGCTGGACGGTCTATCCGCCTCTAAGTGGTATA
ACGAGTCATTCCGGAGGAGCCGTTGATTTAGCCATTTTTAGTCTTCATCTATCAGGTGTTT
CACTTATTTCAGGTGCTATTAATCTTATCAGTACTATCTTTAATATGCGCGGCCCTGGAAT
GACTATGCATAAATTGCCTTTATTTGTGTGGCCTGTTTTAGTGACAGCATTCCTACTCTTAT
TATCCCTTCCAGTACTGGCAGGTGCCATTACCATGTTATTAACTGATAGAAACTTTAATAC
CACCTCTTTTGATCCCGCGGGAGGAGGAGATCCGATTTTATACCAGCATCCATTT

>NC_012821

TACTTTATATTTTTTATTTGGTGTTTGATCTGGTTTAGTAGGTTCTGCTTTGAGTTTATTAA
TTCGAGCTGAATTGGGACAACCTGGTGCTTTATTAGGTGATGATCAATTATATAATGTAAT
TGTTACAGCCCATGCATTTGTAATAATTTTCTTTTTAGTTATACCTGTTATAATTGGTGGTT
TCGGTAATTGATTAGTTCCTTTAATGTTAGGTGCTCCTGATATAGCTTTTCCTCGAATAAA
TAATATAAGCTTTTGATTATTGCCACCTGCTTTAACTTTACTTTTATCTTCCGGTGCTGTTG
AAAGGGGTGTAGGAACTGGATGAACAGTATACCCTCCTTTATCAGGTAATTTAGCTCATG
CTGGAGGTTCTGTAGATTTAGCTATTTTTTCTTTACATTTGGCTGGTGTTTCTTCAATTTTA

GGAGCTATTAATTTTATTACTACTATTATCAATATGCGGTGGCGAGGTATACATTTTGAAC
GTCTTCCTTTATTTGTTTGATCTGTTAAAATTACTGCTATTTTGTTATTGCTATCTTTACCTG
TTTTAGCCGGTGCAATTACTATATTGTTAACTGATCGAAATTTTAATACTGCTTTTTTTGAT
CCTGCAGGAGGAGGGGATCCTATTTTATATCAACATTTATTT

>NC_012826

AACATTATATCTTATGTTTGCATTATTTTCAGGTTTGGTTGGTACAGCTTTCTCAGTTTTAA
TTAGATTGGAGTTATCAGCTCCTGGTGTACAATATATAGCTGATAATCAATTATATAATAG
TATTATTACAGCTCATGCTATTTTAATGATATTTTTTATGGTTATGCCTGCTTTAATAGGTG
GTTTTGGTAATTTTTTATTACCATTATTAGTAGGTGGTCCAGATATGGCATTTCCTAGATT
AAATAATATAAGTTTTTGATTATTAATACCTAGTTTATTATTATTTGTATTTGCTTCTATTA
TAGAAAATGGTGCGGGTACAGGTTGAACATTATATCCTCCATTAGCAAGTATACAAAGTC
ATAGTGGTCCTAGTGTAGATTTAGCTATTTTTGGTTTACATTTAAGTGGTATAAGTTCACT
TTTAGGTGCTATGAATTTTATTACAACAATAATTAATATGAGAAGTCCAGGTATTCGTTTA
CATAAATTAGCTTTATTTGGTTGAGCCGTATTAATAACAGCTGTTTTATTATTATTATCATT
ACCTGTATTAGCTGGTGCAATAACAATGTTATTAACAGATAGAAATTTTAATACATCTTTC
TTTGAATTAGCTGGTGGTGGTGATCCTATTTTATACCAACATTTATTC

>NC_013432

TACACTCTATTTAATTTTTGGAGCGTGAGCAGGAATGGTTGGAACAGCCATGAGAGTTAT
AATCCGAACAGAACTAGCCCAGCCAGGCTCCCTTCTTCAAGACGACCAAGTTTATAAAGT
TGTGGTAACAGCCCATGCTTTAGTTATGATATTCTTTATGGTGATGCCAATAATGATTGGA
GGATTTGGCAAATGATTAATCCCTCTAATGATAGGTGCCCCAGACATGGCTTTCCCACGA
ATGAAAAAAATGAGATTTTGACTAATACCTCCCTCCTTCATTCTTCTTCTTGCCTCTGCAG
GAGTTGAAAGAGGGGCCGGAACAGGATGAACAATTTATCCTCCCCTCTCGAGCAATATTG
CCCACGCAGGAGGGTCTGTTGATCTAGCTATTTTTTCACTACACTTGGCTGGTGCCTCCTC
AATTCTAGCTTCCATAAAATTCATAACCACAATTATTAAAATGCGGACTCCGGGGATAAC
TTTTGATCGACTTCCCTTATTCGTGTGATCCGTATTTATAACTGCTATTCTTTTACTTGTGA
GCCTTCCAGTACTAGCTGGAGCCATAACGATGTTACTAACGGACCGTAAAATTAAAACGA
CTTTTTTTGACCCAGCAGGTGGAGGAGACCCAATATTGTTTCAACACTTGTTC

>NC_013476

TACATTGTATTTAATCTTTGGAGGTTTTTCAGGTGTTCTTGGAACAGCAATGTCTGTTCTTA
TTCGTTTGCAATTGGCCAGTCCTGGAAATCAATTTTTAGGGGGTAATCACCAATTATACAA
TGTTATTGTAACTGCGCATGCTTTCTTAATGATTTTTTTTATGGTTATGCCAATTCTTATTG
GTGGGTTTGGAAATTGGTTTGTACCTTTAATGATTGGTGCTCCTGATATGGCTTTTCCCCG
TATGAATAACATTAGTTTTTGGTTACTACCTCCCTCTTTAATTTTGCTTCTAGCGTCCTCGT
TGGTAGAATCTGGAGCTGGTACAGGTTGGACAGTGTACCCACCTCTTAGTGGTATTCAGG

CACACTCAGGACCTTCTGTTGACTTAGCTATATTTAGTCTTCATCTTTCGGGTGCTGCTTCT
ATTTTAGGGGCTATAAACTTTATTACAACAATTTTTAATATGAGAGCACCCGGTATGACG
ATGGATAGATTGCCCCTTTTTGTATGGTCTGTCTTAATAACAGCGTTCTTATTACTGTTATC
ACTTCCTGTTTTAGCAGGTGGTATTACAATGCTATTAACAGATAGGAATTTTAATACTACT
TTTTTTGATCCGGCCGGTGGTGGTGATCCGGTATTGTATCAGCATTTATTT

>NC_013568

AACTTTACATTTTGTTATTGGAGTTTGGTCAGGGTTTTTGGGAGCTAGAATTAGGTTGATT
ATTCGTACTGAGTTAGGTATGGTAGGAAGACTTATTATAGATGATCAGATTTATAATTCTA
TAGTAACGGCGCATGCTTTTTTAATAATTTTCTTTTTTGTCATACCAGTAGCTGTAGGGGG
GTTTGGTAATTGATTATTGCCTTTGATAATAAATGTGATAGATATGGCTTTTCCTCGTTTA
AATAATTTAAGGTTTTGATTAGTACCTGTTTCTTTAGTGTTTATGTGTATAAGATTATTAGT
GGGTTTAGGTCCTGGTACAGGTTGAACAGTTTATCCTCCTCTTTCTAATTCTGTTTATCATT
TTGGAGGGTCTATTGATTTAGCTATTTTTAGGCTTCATATTGCGGGTGTCTCTTCTATTTTA
GGGAGAATTAATTTTATTACAACATGTATGAAAGGAAAGATTAGATTTGTTATAAGTTTA
GAATACTTGAGTTTATTCACTTGAGCTATGATTGTAACAAGGTTTTTATTAGTTTTAAGTT
TACCTGTATTAGCAGGTGGGATTACTATATTATTGTTGGACCGTAATTTTGGATCTTCTTTT
TTTGATCCTAGAGGAGGAGGTAATCCTATTTTATACCAACATTTATTT

>NC_013660

GGTCTTATACCTCATGTTTGCATTGTTTTCAGGAATGCTTGGAACAGCTTATTCTGTCTTA
ATACGAATGGAATTAGCTTCTCCAGGTGTTCAATACTTACAAGGAGATAATCAATTATAT
AATGTGTTAGTTACAAGCCATGCTTTATTAATGATCTTCTTTATGGTAATGCCGGCCATGG
TGGGAGGATTTGGTAATTGGTTAGTACCAATTATGATTGGTTCTCCAGATATGGCCTTTCC
AAGATTGAATAATATTTCTTTCTGGTTATTACCACCTTCACTGATCCTATTAATTCTTTCTT
CTCTTGTAGAGGGAGGAAGTGGAACAGGGTGGACTTTCTATCCACCTCTCTCAGGAATTG
AGAGTCATTCTTCAGCAGCTGTCGATTTATCAATATTTAGTTTACATTTAGCAGGAATTAG
TTCCATGCTTGGGGCGATGAATTTTATTACAACCATCTTAAACACTTGAGCACCTGGAATG
ACAATGCATAAATTACCTCTATTTGTTTGGTCAATCTTTGTAACAGCAATTTTATTATTATT
ATCCTTACCAGTATTAGCCGGAGGTATTACAATGTTATTAACAGATAGAAATTTTAATAC
ATCATTCTATGAAGTCAGTGGAGGAGGAGATCCCTTGTTATATCAACATCTCTTC

>NC_013710

TACTTTGTATTTAATTTTTGGAGCTATCTCAGGAGTGGCTGGTACAGCGTTGTCTTTATAT
ATACGTATTACTCTATCTCAACCAGATAATGATTTCCTGTCTCACAATCACACTTTTTATA
ACGTGATCGTTACAGGACATGCTTTTATCATGATCTTTTTTATGGTAATGCCTACTCTTAT
AGGTGGTTTTGGTAACTGGTTTGTACCCATAATGATAGGTGCTCCTGATATGGCATTCCCA
AGGATGAATAATATAAGTTTTTGGATGCTTCCTCCTTCAATATCTTTGTTAATCTCTTCTGT

TTTATGTGAAGCAGGAGTAGGTACAGGTTGGACTGTCTATCCTCCTTTATCTAGTATCACG
GCTCATTCAGGTGCAGCTGTAGATCTAGCTATATTTAGTTTACATTTATCTGGAATTTCAT
CATTGTTGGGTGCAATTAATTTTATTTGTACTATTTTTAACATGCGAACAAAAAGTATGCC
TTTCCATAGATTACCTTTATTTGTTTGGTCTATTTTGATTACAGCGTTTTTATTGCTGTTGTC
ATTGCCTGTATTAGCAGGAGCTATCACTATGCTTTTAACAGATAGAAATTTCAATACAAC
GTTTTTTGAGCCTGCAGGCGGAGGTGATCCTGTACTGTTCCAACACCTTTTT

>NC_013837

AACCCTTTATTTAATTTTTGGAGGTATTGCTGGGGTTATGGGGACAACTATGTCAATTTTA
ATTCGGCTAGAGTTAGCCTACCCAGGAAGTCAAATTTTAGCAGGAAATCATCAACTTTAC
AATGTTTTAGTGACTGGTCATGCTTTTGTTATGATTTTTTTTATGGTAATGCCAGTTTTAAT
CGGAGGTTTTGGAAACTGGTTTGTTCCTTTAATGATTGGTGCACCAGATATGGCATTTCCA
CGGATGAATAATATTTCATTTTGGTTATTACCACCTTCATTATTATTACTTTTAGGGTCAAC
TTTAGTAGAAGCTGGGGCTGGTACTGGATGGACAGTTTACCCACCACTAAGTAGTGTTCA
AGCACACTCCGGACCTTCTGTAGACTTAGCAATTTTCAGTTTACATGTATCAGGAGCTGCT
TCTATTTTAGGAGCAATTAATTTTATTACAACCATTTTAAACATGAGAGCACCCGGTATGA
CAATGCATCGATTACCACTTTTTGTTTGGGCTGTATTTATTACTGCAATTTTATTATTATTA
TCTTTACCAGTTTTAGCAGGAGCCATTACTACGTTATTAACTGGTAGAAATTTTAATACAA
CTTTTTATGACCCAGCAGGAGGAGGAGATCCGGTTCTTTACCAACATTTATTT

>NC_013877

TACCCTTTATTTTCTCTTGGGGGCCTGAGCTGGGATGATCGGAACCGGCCTAAGTATTTTG
ATCCGAGCAGAACTAGCCCAACCTGGCCCACTTCTAGGAGACGACCAAATCTATAATGTT
ATTGTCACAGCCCACGCATTTGTAATGATCTTCTTTATGGTCATGCCCATAATGATAGGCG
GATTTGGAAACTGACTACTCCCCCTCATGTTAGGTGCCCCCGACATGGCATTCCCTCGCCT
CAATAATATGAGCTTCTGGCTACTCCCTCCATCCTTTCTTCTTCTTCTCTCCTCCGCCGGAG
TAGAAAGAGGAGTAGGTACTGGATGAACAGTATACCCCCCACTAGCCAGAAATATGGCT
CACGCTGGAGGATCAGTGGATCTTGCAATCTTCTCCCTACACCTTGCCGGAATTTCCTCTA
TCCTCGGGGCTATAAACTTTATGACTACAGTAATAAACATGCGAGCCCCAGGAGTTCGAT
TTGATCGCCTCCCCCTATTTGTTTGATCTGTCTTCATTACGGTAATTCTTCTTCTTCTCTCCC
TACCAGTACTCGCAGGTGCCATCACAATGCTCCTCACGGACCGAAACCTAAACACCTCAT
TCTTTGATCCAGCAGGGGGAGGGGACCCAATTCTCTACCAACACCTCTTC

>NC_013878

AACACTTTATTTAATATTTGGATCTTGAGCAGGAACAGTAGGAACAGCCATGAGAAAAAT
AATTCGAGTAGAATTATCACAACCTGGATCATTAATACAAAAAGACCAAATTTACAAAGT
AATGGTTACATCTCACGCATTAATAATGATATTCTTTATGGTAATGCCTATAATGATAGGA
GGCTTTGGAAAATGACTAGTACCTTTAATGATTGGCGCACCTGACATGGCATTTCCCCGA

ATGAAAAAAATGAGATTTTGACTAATACCGCCTTCATTCCTTCTTCTTTTAGCATCAGCTG
GAAAAGAAAGAGGAGTAGGTACTGGTTGAACTTTATACCCTCCACTTTCCGGTCCAACAG
CACATGGTGGAGGATGCGTTGACTTAGCTATATTTTCCTTACATTTAGCTGGAGCATCATC
AATAATGGCTTCAATAAAATTTATAAGAACCATTATCAACATGCGAGCACCTGGAATGAC
CTTAGACCGAACTCCCTTATTCGTCTGATCTATATTAATAACTACATTTCTATTACTTCTAT
CCCTTCCCGTTCTCGCAGGAGCAATAACCATGCTCCTTACTGACCGTAAAATAAAAACAA
CTTTCTTTGACCCTACCGGAGGAGGAGATCCTATACTCTTCCAACACTTATTT

>NC_013935

TACACTTTATATCATTTTTGGCGCGTTCTCTGGTATCCTGGGAACTTGTTTTTCGATACTTA
TTCGGATGGAGCTTTCACAGCCGGGTAATCAAGTGCTTGCAGGAAACCATAATCTTTACA
ACGTCATCATTACAGCACATGCTTTTCTTATGATTTTTTTATGGTTATGCCAACACTCGTT
GGCGGTTTTGGGAATTGGTTTGTACCGCTTCTTATTGGTGCTCCTGACATGGCTTTTCCAC
GTCTAAATAACATCAGTTTTTGGCTACTACCTCCGTCTCTCGTTCTTCTTCTTAGTTCTGCT
CTTGTGGAAGTAGGTGCTGGTACAGGTTGGACAGTATACCCCCCACTTTCTAGTATTCTTG
CTCATTCTGGGGGGGCGGTTGATCTTGCAATTTTTTCTCTTCACCTTTCTGGAATTTCGTCT
ATTCTAGGAGCTATTAATTTTATTACAACCGTTTTTAATATGCGCGCTCCTGGTATGGCTA
TATATCGTCTTCCGCTTTTTGTTTGGTCAATTCTCATTACAGCGTTTCTTCTTCTCCTTAGTC
TGCCTGTTCTTGCGGGAGCTATTACTATGCTTCTCACTGACCGAAATTTCAATACAAGCTT
TTTTGACCCTGCTGGTGGTGGAGACCCTATTCTGTATCAACACCTTTTT

>NC_013986

TACTCTGTATTTAATATTTGGTATTTTAGCCGGCGTTGTTGGTACGGTTTTATCTATTTTTA
TAAGAATGGAGCTTTCCGCCCCCGGTGATCAAATTTTAGCTGGTAATTATCAGTTATATAA
TGTTATTGTAACAGCTCACGCTTTTGTTATGATATTTTTTATGGTAATGCCAGCGCTAATA
GGTGGCTTTGGTAATTGGTTTGTACCTTTATTAGTCGGTGCACCTGATATGTCTTTTCCAC
GATTAAACAACTTAAGTTTTTGGTTGTTACCTGTTTCTTTAAGCTTGCTTTTACTATCTAGT
TTTGTTGAAGTTGGTGCTGGTACTGGTTGAACGGTTTACCCACCACTAAGTTCTATTCAGG
CTCACTCTGGTGGCTCGGTTGATTTAGCTATATTTAGTTTACACGTTTCGGGTGTTTCATCT
TTACTTGGTGCAATAAACTTTATATGTACTGTTTTTAATATGCGTGTTCCGGGGTTATATA
TGCATAGATTACCTCTTTTTGTTTGGGCGGTTTTAATAACGGCGTTTCTACTACTTTTGTCT
CTTCCTGTTTTAGCGGGTGGTATTACAATGCTTTTGACTGATCGAAATTTTAATACCACTT
TTTTTGATTCTGCAGGTGGTGGTGATCCTATATTGTATCAACATTTATTT

>NC_014043

GACTCTATATTTCATCTTCGGTGCCATTGCTGGAGTGATGGGCACATGCTTCTCAGTACTG
ATTCGTATGGAATTAGCACGACCCGGCGATCAAATTCTTGGTGGGAATCATCAACTTTAT
AATGTTTTAATAACGGCTCACGCTTTTTTAATGATCTTTTTTATGGTTATGCCGGCGATGA

TAGGTGGATCTGGTAATTGGTCTGTTCCGATTCTGATAGGTGCACCTGACATGGCATTTCC
ACGATTAAATAATATTTCATTCTGGTTGTTGCCACCAAGTCTCTTGCTCCTATTAAGCTCA
GCCTTAGTAGAAGTGGGTAGCGGCACTGGGTGGACGGTCTATCCGCCCTTAAGTGGTATT
ACCAGCCATTCTGGAGGAGCAGTTGATTCAGCAATTTCTAGTCTTCATCTATCTGGTATTT
CATCCATTTTAGGTTCTATCAATTTTATAACGACTATTTCCAACATGCGTGGACCTGGAAT
GACTATGCATAGATCACCCCTCTTTGTGTGGTCCGTTCTAGTGACAGCATTCCCACTTTTA
TTATCACTTCCGGTACTGGCAGGGGCTATTACCATGTTATTAACCGATCGAAACTTTAATA
CAACCTTTTCTGATCCCGCTGGAGGGGGAGACCCCATATTATACCAGCATCTCTTT

>NC_014505

AACAATATATTTAATTTTTGGAGTTTGATCAGCGATAATGGGGACAGCTTTGAGAATATT
AATTCGAATTGAGTTAGGAACTCCTTCTTCATTAATTGGGGATGATCAAATTTATAATGTT
ATTGTCACTTCTCACGCTTTTGTAATAATTTTTTTTATAGTTATACCAATAATAATTGGGGG
TTTTGGTAATTGGTTAGTTCCTATTATAATTAATGCTCCTGATATAGCTTTCCCACGAATA
AATAATATAAGATTTTGGTTACTTCCTCCTTCTTTATTATTATTAATTTCTTCTTCCATAGT
TGAAATAGGAGCTGGGACAGGATGAACAGTCTATCCTCCTTTATCTAGAAGTCTTGCTCA
TTCAGGTGCATCTGTAGATTTAACTATTTTTTCTTTACATTTAGCAGGGGTTTCTTCAATTT
TAGGAGCTATTAATTTTATTAGAACAATTTTAAATATGCGTACTGTAGGTATGAAATTAG
AACAAATACCTTTATTTGTTTGAAGAGTTTTAATTACAGCTGTGTTATTATTATTATCATTA
CCAGTGTTAGCAGGAGCTATTACTATACTTTTAACAGATCGAAATTTTAATACTTCATTTT
TTGACCCAGCAGGAGGTGGGGACCCAATTTTGTACCAACATTTATTT

>NC_014853

TACACTATATTTATTATTTGGTGCCTTCGCTGGTATGATTGGTACAGCGTTTAGTATGTTG
ATAAGATTAGAATTATCAGCACCCGGAACAATGTTAGGAGATGATCAATTATATAACGTA
ATAGTTACTGCTCATGCGTTTGTAATGATTTTCTTTCTAGTTATGCCTGTTATGATAGGAG
GGTTTGGTAATTGATTATTACCATTATATATAGGAGCCCCAGATATGGCTTTTCCTCGATT
AAATAACATAAGCTTCTGATTATTACCACCTGCTTTAATTTTATTATTAGGTTCAGCTTTTG
TAGAACAAGGGGCAGGGACAGGGTGAACAGTATATCCACCATTAGCTGGGATACAATCA
CACTCAGGAGGATCAGTAGATATGGCAATATTTAGTCTTCATTTAGCAGGGGCTTCATCA
ATACTTGGTGCAATAAATTTTATTACAACTATATTTAATATGCGAGCTCCAGGAATAACA
ATGGATCGACTTCCATTATTTGTTTGATCAATTTTAATTACTGTATTTTTACTATTATTATC
TTTACCAGTCTTAGCAGGTGCTATAACTATGTTATTGACTGATCGAAATTTTAATACAACT
TTCTTTGACCCAGCTGGAGGTGGAGATCCTATATTATATCAACATTTATTC

>NC_015121

GACTCTATATTTCATCTTCGGTGCCATTGCTGGAGTGATGGGCACATGCTTCTCCGTATTG
ATTCGTATGGAATTAGCACGACCCGGCGATCAAATTCTTGGTGGGAATCATCAACTTTAT

AATGTTTTAATAACGGCTCACGCTTTTTTAATGATCTTTTTTATGGTTATGCCGGCGATGA
TAGGTGGATCTGGTAATTGGTCTGTTCCGATTCTGATAGGTGCACCTGACATGGCATTTCC
ACGATTAAATAATATTTCATTCTGGTTGTTGCCGCCAAGTCTCTTGCTCCTATTAAGCTCA
GCCTTAGTAGAAGTGGGTAGCGGCACTGGGTGGACGGTCTATCCGCCCTTAAGTGGTATT
ACCAGCCATTCTGGAGGAGCAGTTGATTCAGCAATTTCTAGTCTTCATCTATCTGGTGTTT
CATCCATTTTAGGTTCTATCAATTTTATAACAACTATCTCCAACATGCGTGGACCTGGAAT
GACTATGCATAGATCACCCCTATTTGTGTGGTCCGTTCCAGTAACAGCATTCCCACTTTTA
TTATCACTTCCGGTACTGGCAGGGGCAATAACAATGTTATTAACCGATCGAAACTTTAAT
ACAACCTTTTCTGATCCCGCAGGAGGGGGAGACCCCATCTTATACCAGCATCTCTTT

>NC_015235

CACCTTGTATTTTATTTTCGGCGCCTGAGCCGGAATAGTAGGCACAGCCATAAGCCTATTA
ATCCGAACAGAGCTCAGCCAGCCAGGTCCCTTCATAGGAGATGACCAAATTTATAATGTT
ATTGTTACAGCACATGCCTTTATCATAATTTTCTTTATAGTTATACCAATTATGATCGGGG
GATTTGGAAATTGACTACTCCCATTAATAATTGGGGCACCCGACATAGCATTCCCTCGCA
TAAACAACATAAGCTTCTGATTGCTGCCCCCATCATTTACCCTACTTCTCTTTTCCGCCTTT
ATTGAAACTGGGGCTGGCACCGGATGAACAGTCTACCCACCACTAGCTGGAAACCTAGCC
CACGCCGGACCGTCAGTAGACCTCACTATCTTCTCCCTTCACCTTGCTGGGGTATCATCCA
TCCTTGGAGCAATCAACTTTATTACCACAGCTATCAACATAAAACCCCCAGCAATGTCAC
AACAACAAACACCCCTTTTTGTATGGTCTGTTCTAGTTACAGCTGTTCTCCTACTGCTCTC
ACTACCCGTCCTAGCTGCAGGAATTACCATATTACTTACTGACCGAAACTTGAACACCAC
TTTCTTTGACCCGGCAGGAGGAGGAGACCCAATCCTATACCAACACCTTTTC

>NC_015619

AACTTTATATTTAATTTTTAGTGCTTTTGCTGGTGTTGTTGGTACAACATTTTCTCTTTTAA
TTAGAATGGAATTAGCACAACCAGGTAATCAAATTTTTATGGGAAATCATCAATTATATA
ATGTTGTTGTTACCGCACATGCTTTTATTATGGTTTTCTTTTTAGTTATGCCTGCTTTAATC
GGTGGTTTTGGTAATTGGTTTGTTCCTTTAATGATAGGTGCTCCTGATATGGCTTTTCCTCG
TATGAATAATATTAGTTTTTGGTTATTACCTCCTTCTTTATTATTATTAGTTTCTTCAGCTA
TCGTTGAATCTGGGGCTGGTACTGGTTGGACAGTTTATCCACCATTATCTAGTGTTCAAGC
ACATTCAGGACCTTCTGTAGATTTAGCTATTTTTAGTTTACATTTATCAGGTATTTCTTCTT
TATTAGGTGCTATTAATTTTATTTCAACAATTTATAATATGAGAGCTCCTGGTTTAAGTTTT
CATAGACTACCTTTATTTGTATGGTCTATATTAATTACTGCATTTCTTTTATTATTAACTTT
ACCTGTACTAGCTGGGGCAATTACTATGTTACTAACTGATAGAAATTTAAACACTTCATTT
TATGATCCATCAGGTGGAGGTGATCCAGTATTATATCAACATTTATTT

>NC_015646

AACTATTTATTTTATTTTTGGAATATGGGCAGGTTTATTAGGAAGGGCTCTTTCTTATTGA
ATTCGAGTAGAACTTTCTCAGCCTGGATCTTTATTAAAAGACGAACAGTTATATAACTGT
ATTGTTACAGGTCATGGGTTAATTATAATTTTCTTTTTTGTTATACCAGTTATAATTGGAG
GGTTTGGAAATTGATTAATCCCATTAATGTTAAAAACTCCCGATATAGCCTTTCCTCGGAT
AAATAATATAAGGTTTTGATTATTACCTCCTTCATTAACTTTATTATTATGTTCATTTTTAG
TTGAATCTGGAAGAGGTACAGGGTGAACTATATATCCTCCTTTATCTGATAGGTTAGCAC
ACAGAAGGAAAAGAGTAGACTTAACAATCTTCTCCCTTCATTTGGCAGGAGTATCATCAA
TTTTAGGTGCTATTAATTTTATTACAACTATACTTAATATACGGCCTCGAGGTATAAAAAT
ACATTATTTACCATTATTATGTTGATCTATTTTAATTACAGCAATTTTATTACTACTTTCTT
TACCAGTTTTAGCAGGAGCAATTACTATGCTATTAACAGACCGAAATTTTAATACTTCATT
TTTTGATGCTAGAGGTGGGGGAGATCCTATTTTAATGCAAGGGTTATTC

>NC_015649

TTTTATTTCTGCTATTTCTGGGGGAGGGCTGAGTATACAAATGCGACGTAGTTTGAGAACT
TATTCTTGGTGTGAATTTTCTTCTATTTTGGAGGGTGGTTTAAGTATCACTTCTGATCATTA
TAATTCTATGGTTACGGCTCATGGGTTAGTTATGATTTTTTTTTTATTATGCCTGTTATGA
TTGGAGGTTTTGGGAATTGGTTAATTCCTTTGATGATTGGGTCTGCTGATATGGCTTTCCC
TCGGCTTAATAATTTAAGTTTTTGACTTTTACCACCAGCTTACTTTATGCTTTGGGTGGGTT
TATTTTCTGGTAAGGTTGGGATAGGTTGGACTATTTATCCCCCTCTAAGAGGGGGAAACTT
TTCTTCGGGGTGATTCGGGGACTTTCTTTTAATTTCTTTACATATTGCTGGGGCGGGTTCA
ATTTTGGGGGGTATAAATTTTATTACTACTATGAGACAGTTGCGGGTTAGTGGTATGAGTT
TTAATAGGTTGCCTGTTTTTTGTTGGGCTTTACTTGTAGCTTCTGTTTTGTTGGTTGTTGCT
ATGCCTGTTTTGGCTGGGGCTATTTCTATGTTGTTAGCGGACCGTCATTTTGGTGGGAGTT
TTTTTGATCCTTTGGGGGGTGGTGACCCTATTCTTTGACAACATTTATTT

>NC_015789

TACATTATATCTAATATTTGCTTTATTTTCTGGATTATTAGGTACTGCTTTTTCTGTCTTAA
TAAGATTAGAATTAAGTGGACCTGGTGTTCAATATATTGCAGATAATCAACTATATAATA
GTATCATTACAGCTCATGCTATTATAATGATATTCTTTATGGTTATGCCTGCTTTAATAGG
AGGTTTTGGTAATTTCTTATTACCATTATTAGTAGGAGGTCCAGATATGGCCTTTCCTAGA
CTTAACAACATTAGTTTCTGATTATTACCACCTAGTTTATTATTATTTTTATTTGCTAGTGG
TATAGAAAACGGGGCAGGTACTGGTTGAACATTATACCCACCTTTAAGTGGAGTACAAAG
TCACAGTGGGCCTAGTGTTGATTTAGCTATATTTGGGCTACACTTATCTGGTATAAGTAGT
TTATTAGGAGCTATAAACTTCATTACTACTATATTAAATATGAGAAGTCCTGGTATTAGAC
TACACAAGTTAGCTTTATTTGGATGAGCTGTAGTAGTTACAGCTGTATTATTATTATTATC
ATTGCCTGTGTTAGCCGGAGGTATAACTATGGTTTTAACTGATAGAAACTTTAATACATCA
TTCTTTGAAGCAGCAGGAGGTGGTGACCCTATACTATACCAACATCTTTTC

>NC_015890

AACAATATATTTTATTTTTGGTGTTTGGTCTGCTATGGTTGGGACTTCTTTAAGTTTATTGA
TTCNGACTGAGTTAATAGTAATGGGTAATTTATTAGGGGATGATCAGTTATTTAATGTGAT
TGTAACTGCTCATGCTTTTGTAATAATTTTTTTATAGTAATACCTATTATAATTGGAGGGT
TTGGAAATTGATTAGTACCTTTAATACTTGGTGCNCCGGATATGGCATTTCCTCGTTTAAA
TAATTTAAGATTTTGGTTGTTACCNCCTTCATTTTTTTATTATTAAGTTCTTCAATAGTTG
AAAGNGGGGCTGGNACTGGATGGACAGTTTATCCTCCTTTATCAAGAAATTTAACTCATA
GTGGTGGGTCAGTTGATTTAACTATTTTTTCTTTACATCTTGCTGGAGTTTCTTCANTTTTA
GGAGCTTTAAATTTTATTACTACAGTAATTAATATACGTACTTTNGGTATAGTATTTGAGC
GTGTTCCTTTATTTGTTTGATCAGTAAAGATTACTGCTATTTTATTATTATTATCATTACCT
GTTTTAGCAGGTGCTATTACTATATTATTAACTGATCGAAATTTAAATACATCATTTTTTG
ACCCAGCTGGGGGTGGAGATCCAATTTTATATCAACATTTATTT

>NC_016117

TCTATATTTTTTTTTTTCTATAATTATGGGTTTTTGTGCTTTTTTCTATTCTTTTGTCATGCG
TTTAGCTTTAGTTTGACCTTTTGCTTTTATCGAATCTGGTATTATTTATTTATATTACGTCA
CTTTACATGCTGTTTACATGATTTTTTTTTTTGTTATGCCTTTTAGTATAGGTGGTTTATCA
AATTTATTAATTCCATTATGTTTTCATTTGGCTGATATGTGTTTACCTCGTATTAATAATTT
ATCTTTCTGACTTTTATTTGCTTCTTTTATTATCTCTCTTTTATCCTCTTTTCACTACTATGG
TCCAAGTTCAGGATGAACTTTATACCCTCCTTATTCTTCTTATCCTGCTAGTGCCTATTTAT
CAACTGATTTAATAATTTTCTCTTTACATTTAGCGGGTGCTAGTTCTATATTATCATCCATA
AACTTTATAGTTACTGTATTTATTTTACCCATAAATACTTCGTTTTCATTTTTTCAATATCC
TTTATTTATTGTCGCCCAAATTACTGTTTCTTTTCTTCTTTTAATATCATTACCTGTATTAGC
TGCAGCTATTACTATGTTACTTTTTGATCGTAATTTTAATACTTCTTTTTTTTCAAATTATCT
TGGTGGTGATGCTTTACTTTATCAACATTTATTT

>NC_016122

TACTCTATATTTAATTTTCGGTGCCATTGCTGGAGTAATGGGTACATGCTTCTCAGTACTA
ATTCGTATGGAATTAGCACAACCCGGCAATCAAATTCTTGGTGGAAATCATCAACTTTAT
AATGTATTAATAACAGCTCACGCTTTTTCAATGATCTCCTTCATGGTTATGCCGGCGATGA
TAGGTGGTTTTGGTAATTGGTTCGTTCCTATTCTTATAGGAAGTCCGGATATGGCATTCCC
CAGATTAAATAATATTTCATTTTGGCTTTTGCCACCGTCATTGTTACTTCTTTCAAGCTCAG
CCTTAGTAGAGGTGGGTTGCGGTTCGGGGTGGACGGTCTATCCACCCTTAAGTGGTATAA
CCAGTCATTCTGGAGGATCTGTTGATTTAGCCATTTCTAGCCTTCATTTATCAGGTGTTTCT
TCTATTTTAGGTTCTATTAATTTTATAACAACTATCTTCAATATGAGGGGCCCTGGATTGA
CTATGCATAGATTACCTCTATTTGTGTGGTCTGTTTTAGTGACAGCTTTCCTACTTTTATTA

TCCCTTCCAGTACTGGCAGGTGCAATTACCATGTTATTAACTGATAGAAATTTTAATACAA
CCTTTTTTGATCCTGCTGGTGGCGGGGATCCCATTCTATACCAGCATCTTTTT

>NC_016676

GACTTTATACTTAATTTATGGCCTATGGTCAGCGATACTAGGATCTTCTTTTAGTTTTATTA
TTCGTATTGAATTAAGAAGCCCTGGTACATTAATTGGAAATGACCAGATTTATAATGTCA
TCGTTACAGCCCATGCTTTTATTATAATTTTTTTTATAGTGATACCAATTATAATTGGMGG
ATTTGGTAACTGATTAATCCCACTAATATTAAATGCTCCGGATATAGCCTTTCCTCGAATA
AACAATATAAGATTTTGATTACTCCCTCCTTCCTTATCTTTTCTCCTGGTTTCCTCTTTTATT
GATATGGGAAGTGGAACTGGATGAACAATTTATCCTCCTCTATCTTCAAATCAAGCACAT
AGAGGAAATTCTATAGATTTTACTATTTTTTCTTTACACTTAGCAGGAATATCTTCTATTCT
AGGGGCAATTAATTTTATCACCACCATTTTAAATATACGTCCTTCCTACCTACAGATAGAC
CGAGTTCCTATATTTGTTTGATCTGTCTTAATTACAGCAATTCTTTTACTTCTTTCTCTCCCT
GTTTTAGCGGGGGCTATTACAATACTTTTAACTGATCGAAACTTTAATACAAGTTTTTTTG
ACCCCACAGGAGGAGGAGACCCTATTTTATTTCAACACTTATTT

>NC_016754

GATTATATATTTTATTTTTTCATTATGGAGAGGTTTAATAGGGGTTGTAATAAGGTTAATT
ATTCGTTTAGAGCTGGGGAGGGTCGGGTCGTTACTAATGGATGACCACTTGTATAATGTG
CTAGTTGGATCACATGCTGTTATAATGATTTTCTTTTTAGCAATACCGGTGATAATAGGAG
GGTTTGGTAATTGGCTGATTCCTATTATGTTGGGAGTAGGGGATATAGCTTTTCCACGGTT
AAATAATTTTAGGTTTTGACTACTTCCAGTTTCTATGGCATTTATAATAATATCCCTGATG
ATAGGGGGTTCTGGCTCAGGGTGGACTATTTACCCTCCTTTAAGTAGGTGGGTCTATAGA
AGTGGAGCATCTGTGGATTTGATGGTTTTAGCCTTACATGTGGCGGGCATTTCATCTGTAT
TAGCTTCTGTGAATCTGATGGCTACTGTATGGGGTGCATGTAGTGATAGTGGGGTGAGTT
TTGAGAAGTTGACTTTATTTACATGGTCTTTGATGGTCACTGCAGGGCTGATTATTTTAAC
TGTTCCTGTTTTAGCAGCAGCTTTAACCATATTGTTGTTGGACCGAAATTTAATAACTACT
TTCTTTGATCTCTCTGGAGGGGGCAGGGTGATTATGTATCAGCATTTGTTT

>NC_016951

CACCTTATATTTAATTTTTGGGGCCTGAGCAGGTATAGTAGGCACAGCATTAAGTTTATTA
ATCCGCGCAGAATTAAGCCAACCGGGGGCCCTCCTAGGGGACGACCAAATCTATAATGTT
ATCGTTACAGCCCATGCCTTTATTATAATTTTCTTCATGGTTATACCCATTATAATCGGCG
GCTTTGGAAACTGACTTGTGCCACTAATAATTGGGGCGCCAGATATGGCATTCCCACGTA
TAAACAATATAAGCTTCTGGCTTCTACCACCATCCCTACTTTTACTTCTAGCCTCCTCAGG
AATTGAAGCAGGCGCAGGCACAGGCTGAACTGTGTACCCGCCATTAGCTGGAAACCTGG
CCCACGCTGGTGCCTCTGTAGATTTAACTATCTTTTCCCTTCACCTAGCAGGTGTGTCATC
AATTTTAGGGGCCATCAACTTTATCACCACAGCAATTAACATAAAATCTCCAGCTATATC

ACAGTACCAAACACCTTTATTTGTATGATCCGTACTTATCACAGCCGTCCTATTACTACTC
TCACTACCAGTACTCGCCGCAGGTATTACTATACTACTCACAGACCGAAACCTAAATACA
ACCTTCTTCGACCCTTCAGGGGGAGGGGACCCAATTTTATACCAACACCTGTTT

>NC_017749

AACATTATATTTTATTTTTGGTATTTGATCAGGTTTATTAGGTACTTCATTAAGTTTAATAA
TTCGAAGAGAATTAGGAAAACCAGGTACTCTATTAAATGATGATCAATTATATAATGTTG
TAGTAACCGCCCACGGTTTTATCATAATTTTCTTTTTAGTTATACCTATTATAATTGGAGGT
TTTGGTAATTGATTAGTTCCCTTAATATTAGGGGCACCAGACATAGCCTTCCCTCGAATAA
ATAATATAAGTTTTTGGTTATTACCTCCATCTTTAACTCTTTTATTATCATCCTCAGCTGTA
GAAAGAGGTGCTGGAACTGGATGAACAGTATATCCTCCCTTATCTAGTAATCTATCTCAT
GCTGGCCCATCTGTAGATTTAGCTATTTTTTCTTTACATTTAGCTGGTGTTTCCTCAATCTT
AGGTGCTATTAATTTTATTACAACTATTTTAAATATACGGTGAGAGGGTTTACAAATAGA
ACGACTTCCTTTATTTGTTTGATCCGTATTTATTACAGCTATTTTACTACTATTATCCTTAC
CAGTTTTAGCTGGAGCCATTACTATATTATTAACCGATCGAAATTTTAATACAACATTTTT
TGATCCTAGAGGAGGAGGTGACCCTATTTTATATCAACATTTATTT

>NC_017755

TACTCCATATCGTATTTTTGGTGCCATTGCTGGAGTAATGGGTACATGCTTTTCCGTACTA
ATTCGTATGGAATTAGCACAACCTGGCAATCAAATTCCTGGTGGGAATCATCAACTTTAT
AATGTGTTAATTACAGCTCATGCTTTTCCAATGATCTTTTTTATGGTTATGCCAGCGATGA
TAGGTGGATTTGGTAATTGGTTCGTTCCGATTCTTATAGGTGCACCTGACATGGCATTTCC
ACGATTAAATAATATTCCCTTTTGGTTGTTGCCACCTTCGCTGTTGCTTTTATTAAGCTCAG
CCTTGGTAGAGGTGGGTAGCGGCACTGGGTGGACGGTCTATCCGCCCTTAAGCGGTATAA
CCAGCCATTCCGGAGGATCCGTTGATTCAGCAATTTTTAGTCCTCATTTATCAGGTGTTTC
ATCTATTTTAGGTTCTATAAATTTTATTACTACTATTTTCAACATGCGCGGCCCTGGAATG
ACCATGCATAGATTACCTCTATTTGTTTGGTCTGTTTCAGTCACCGCATTCTCACCTTCATT
ATCCCTTCCTGTATTGGCAGGTGCAATTACCATGTTATTAACCGATAGAAATTTTAATACA
ACCTTTTTTGATCCTGCTGGAGGGGGAGATCCAATTTTATACCAGCATCTTTTT

>NC_017837

TACTTTATATTTAATTTTTGGCGCTTTCTCCGGTATCTTAGGTGCTTGCGCGTCTATATTGA
TCCGAATGGAACTAGCACAACCAGGTAATCAACTATTATTAGGCAATCATCAAGTGTATA
ACGTACTAGTTACAGAGCACGCATTTTTGATGATTTTCTTTATGGTTATGCCCGTCCTAAT
TGGAGGATTTGGAAACTGATTCGTACCTATTATGATAGGTGCTCCAGATATGGCCTTTCCT
AGATTAAATAATATAAGTTTTTGACTACTACCTCCATCATTGTGTCTTCTTTTAGGATCTG
CGATGGTAGAAGTAGGCGCTGGCACAGGCTGAACTTTATATCCGCCTTTGAGCTCTATTC
AGAGCCATTCAGGCGGTGCTGTTGATCTTGCTATTTTTAGTTTACACTTGTCAGGTGCTTC

TTCTATATTAGGAGCTATTAATTTCATTACGACGATATTTAATATGCGCAATCCAGGACAA
AGTATGTATCGAATACCGCTATTTGTTTGATCTATCCTCATTACTGCGTTTCTTTTACTACT
AGCAGTACCTGTCTTGGCAGGGGCCATCACAATGCTGTTAACAGATAGAAACTTTAATAC
AACATTTTTTGACCCTTCAGGTGGTGGCGATCCTGTATTGTATCAGCATTTATTC

>NC_017877

GACGTTATATTTTTTATTTGGTATTTGGTCTGGGCTTGTTGGTACTGGGTTAAGAATGTTG
ATTCGGGCTGAGTTGGGACAGCCTGGTGCTTTGTTAGGGGATGATCATTTGTATAATGTA
ATTGTGACAGCGCATGCTTTTGTTATGATTTTTTTCTTGTTATGCCAGTTATGATTGGTGG
TTTTGGGAATTGGTTGGTGCCTTTAATGTTAGGGGCACCTGATATGGCTTTTCCTCGTATG
AATAATATAAGTTTTTGGTTGTTACCGCCGTCTTTGTTGTTGTTGTTGAGTTCTGCTGCTGT
AGAGGGGGGGGTAGGTACAGGTTGAACTGTTTATCCACCTTTAGCTGGAAATACTGCGCA
TGCTGGTGGGTCTGTTGATCTAGCTATTTTTTCTTTACATTTGGCAGGGGTTTCTTCTATTT
TAGGTGCTATTAATTTTATTACGACTGTTTTAAATATGCGATGGCGTGGTTTACAGTTTGA
GCGTCTTCCTTTATTTGTTTGGTCTGTAAAAATTACGGCTGTTTTGTTGTTGTTGTCTTTAC
CTGTTTTAGCTGGAGCTATTACTATATTGTTGACTGATCGAAATTTTAATACTTCTTTTTTT
GATCCTGCTGGAGGTGGGGATCCTATTTTATATCAGCATCTTTTT

>NC_018056

AGTATTATACTTTATCTTTGCAATTTTTTGTGGTATGGCTGGTACAGCAATGTCATTAATT
ATTAGATTAGAATTAGCTGCTCCAGGTGTACAATATTTAGGTGGTAATAATCAATTATTTA
ATGTATTAGTAGTAGGACATGCAGTATTAATGATTTTCTTCTTAGTAATGCCTGCATTAAT
TGGAGGGTTTGGTAACTATTTATTACCTTTAATGATTGGTGCATCTGATATGTCATTTGCA
AGATTAAATAATATTAGTTTTTGATTATTACCTCCTGCATTAGTATGTTTAGTAACTTCTAC
ATTAGTTGAATCAGGAGCAGGTACAGGATGAACTGTATATCCTCCTTTATCATCTATTCAA
GCACATTCAGGACCTAGTGTAGATTTAGCAATTTTTGCATTACATTTAACATCAATTTCAT
CATTATTAGGTGCTATTAATTTCATTGTAACAACATTAAATATGAGAACAAATGGTATGA
CAATGCATAAATTACCATTATTTGTATGAGCTATTTTTATTACAGCTTTCTTATTATTATTA
TCATTACCTGTATTATCAGCAGGTGTTACAATGTTATTATTAGATAGAAATTTTAATACTT
CATTCTTTGAAGTAGCTGGTGGTGGTGACCCTGTGTTATATCAGCATTTATTC

>NC_018822

CACCCTCTACATAGTCTTCGGTGCCTGGGCCGGGATGGTTGGGACTGCCCTAAGCCTCCTC
ATCCGGGCCGAATTGAGTCAGCCCGGAGCCCTGCTCGGGGATGACCAAATCTATAATGTT
CTTGTTACCGCCCACGCTTTCGTTATAATCTTTTTTATAGTGATGCCTATCATAATCGGCG
GCTTTGGAAACTGACTTATCCCCCTCATAATTGGGGCCCCAGACATAGCCTTCCCGCGAA
TAAATAACATAAGTTTTTGACTTCTCCCCCCCTCATTCTTACTTCTACTGGCAGGCTCCGG
GGTAGAAGCTGGGGCCGGTACCGGTTGGACCGTATATCCCCCCCTTGCTAGTAATCTAGC

CCATGCCGGGGCTTCAGTAGACTTAACAATTTTTTCTCTCCACCTAGCTGGGGTTTCTTCA
ATTCTCGGTTCAATCAATTTTATCACAACAATTATTAATATAAAACCCCCTGCAGCCTCTC
AATACCAAACCCCCCTATTTATCTGATCTGTAATAATTACAACAGTTCTTTTGCTTCTCTCC
CTCCCAGTTCTTGCTGCCGGCATCACCATACTTCTAACAGATCGAAATCTAAACACAACG
TTCTTTGACCCAGCAGGTGGAGGAGACCCCATTTTATACCAACATCTTTTC

>NC_019623

AACAATATATCTAATTTTTGGAATNGAGCCGCTATAGTAGGGACTGCCCTTAGAATTCT
AATTCGAGCTGAACTTGGACAACCAGGATCATTAATTGGGGATGATCAAATTTATAATGT
AATCGTAACAGCCCACGCATTTGTAATAATTTTCTTTATAGTAATACCAGTAATAATTGGA
GGTTTTGGAAACTGATTAACTCCCTTAATATTAGGAGCTCCAGACATAGCATTTCCCCGCT
TAAATAACATAAGATTTTGATTACTCCCCCCTTCTTTCCTTCTTCTTCTCAGATCAGCCGCA
GTTGAAAGAGGGGCAGGAACAGGATGAACCGTTTATCCACCCTTAGCCTCAAATATTGCA
CATGCAGGAAGATCAGTTGATTTAACAATCTTTTCTCTCCACTTAGCAGGAATTTCTTCAA
TTTTAGGAGCCATTAATTTTATCACAACAATCATTAATATACGAACATCAGGAATGGTATT
AGAACGTATACCACTATTTGTATGATCTGTCAAAATTACAGCAATTCTATTACTTCTTTCT
TTACCTGTTTTAGCAGGTGCTATTACAATACTCCTAACAGATCGAAATTTTAATACATCCT
TCTTTGACCCAGCAGGTGGAGGTGATCCAATCCTATACCAACATTTATTT

>NC_019807

TATACTTTATTTTTTGATCAGGGTGTGAGGTGGTTTATTAGGGTTTAGGTTGAGTGGGTTG
ATTCGACTTGAGTTAAGGGTTGGTGGTTGTTGATTAGGGAGAGAAGCTCTTTATAACATA
ATTGTGACAAGTCACGCTATTTTAATAGTGTTCTTTTTAGTGATGCCTTTATTTATAGGGG
GTTTAGGGAATTTGTTAACACCTTTGATAATAGGGGTAAGGGATATAGCTTTTCCTCGCTT
AAACAATATCAGGGTTCTAATAGTTTATTTATCTTTAGGTTTGTATATGTTTAGGATGCTC
AAAGAGGGCCTCAGACCAGGGTGGACTTTTTATCCTCCCTTATCTTCTTCAGTGTTCAGTC
CTGATGTTAGGGTGGATATCAGAATTTTCAGGTTGCATTTATTAGGGGTTTCCTCAATTTT
GAACTCCATTAATATTTTAAGTACTGTTTATATAGGGTTGAGGAGAAATAGAATAAGGTT
AGAGATAACTCCGTTGTTTGTATGGTCTTTATCTGTTACAAGGGTTTTAGTTGTAATCACT
ATTCCAGTTTTGGCCGCGGCTTTAGCGATATTATTGTTAGATCGTAGGTTAAATTCTTCGT
TTTTTGATCCTGCAGGGGCAGGGGGTTTAGTGCTTTATCAACATTTGTTT

>NC_020335

CACTATATATTTAATTTTTGGAACATGAGCTGGTATAATAGGATTATCTATAAGAATTCTT
ATTCGAATAGAATTAGGACAACCTGGAACTTTAATCGGTAATGATCAAATCTATAACGTA
ATTGTTACAGCTCATGCATTTATTATAATTTTTTTTATAGTAATACCTATAATAATTGGAG
GATTCGGAAATTGGTTAGTACCAATTATATTAGGTGCTCCAGATATAGCATTTCCTCGAAT
AAATAATATAAGATTTTGATTACTACCTCCATCATTATTTTTATTAATTAGATCTTCCCTAA

TTGAAAGAGGAGCTGGGACAGGTTGAACTGTTTATCCCCCTCTTTCATCTAACTTATCTCA
CTATGGCCCTTCTGTTGATATAGCAATTTTCTCATTACATTTAGCAGGAGCTTCATCAATT
TTAGGAGCAATTAATTTTATTACAACTATTATTAACATACGTTCTATCGGAATAACATTAG
AACGAATACCTTTATTTGTTTGATCAGTATTAATTACTGCAATTTTACTTCTTTTATCTTTA
CCTGTTTTAGCAGGTGCTATTACTATATTACTAACAGATCGAAATTTTAATACATCTTTTTT
CGACCCTTCAGGGGGAGGAGATCCAATTTTATATCAACATTTATTT

>NC_020654

CACTCTTTATCTAATCTTTGGTGCCTGAGCCGGAATAGTAGGAACCGCATTAAGCTTATTA
ATTCGCGCAGAACTTGGTCAACCTGGAGCTCTAATAGGAGATGACCAGATTTATAATGTA
ATCGTCACTGCTCATGCATTTGTAATGATTTTCTTTATAGTAATACCAATTATAATCGGCG
GATTTGGCAACTGACTTGTACCGCTAATAATTGGAGCCCCTGATATAGCCTTTCCTCGCAT
AAATAATATAAGCTTCTGACTCCTTCCACCCTCACTTCTTCTTCTACTTACCTCCTCCGCGG
TAGAAGCCGGAGTGGGGACCGGATGAACCGTATACCCCCCACTAGCCGGAAACCTGGCA
CATGCAGGAGCATCAGTTGACTTAGCTATCTTCTCACTCCATTTAGCCGGTATTTCTTCAA
TTCTTGGAGCCATTAACTTTATTACTACAATCGTTAATATGAAACCACCATCTACTTCACA
ATATCAAACCCCCCTATTTGTGTGATCAGTCCTAGTTACCGCAGTACTCCTACTATTATCC
CTCCCAGTACTAGCCGCAGGTATTACAATGCTCCTTACAGACCGGAACCTCAATACTACT
TTCTTTGACCCCGCAGGCGGGGGAGACCCGATCCTCTATCAACACTTATTC

>NC_021113

TCTTTATCTAATCTTCCGATTAGCTTCACGACTTATCCGTACCTCCCTCAGCTACATCGTAA
GGCTAGAGGCCTCGTCCAGAAGCCGAGACATCTATTTTGACCGTTCCATCTAGAATACTG
CTATCACCAACCATCGGCTAATCATGATCTTCTTCCTAATTATGCCAGTCATGATCCGAGG
GTTCCGTAACTGACTCATCCCTCTCTATTTAATGGCACCAGATATGGCCTTTCCTAGGCTA
AACAATGTCAGTTTCTGGCTCCTACCCCCATCATTCCTGTGTCTTATTAACGCCTTTCTCAT
TGAAGATCGAGTACGAACTCGATGAACTCTCTATCCACCTCTCTCAGATCTTCTACGACAC
AGTCGTAAAAACGTAGACTTAGCCATTTTCAGTCTTCATCTAGCACGTCTATCTTCCATTC
TACGGGCCATCAATTTCATCACTACAATCACAAACATGAGGCGATGAAACTCTCCTTCTTT
AAGTCTAAACCTATTCTGTTGATCCATGCTTGTAACAGTTGTTCTCCTACTCCTATCTTTAC
CCGTTCTAGCCCGAGCCATCACCATGCTACTTTTTGACAGAAATCTAAACACTAGTTTCTT
TGAGCCATCCCGAGGTCGAGATCCTATCCTTTTCCAGCACCTCTTC

>NC_021126

GATTCTCTACATATTGTTTGGAACTCTTGCTGGTATTACAGCAACGACCATTTCAATTGTA
ATGAGAATGGAACTTGCTGCTCCAGGAGATCAGTTACTTGGTGGTAATTATCAACTTTAC
AATGTTTTAATTACTGCGCATGGATTATTAATGCTTTTCTGGGTTGTAATGCCTATTGTTTT
AGGTGGCTTTGGTAATTTCTTTGTACCTTTATTAATTGGTGCACCTGATATGGCATTCCCT

AGATTAAATAATATTAGTTTTTGGTTACTTCCACCTTCACTTGTTTTACTCGTGCTTTCAGC
TCTTGTTGAAACAGGGGCAGGTACAGGATGGACAGCATATCCTCCATTATCTGGAATACA
ATCACATTCTGGAGCTTCAGTCGATCTTGCTATTTTTAGCTTGCACCTCTCAGGTACATCG
TCTATTTTGGCTTCTATTAACTTTATTGCTACGATCTTTAATATGAGAGCTCCTGGTATGAC
AATGCATAAAATGCCTTTATATGTATGGTCAATCCTTGTAACTTCTTTCTTGCTTGTACTAT
CTCTTCCTGTGTTTGCAGGTGCAATTACAATGCTTCTTACTGATCGTAATTTTAATACTAC
ATTCTTTGATCCTGCTGGAGGTGGTGATCCAATCTTGTTCCAACACTTATTT

>NC_021152

GACTCTCTATTTCATCTTCGGTGCCATTGCTGGAGTGATGGGCACATGCTTCTCAGTACTG
ATTCGTATGGAATTAGCACGACCCGGCGATCAAATTCTTGGTGGGAATCATCAACTTTAT
AATGTTTTAATAACGGCTCACGCTCCTTTAATGATCTTTTTTATGGTTATGCCGGCGATGA
TAGGTGGATCTGGTAATTGGTCTGTTCCGATTCTGATAGGTGCACCTGACATGGCATTTCC
ACGATCAAATAATATTTCATTCTGGTTGTTGCCACCAAGTCTCTTGCTCCTATTAAGCCCA
GCCTTAGTAGAAGTGGGTAGCGGCACTGGGTGGACGGTCTATCCGCCCTTAAGTGGTATT
ACCAGCCATTCTGGAGGAGCAGTTGATTCAGCAATTTCTAGTCCTCATCTATCAGGTGTTT
CATCCATTTTAGGTTCTATCAATTTTATAACAACTATCTCCAACATGCGTGGACCTGGAAT
GACTATGCATAGATCACCCCTATTTGTGTGGTCCGTTCCAGTGACAGCATTCCCACTTTTA
TTATCACTTCCGGTACTGGCAGGGGCAATTACCATGTTATTAACCGATCGAAACTTTAATA
CAACCTTTTCTGATCCCGCTGGAGGGGGAGACCCCATATTATACCAGCATCTCTTT

>NC_021932

AACAATATATTTAATTTTTGGAGCCTGAGCAGCCATAGTAGGGACAGCTTTAAGAATTCT
AATTCGATTAGAACTAGGACAACCAGGAAGTCTAATTGAAGATGATCAAATTTACAATGT
AATTGTAACCGCCCACGCCTTTGTTATAATTTTCTTCATAGTTATACCCATCATAATTGGA
GGATTTGGAAATTGATTAGTGCCTCTAATATTAGGAGCACCAGATATAGCCTTTCCCCGA
CTCAACAACATAAGATTTTGATTACTTCCACCAGCTTTCTTCCTTCTTTTAGCCTCCTCTGC
AGTAGACAAAGGAGTAGGTACAGGATGAACAGTATACCCCCCTCTTGCCAGAAATATTG
CCCACTCAGGGCCAGCCGTAGACATAGCAATTTTTTCCCTACACCTAGCAGGAGCTTCTTC
AATCTTAGGAGCAATTAATTTTATTACAACAATTATTAATATACGGTCCAGAGGAATATT
ATTTGAACGAATACCTTTATTTGTATGGGCTGTTAAACTAACAGCAATTCTTCTACTTTTA
TCTTTACCAGTACTAGCAGGGGCAATTACTATGCTCCTAACAGATCGTAACTTTAATACTT
CATTTTTTGACCCCGCGGGAGGAGGGGACCCTATCCTATACCAACACTTATTC

>NC_022146

AACCTTATACCTACTATTTGGCGCATGATCTGGCCTAATTGGGGCCTGCCTAAGCATCCTA
ATACGCATAGAACTAACCCAGCCAGGCTCACTACTAGGTAGTGACCAGATCTTTAATGTC
CTAGTTACAGCCCACGCATTCATCATAATTTTCTTCATAGTAATACCAATTATAATCGGGG

GCTTCGGTAACTGACTAATCCCCCTAATAATTGGAGCCCCAGACATAGCCTTTCCACGTAT
AAACAATATAAGCTTTTGACTACTACCACCAGCACTCCTTCTTCTACTATCCTCTTCCTAC
GTTGAAGCTGGGGCCGGCACAGGATGAACCATCTACCCACCACTATCAGGAAACCTAGT
ACACTCAGGCCCATCAGTAGACCTAGCAATCTTCTCCCTCCATCTAGCAGGCGCCTCCTCC
ATCCTGGGGGCAATCAACTTCATTACAACATGTATCAACATAAAACCTAAATCTATACCA
ATATTCAACATCCCCCTGTTCGTTTGATCAGTACTTATCACTGCTATTATACTACTTTTAGC
TCTACCTGTGCTGGCAGCAGCAATTACCATACTACTAACTGACCGAAACCTAAACACCTC
ATTCTTTGACCCTTGCGGGGGAGGAGACCCAGTACTATTCCAACACCTGTTT

>NC_022256

AACCCTTTATTTAATTTTTGGAGCTATTGCTGGAGTTGCTGGAACAACTCTTTCTGTTCTA
ATTCGATTAGAATTAGCTCAACCAGGAAATCAGTTTTTATCTGGAAACAATCAGTTATAC
AACGTTATTGTAACCGGACACGCATTTGTTATGATTTTTTTCTTCGTAATGCCTGTTCTTAT
TGGCGGTTTCGGTAACTGGTTTGTACCTTTAATGATTGGTGCTCCTGACATGGCTTTTCCA
CGAATGAATAACATTAGTTTTTGGTTATTACCTCCTTCTCTTATCCTTTTATTGGCTTCAAC
TTTTGTTGAGGCTGGAGCAGGAACTGGTTGGACTGTATATCCTCCTTTAAGTGGTGCGCA
AGCTCACTCGGGACCTTCTGTGGATTTAGCTATATTTAGTCTTCACTTATCAGGTGCTGCT
TCAATTTTAGGTGCTATTAATTTTATTACTACTATTTTTAATATGCGTGCACCTGGTATGAG
TATGCATAGATTACCTTTATTTGTGTGGTCTGTTTTAATTACTGCTTTCTTACTTTTATTGTC
GCTTCCTGTTTTTGCTGGAGCGATTACTATGCTGTTAACTGATAGAAACTTTAATACTACT
TTTTACGATCCTGCAGGTGGTGGTGATCCTGTATTATACCAGCATTTATTC

>NC_022674

AACAATGTACTTAATCCTTGGGGCCTGATCTGCCATATTAGGAACTGCCCTTAGAATGCT
AATCCGGGCCGAACTAGGTCAACCGGGAAGTCTCATTGGAGATGACCAAATCTACAACG
TAATTGTAACAGCCCACGCCTTCATCATAATTTTCTTTATGGTAATGCCCATTATAATTGG
AGGGTTCGGAAATTGACTTGTTCCACTAATGTTAGGTGCCCCCGATATGGCCTTCCCACG
GCTAAACAACATAAGATTCTGACTCCTTCCCCCCTCTCTAACCCTGCTCCTAGCGGGGAGT
GCTGTGGAAAATGGAGCCGGAACTGGATGAACAGTCTATCCTCCCCTCGCTTCCAATATT
GCCCATGCCGGAGCTTCCGTAGACCTAACCATTTTCTCCCTTCATTTAGCTGGGGCATCCT
CAATCCTAGGAGCCATTAACTTCATTACCACTGTTATTAATATACGAACACAGGGAATAA
CCATAGAACGAATGCCACTATTTGTATGAGCGGTCTTCATTACAGCCTTCCTACTACTATT
ATCACTTCCCGTACTGGCCGGTGCCATTACAATACTGCTCACAGACCGTAATTTAAACACT
TCATTCTTTGATCCAGCAGGAGGGGGAGACCCCATTCTATATCAACACCTATTC

>NC_022796

GACTCTATATTTCATCTTCGGTGCCATTGCTGGAGTGATGGGCACATGCTTCTCAGTACTG
ATTCGTATGGAATTAGCACGACCCGGCGATCAAATTCTTGGTGGAAATCATCAACTTTAT

AATGTTTTAATAACGGCTCACGCTTTTTTAATGATCTTTTTTATGGTTATGCCGGCGATGA
TAGGTGGCTTTGGTAATTGGTTTGTTCCGATTCTGATAGGTGCACCTGACATGGCATTTCC
GCGATTAAATAATATTTCATTCTGGTTGTTGCCACCAAGTCTCTTGCTCCTATTAAGCTCA
GCCTTAGTAGAAGTGGGTAGCGGCACTGGGTGGACGGTCTATCCGCCCTTAAGCGGTATT
ACCAGCCATTCTGGAGGAGCAGTAGATTTAGCAATTTTTAGTCTTCATCTATCTGGTGTTT
CGTCCATTTTAGGTTCTATCAATTTTATAACAACTATCTTCAACATGCGTGGGCCTGGAAT
GACTATGCATAGATTACCCCTTTTTGTGTGGTCCGTTCTAGTGACTGCATTCCTACTTTTAT
TATCACTTCCGGTACTGGCAGGGGCAATTACCATGTTATTAACCGATCGAAACTTTAATA
CAACCTTTTCTGATCCCGCTGGAGGGGGAGACCCCATATTATACCAGCATCTCTTT

>NC_022860

TACCCTTTATTTAATTTTCGGAGCTATCGCCGGGGTTATGGGTACTTGTATGTCAGTACTA
ATTCGTATGGAATTAGCACAACCAGGTAATCAAATTCTTGGTGGAAATCATCAGTTATAC
AATGTGTTAATTACAGCACATGGCTTTTTAATGATATTTTTCATGGTTATGCCTGCTTTAAT
CGGCGGGTTCGGAAATTGGTTCGTGCCTATTCTAATAGGTGCACCGGATATGGCATTCCC
ACGTTTGAATAATATCAGTTTTTGGCTACTACCTCCTTCATTATTACTTCTTCTAAGTTCTG
CTTTAGTTGAAGTAGGTGCTGGTACTGGATGGACGGTCTATCCTCCTTTAAGTGCCATAAC
CAGCCATTCAGGTGGTGCTGTAGATTTAGCTATATTTAGCCTTCATCTATCAGGAATTTCT
TCAATTTTAGGTTCTATTAACTTTATCACAACTATTTTTAACATGCGTGGTCCCGGAATGA
CTATGCATAGAATACCACTCTTTGTATGGTCTGTACTAGTTACTGCATTCTTGTTGTTATTA
TCACTGCCAGTATTTGCTGGTGCAATTACTATGCTTTTAACAGATCGAAATTTCAATACTA
CTTTCTTTGATCCTGCAGGCGGAGGAGATCCGCTTTTATATCAACATCTTTTC

>NC_023337

GACTCTATATTTCATCTTTGGTGCCATTGCTGGAGTGATGGGCACATGCTTCTCAGTACTG
ATTCGTATGGAATTAGCACGACCCGGCGATCAAATTCTTGGTGGGAATCATCAACTTTAT
AATGTTTTAATAACGGCTCACGCTTTTTTAATGATCTTTTTTATGGTTATGCCGGCGATGA
TAGGTGGATCTGGTAATTGGTCTGTTCCGATTCTGATAGGTGCGCCTGACATGGCATTTCC
ACGATTAAATAATATTTCATTCTGGTTGTTGCCACCAAGTCTCTTGCTCCTATTAAGCCCA
GCCTTAGTAGAAGTGGGTAGTGGCACTGGGTGGACGGTCTATCCGCCCTTAAGTGGTATT
ACCAGCCATTCTGGAGGAGCAGTTGATTCAGCAATTTCTAGTCCTCATCTATCTGGTATTT
CATCCATTTTAGGTTCTATCAATTTTATAACAACTATCTCCAACATGCGTGGACCTGGAAT
GACTATGCATAGATCACCCCTATTTGTGTGGTCCGTTCTAGTGACAGCATTCCCACTTTTA
TTATCACTTCCGGTACTGGCAGGGGCAATTACCATGTTATTAACCGATCGAAACTTTAATA
CAACCTTTTCTGATCCCGCTGGAGGGGGAGACCCCATATTATACCAGCATCTCTTT

>NC_023355

GACTCTTTACCTCATCTTTGCAGCCTTCTCTGGAGTTCTTGGAACTTGCTTCTCTATTCTTA
TTCGAATGGAACTTGCAGCTCCAGGAAACCAAGTTTTGAGCGGAAACCATCAGGTGTATA
ACGTGATTGTTACCGCACACGCTTTCCTCATGATTTTCTTCATGGTTATGCCTGCTCTTATC
GGAGGGTTTGGAAACTGGTTGGTTCCTCTTATGATTGGAGCTCCAGATATGGCGTTCCCA
CGATTGAACAACATTTCATTCTGGCTTCTTCCGCCAAGTTTGATCCTTCTCCTTACTTCTGC
TCTTGTAGAGGGAGGGGTAGGAACCGGATGGACTGTCTATCCTCCTCTTTCAACTCATGC
TTTCCATAGTGGGGGAGCTGTTGATCTTGCGATTTTCTCTCTTCACGTGAGTGGTATGAGT
TCGATTCTTGGAGCAATCAACTTCATTTCAACTATCTTTAACATGCGAGGACCAGGAATG
GAGATGCATCGAATGCCTCTTTTTGTTTGGTCTGTGTTGATTACTGCCTTCCTCCTTCTTCT
TAGCCTTCCTGTCCTTGCAGGAGCGATCACTATGTTGTTGACTGACCGAAACTTCAACACA
AGTTTCTTTGATCCAGCTGGTGGAGGAGACCCTATCCTCTATCAGCACCTCTTC

>NC_023504

TACTTTATATTTTTTGTTTGGTTTGTGGTCTGGTATGGTTGGTACTAGTTTGTCTTTGGTAA
TTCGTTTGGAGTTGGCTAAACCTGGTCTTTTTTTGGGTAATGGTCAGTTATATAATTCTATT
ATTACTGCTCATGCTATTTTGATGATTTTTTTATGGTTATGCCTACTATGATTGGTGGTTT
TGGTAATTGGATGTTACCTTTGATATTGGGGGCTCCTGATATGAGTTTTCCTCGTTTAAAT
AATTTAAGTTTTTGGTTGTTGCCTACCGCTATGTTTTTAATTTTGGATGCTTGTTTTGTTGA
TATGGGTTGTGGAACTAGTTGAACTGTTTATCCTCCTTTGAGAACTATAGGTCATCCTGGT
AGAAGGGTTGATTTGGCTATTTTTAGTTTGCATTGTGCTGGTATTAGTTCTATTTTGGGTG
CTATTAATTTTATGACTACTACTAAGAATCTTCGTAGGAGTTCTATTTCTTTGGAACATAT
GAGGTTGTTTGTTTGGACTGTTTTTGTGACTGTTTTTTTGTTGGTTTTATCTTTACCTGTCTT
AGCTGGGGCTATTACTATGTTGTTGACTGATCGTAATCTTAATACTTCTTTTTTTGATCCTA
GGACTGGGGGTAACCCTTTAATTTATCAGCATTTGTTT

>NC_023545

TATTTTATACCTTTTTTTTGGTGTGTTCAGTGGCTTAATTGCAACCATTATGTCCGTTTTAA
TGCGAATTGAATTGGGTAACCCTGGTGATCAGGTGTTTATGGGTAACTATCAATTATACA
ACGTTATTATTACTGCTCATGGTTTATTGATGTTGTTTTGGACTATTATGCCTATTTTAGTA
GGAGGTTTTGGTAACTTTTTTATTCCTATTATGATTGGTGCTCCTGATATGTCCTTTCCTCG
AATGAATAACTTGAGTTTTTGGATGTTACCACCATCTTTATTATTATTGTTATCTTCTGCTT
TAGTTGAAGTTGGAGCTGGTACTGGTTGGACTGCGTATCCTCCCTTATCTAGCGTAACAGC
TCATTCTGGTCCATCTGTTGATTTGGCTATCTTTGCATTACATTTAAATGGTATGTCTTCTA
TTTTGGGTTCTATCAATTTACTTGCTACTATGTTTAATATGCGTGCTCCTAATATGCCAATT
CACAAAATGCCATTATATTGTTGGTCTGTTGTTGTAACTGCTTTCTTATTGGTATTTTCCTT
ACCGGTGTTTGCAGGTGGTATTACTATGTTATTAACTGATCGTAATTTCAACACATCTTTC
TTTGATCCAGCTGGTGGAGGAGATCCAATCTTATTCCAACATTTATTT

>NC_023773

AACACTTTATTTGATTTTTGGGGCCTGAGCCGGCATGGTAGGAACAGCTATGAGTGTAAT
TATTCGTGCCGAATTAGCGCAACCCGGTTCTTTACTAAAAGATGACCAAATTTACAAAGT
AGTCGTTACCGCACACGCACTGGTCATGATCTTCTTTATGGTGATGCCAATAATGATTGGA
GGATTTGGTAATTGACTCATTCCACTAATGATTGGTGCACCAGATATGGCTTTCCCCCGTA
TGAAAAACATGAGATTTTGACTTATTCCTCCTTCTTTCATCTTACTTCTAGCTTCCGCAGG
AGTGGAAAGAGGAGCAGGAACTGGATGAACTATTTATCCTCCTCTTTCCAGTAAAATAGC
ACACGCCGGAGGATCTGTTGATCTAGCAATCTTTTCTCTTCACCTTGCCGGTGCCTCCTCC
ATTTTAGCCTCAATTAAATTTATAACAACAATAATTAATATGCGGACACCAGGGATGTCT
TTTGACCGTCTTCCGTTATTTGTTTGGTCTGTTTTTGTTACTGCATTCTTACTACTTCTTTCT
CTCCCAGTACTAGCTGGGGCAATTACTATGCTCCTCACAGATCGAAACATAAACACAACC
TTCTTTGACCCAGCAGGAGGAGGAGACCCAATTTTATTCCAACACTTATTC

>NC_023931

TTCTTTGTATTTTTTATTTGGTGTGTGATCTGGGTTAGTAGGTACTGCTTTAAGTTTGTTGA
TTCGTGCAGAGTTGGGGCAGCCTGGTGCGTTGTTAGGTGATGATCAGTTGTACAATGTTA
TTGTTACAGCGCATGCTTTTGTGATGATTTTTTTTTAGTTATGCCTGTTATGATTGGTGGA
TTTGGAAATTGGTTGGTTCCTTTAATGTTAGGGGCTCCAGATATGGCTTTTCCTCGTATAA
ATAATATGAGATTTTGGCTTCTTCCTCCTGCTTTAACTCTTCTTCTGTCTTCTGGAGCTGTT
GAAAGTGGTGCTGGTACTGGATGGACTGTTTATCCGCCTCTATCTAGAAATATTGCTCATG
CTGGTAGCTCTGTGGATTTAGCTATTTTTTCTTTGCATTTGGCGGGTGTTTCTTCTATTTTG
GGTGCTATTAATTTTATTACTACTATTATTAATATGCGTTGGTATGGAATGCAGTTTGAAC
GTTTGTCTTTATTTGTTTGACCCGTAAAAATTACTGCTATTTTGTTGTTGTTGTCTCTTCCT
GTTTTGGCAGGTGCAATTACTATATTGTTAACTGATCGGAATTTTAATACTTCTTTTTTTGA
TCCTGCTGGGGGTGGTGACCCTATTTTATATCAGCATTTGTTT

>NC_023945

AACTTTATATTTAATTTTTGGAGCTTGGTCGGCTATAGTAGGAACGGCTTTAAGAATACTT
ATTCGAGCAGAACTAGGACAACCCGGAAGCTTAATTGGAGATGATCAAATCTACAATGT
AATTGTCACAGCTCATGCCTTTATTATAATTTTTTTTATAGTTATACCTATTATAATTGGTG
GTTTTGGAAATTGATTATTACCATTAATATTAGGAGCTCCAGACATAGCTTTCCCTCGGTT
AAATAATATAAGATTTTGATTATTACCCCCAGCTCTAATATTACTTATTAGCGGATCTTTA
GTTGAAGCTGGTGCCGGAACAGGATGAACAGTTTACCCACCTTTATCAAGCAATATTGCT
CATTCAGGAGCTTCTGTAGATTTATCAATTTTTTCTTTACACTTAGCTGGAGCTTCTTCAAT
TCTTGGAGCAATTAATTTTATATCCACAGTTATTAATATACGAGCAGAGACCTTAACTTTT
GATCGACTTCCACTATTTGTTTGAAGAGTTTTTATTACAGTAATTCTTCTTCTTCTTTCATT

ACCTGTATTAGCTGGAGCAATTACTATACTTCTTACTGATCGGAATCTAAATACATCGTTT
TTTGATCCAACTGGTGGAGGAGATCCAATTTTATATCAACATTTATTT

>NC_024090

TACGTTATATTTAGTCTTTGGGATTGGGGCAGGCATGCTTGGCACGGCCTTCAGTATGTTA
ATAAGATTAGAGCTCTCGGCTCCGGGGGCTATGTTAGGAGACGATCATCTTTATAATGTA
ATTGTTACGGCACATGCTTTTATTATGATTTTTTTTTGGTTATGCCAGTGATGATAGGGG
GGTTTGGGAATTGGTTGGTTCCATTATATATTGGTGCTCCCGATATGGCCTTTCCCCGGCT
TAATAATATTAGTTTTTGGTTGTTGCCCCCTGCTTTAATATTGTTATTAGGCTCTGCTTTTG
TTGAACAAGGAGCTGGCACCGGGTGGACGGTCTATCCTCCTCTAGCGAGCATCCAGGCCC
ACTCTGGTGGGGCGGTGGATATGGCTATTTTTAGCCTTCATTTAGCTGGGGTGTCTTCGAT
TTTGGGCGCAATGAATTTTATAACAACTATATTTAACATGCGGGCTCCTGGGATAACATT
AAATAAAATGCCCTTATTTGTGTGGTCTATCTTGATTACTGCTTTTTTATTATTATTATCTT
TGCCAGTATTAGCGGGGGCCATAACCATGCTTTTAACGGATAGAAATTTTAATACCACTT
TCTTTGATCCTGCAGGGGGGGGAGACCCAATTTTATTTCAGCATTTGTTT

>NC_024103

AACTTTGTATTTTGTTTTTAGAATTTGGTCAAGATTTATTGGAACTGGGATAAGTGTTTTT
ATTCGTTTAGAGTTGTCTCAAATTAGGCAAGTTGTTGGGGATGGGCAGCTCTATAATGTTA
TTGTAACTGCTCATGCTTTTGTAATAATTTTTTTCTTTGTTATACCTATGATGATTAGGGGG
TTTGGTAACTGATTGTTACCTCTTATAGTGGGGAGTCCAGATATAGCTTTCCCACGTTTAA
ATAATATGAGTTTTTGGTTATTGCCTCCTGCACTATTTTTTCTTTTCATTAGTTCTATAATT
GAAAGTGGGGTGGGAACGGGTTGGACAGTCTATCCCCCTTTGTCAAGAAATTTAGCTCAT
TCTGGGGCTGCATTAGATTGTGCTATTTTTTCACTTCATTTGGCTAGGGTTTCTAGTATTTT
AAGGTCTTTAAATTTTATAACTACTTTGTTTAATATAAAAGTTAAGAGGTGAGGGATGTTC
TCTATATCTCTGTTTTGTTGAACTGTATTAGTTACTACTATTTTGTTATTATTATCTTTACCT
GTTTTAGCTGCAGCTATTACAATATTACTTTTCGATCGAAATTTTAATACTTCTTTTTTTGA
TCCCTCTGGGAGAAGAGATCCGGTTTTGTATCAGCACTTGTTT

>NC_024173

TACTTTATATATTTTATTCGGAATTTGAGCCGGTTTAGTTGGTACTGCCTTAAGTCTTTTAA
TCCGAGCAGAATTAGGACAACCCGGAGCTTTACTTGGAGATGATCAATTATATAATGTTA
TTGTTACAGCTCATGCCTTTGTTATAATCTTTTTTTTAGTTATACCTATAATAATTGGAGGA
TTTGGAAATTGACTGGTTCCTTTAATACTAGGTGCGCCTGACATAGCATTCCCACGATTAA
ATAACATAAGATTTTGGTTGTTGCCTCCCGCTTTATGCTTGTTATTAGGTTCCGCTGCTGTG
GAAAGAGGAGCAGGAACGGGTTGGACTGTTTATCCTCCTTTAGCAAGTAATATTGCTCAT
GCAGGAGGCTCTGTAGATTTAGCAATTTTTTCTCTTCATCTAGCAGGTGTTTCTTCAATTTT
AGGAGCTGTTAATTTTATTACAACTGTGTTTAATATACGATGAAGAGGAATACAGCTAGA

ACGTTTACCTCTATTTGTATGATCTGTAAAAATTACAGCAATTTTATTACTTCTTTCTCTAC
CAGTATTAGCAGGAGGAATCACTATACTTCTAACTGATCGGAATTTTAATACTTCCTTCTT
TGATCCTGCGGGAGGGGGAGATCCTATTTTATATCAACATTTATTT

>NC_024290

GACTCTATATTGCATTTTCGGTGCCATTGCCGGAGTAATGGGTACATGCTTCTCGGTACTA
ATTCGTATGGAATTAGCACAGCCTGGCAATCAAATTCTTGGTGGAAATCATCAACTTTAT
AATGTGTTAATAACAGCTCACGCTTTTTCAATGATTTTTTTATGGTTATGCCCGCGATGA
TAGGTGGATTTGGTAATTGGTTCGTTCCGATTCTTATAGGTGCACCTGATATGGCATTTCC
GCGATTAAATAACATTAGTTTTTGGTTGTTACCACCGTCACTGTTACTTCTTCTAAGTTCTG
CTTTGGTAGAAGTGGGCGCTGGTACCGGATGGACGGTTTATCCGCCCTTAAGTGGTATAA
CGAGTCATTCTGGAGGATCCGTTGATTTAGCCATTTTCAGTCTTCATTTATCGGGTGTTTC
CTCTATTTTAGGTTCTATCAATTTCATCACTACTATTTTTAACATGCGTGGGCCAGGAATG
ACCATGCATAGATTACCTTTATTCGTGTGGTCCGTATTAGTGACAGCATTCCCACTTTTAT
TATCTCTTCCAGTATTGGCAGGTGCAATTACCATGTTATTAACTGATAGAAACTTTAATAC
AACCTTTTTCGATCCTGCAGGAGGAGGAGATCCAATTTTATACCAGCATCTGTTT

>NC_024439

GACACTTTATCTAATTTTTGGAGCTTGAGCTGGGATAGTAGGAACAGCTTTAAGTCTTCTT
ATTCGAGCAGAATTAGGTCAACCAGGAAGACTTATTGGGGATGACCAAATTTATAATGTA
GTTGTTACAGCCCACGCATTTATTATAATTTTTTTTATAGTTATACCTATTCTAATTGGAGG
TTTCGGAAATTGATTAGTGCCATTAATATTAGGAGCACCTGATATAGCTTTTCCACGATTA
AATAATATAAGATTTTGATTATTACCCCCTGCCCTCACCCTATTATTATCAGGTGGTGCTG
TAGAAGGAGGTGCAGGAACAGGTTGAACTGTATATCCACCATTGTCAGGTGGTATTGCTC
ATGCAGGGGCATCAGTAGATTTAAGTATTTTTTCACTACATCTAGCCGGTGTTTCATCAAT
TTTAGGTGCTATTAACTTTATCACAACAATCATCAATATACGAACTAGAGGTATATCCTTG
GACCGAATTCCTTTATTCGTATGAGCTGTAGGAATTACAGCACTACTACTTCTTCTTTCTC
TTCCTGTTTTAGCAGGTGCTATCACTATATTATTAACTGATCGTAACTTAAATACTTCGTTC
TTTGATCCAGCTGGTGGTGGAGACCCAATTTTATATCAGCATCTATTT

>NC_024520

GACTCTATATTGCATTTTTGGTGCCATTGCCGGAGTAATGGGTACATGCTTCTCGGTACTA
ATTCGTATGGAATTAGCACAGCCTGGCAATCAAATTCTTGGTGGAAATCATCAACTTTAT
AATGTGTTAATAACAGCTCACGCTTTTTTAATGATCTTCTTTATGGTTATGCCTGCGATGA
TCGGTGGATTTGGTAATTGGTTCGTTCCGATTCTTATAGGTGCACCCGATATGGCATTTCC
ACGATTAAATAACATTAGTTTTTGGTTGTTACCACCGTCACTGTTACTTCTTCTAAGTTCTG
CTTTGGTAGAAGTGGGCGCTGGTACCGGATGGACGGTTTATCCGCCTTTAAGTGGTATAA
CCAGTCATTCTGGAGGATCTGTTGATTTAGCCATTTTTAGTCTTCATTTATCGGGTGTTTCC

TCTATTTTAGGTTCTATCAATTTTATCACTACTATTTTCAACATGCGTGGGCCAGGAATGA
CCATGCATAGATTACCTTTATTCGTGTGGTCCGTATTAGTGACAGCATTCCTACTTTTATT
ATCTCTTCCAGTACTGGCAGGTGCAATTACCATGTTATTAACTGATAGGAACTTTAATACA
ACCTTTTTTGATCCTGCAGGAGGGGGAGATCCAATTTTATACCAGCATCTGTTT

>NC_024521

CACTCTATATTGTATTTTCGGTGCCATTGCTGGAGTAATGGGTACATGCTTTTCAGTACTA
ATTCGTATGGAATTAGCACAACCTGGCAATCAAATTCTTGGTGGAAATCATCAACTTTAT
AATGTGTTAATAACAGCTCACGCTTTTTTAATGATCTTTTTTATGGTTATGCCTGCGATGA
TAGGTGGATTTGGTAATTGGTTTGTTCCAATTCTTATAGGTGCACCTGATATGGCATTTCC
ACGATTAAATAATATTAGTTTCTGGTTGTTACCACCGTCATTGTTACTTCTTCTAAGTTCTG
CTTTGGTAGAAGTGGGTGCTGGTACTGGATGGACGGTCTATCCGCCCTTAAGTGGTATAA
CCAGTCATTCTGGAGGATCTGTTGATTTAGCCATTTTTAGTCTTCATTTATCGGGTGTTTCC
TCTATCTTAGGTTCTATCAATTTTATCACTACTATCTTCAACATGCGTGGCCCAGGAATGA
CCATGCATAGATTACCTTTATTTGTGTGGTCTGTATTAGTGACAGCATTCTTACTTTTATTA
TCTCTTCCAGTATTGGCAGGTGCAATTACCATGTTATTAACTGATAGGAACTTTAATACAA
CCTTTTTTGATCCTGCAGGAGGAGGAGATCCAATTTTATACCAGCATCTTTTT

>NC_024596

CACCCTTTACCTGATTTTTGGTGCATGAGCAGGTATAGTTGGAACAGCCCTAAGTCTCCTA
ATTCGAGCTGAACTTGGGCAACCTGGATCACTTTTAGGGGATGATCAGATTTATAATGTA
ATTGTAACCGCCCACGCTTTTGTAATAATCTTTTTTATAGTTATACCAATTATAATTGGTG
GTTTCGGAAATTGATTAGTTCCTTTAATAATTGGTGCACCAGATATAGCCTTCCCACGAAT
AAATAACATAAGTTTCTGACTTCTTCCACCATCATTTCTTCTTCTCCTCGCCTCTGCTGGAG
TAGAAGCTGGAGCAGGTACTGGTTGAACAGTCTATCCCCCATTAGCTAGTAACCTAGCAC
ATGCTGGACCATCTGTTGACTTAGCTATTTTCTCTCTTCACTTAGCCGGTATTTCATCAATT
TTAGCTTCAATTAATTTTATTACAACAATTATTAACATAAAACCACCAGCCATTTCACAAT
ATCAAACACCATTATTTGTTTGATCTATTCTTGTAACCACTATTCTTCTTCTCCTCTCACTT
CCAGTTCTTGCAGCAGGGATTACAATACTACTTACAGATCGTAACCTCAACACTACATTCT
TTGACCCTGCAGGTGGAGGAGATCCAATTCTTTATCAACATTTATTT

>NC_024626

AACTCTGTATTTAATTTTTGCAGCTTTTTCAGGAGTATTGGGAACTATGTTCTCAGCATTA
ATTCGTATGGAGCTAGCACAACCTGGAAACCAAATTCTAGGCGGAAACCACCAATTATAT
AACGTAATTATTACAGCTCACGCATTTTTAATGGTATTCTACTTAGTAATGCCTGCTTTAA
TGGGTGGTTTTGGTAATTGGTTTGTACCAATTTTAATTGGTGCCCCAGATATGGCATTCCC
TCGTTTGAATAATATTAGTTTCTGGTTATTACCTCCATCTCTTCTATTACTTTTAAGCTCTG
CTCTTGTTGAAGTTGGTGCGGGTACAGGGTGGACAGTGTATCCTCCATTAGCAGGTATTG

CTAGCCACTCTGGTGGATCTATTGATCTTGCAATTTTCAGTTTACACCTTGCAGGTGCAAG
TAGTATTATTGGTTCTATTAATCTGATTACTACAATTTTTAATATGAGAGCTCCTGGTATG
AGTATGCACAGATTACCTCTTTTTGTTTGGTCTGTGTTTATTACAGCTTTCTTATTGATTCT
TTCACTTCCTGTTTTAGCTGGTGGTATCACTATGTTATTAACAGATAGAAATTTCAACACT
ACATTTTTTGATCCAGCAGGTGGGGGGGATCCAATTCTTTACCAACACTTATTC

>NC_024944

TACTTTATATATCATCTTCTCTATTTTTGCGGGTATGATTGGTACTGCTTTTTCTATGTTAA
TTAGACTAGAATTAGCAGGACCTGGAATTCAATATCTACAAGGTGATCATCAATTATATA
ATGTAATTGTTACTGCTCACGCATTTGTAATGATTTTCTTCTTAGTAATGCCTGCAATGATT
GGTGGATTCGGTAACTGGTTTGTTCCTTTAATGATTGGAGCTCCAGATATGGCTTTCCCTC
GATTAAATAATATTTCATTCTGGTTATTACCACCTTCTTTAATTCTTTTAGTAGCTAGTGCT
TTCGTAGAAAACGGAGCTGGTACTGGATGGACTGTTTACCCTCCATTATCAGGTATTGCTT
CTCACTCTGGTGGATCTGTAGATTTAGCGATCTTTAGTCTTCACTTATCAGGTATTTCTTCT
ATGTTAGGTGCAATGAACTTCATTACAACAATTATTAATATGAGAGCTCCTGGAATGTCA
TTCCATAAAATGCCTCTATTTGTATGGGCTGTATTAATTACAGCAGTTCTATTACTTCTATC
TTTACCTGTATTAGCCGGTAGAATTACTATGTTATTAACAGATCGAAACTTTAATACTTCA
TTCTACGAGCCAGCAGGTGGAGGAGATCCATTACTATACCAACAC

>NC_025284

CACACTTTACTTACTTTTAGGAATATGGTCAGGTCTAGTAGGCGCGGGCCTAAGAATAAT
AATCCGAATCGAACTAGGTCAACCCGGTTCTAGCTTTATAGGCAATGACCAACTCTATAA
TGTAATTGTAACCGCTCATGCTTTTGTGATGATTTTTTTTATAGTAATACCAATAATAATA
GGAGGATTTGGAAATTGAATATTACCTCTAATATTAGGAATTCCAGACATAGCCTTTCCC
CGAATAAATAACTTAAGATTTTGATTATTACCACCCTCACTCTTATTACTTTTAAGCTCAG
CAGGAGTAGAAAGAGGGGCCGGGACTGGGTGGACTGTATACCCCCCTCTTTCAAGAACC
TTAAGACATTCTGGCGCCAGTGTAGATCTAGCTATTTTTTCTCTTCACTTAGCAGGAGCAT
CTTCTATTCTAGGATCCATTAACTTTATTACAACTGTTATTAATATACGCTCACAAGGACT
ACAGATAGAACGACTTCCTTTATTTGTTTGATCAGTAAAAATTACAGCAATTCTTCTTCTT
CTGAGCCTCCCCGTTCTTGCTGGTGCAATTACAATACTCCTAACAGACCGAAATTTCAATA
CAGCTTTTTTTGACCCTGCTGGAGGTGGAGACCCTATTTTATACCAACATTTATTC

>NC_025293

TACATTATATCTTATTTTCGGTGCATTTTCTGGTATTATAGGTGCAATTTTATCATTATTTA
TACGTATGGAGTTAAGCCAACCAGGTAATCAAATATTAGGTGGTAATCATCAATTGTATA
ACGTAATAATAACAGCACATGGTTTATTAATGGTTTTTTTTTTAGTAATGCCCGTATTAAT
TGGGGGTTTCGGGAATTGGTTTGTTCCATTATTAATTGGAGCTCCTGATATGGCGTTTCCT
CGTTTAAATAATATTAGTTTTTGGTTATTACCTCCTTCTTTGTTGTTGTTATTAACCTCAGC

TTTTGTTGAAGTTGGAGCAGGAGTTGGTTGGACCGCGTATCCACCGTTATCAAGTCTTCAG
GGGCATGCTAGTCCTTCGGTTGATCTAGCTATTTTTAGTTTACATGTGTCTGGTGTAGGTA
GTATTTTGGGTGCTATTAATTTTATTACAACCATATTTAATATGCGTACCCCTGGTATGAC
AATGCATCGTTTACCTTTGTTTGTTTGGTCTGTTTTAATTACTGCTTTTTTATTGTTATTATC
ATTGCCGGTTTTTGCGGGTGCTTTAACTATGTTAATTACGGATCGTAATTTTAACACAACT
TTTTTTGATCCGGCGGGAGGTGGTGATCCTATTTTGTATCAGCATTTGTTT

>NC_025751

ATCACTATACATTTTATTTGGCGTAGTTGGAGGAATACTAGGAATCTCCCTATCCGCTCTA
ATCCGAATAATCCTATCTTCACCATGAACATTTAATGTAAATGGACTAATTTCAGAATACT
ATAATATAATCGTCACTGCTCACGCACTCACCATAATCTTCTTTATAGTGATACCTATACT
ACTAAGAGGATTCGGAAACTGACTAATCCCAATTATAGTAGGAGCAACCGATATATCCTT
TCCACGACTAAACAATCTAGGATTTTGATTACTACCTCCCGCTTTAATACTATTAAGCTCA
TCTATACTCATTAACATAGGAGCTGGAACAGGATGAACTATTTATCCTCCACTATCCACTT
GGCTCGGACACCCCAACCACAGTGTTGACTTAGTAATCTTCTCCCTACACCTAGCCGGCA
TCTCATCAATTGCAGGAAGAATCAACTTTATTACCACATGTTTCTTATCACGACCTGTTAT
CTACACACTAGAACGATTAAGCATATTTGTTTGAAGAGTATTAATTACATCATTCATACTA
ATTATTTCACTACCAGTTTTAGCAGGCGGAATCACCATACTCTTAACAGACCGTAACTTCG
GCACAACTTTCTTTGAAACCTCAGGAGGAGGAGACCCCATCCTATTTCAACATATATTC

>NC_026310

GGTGCTATATTTTGGCTTTGGCCTTTTATCAGGAGTTATCGGAACGGTTTTATCAGTAATT
ATTCGATGTGAGTTGTCTATGCCTGGCAGTCCAGTCTTACATGGTAATTACCAATTATTTA
ATGTTTTAGTAACAGCACATGCTTTTGTTATGATTTTTTTTATGGTCATGCCTATTTTAATT
GGAGGTTTTGGAAATTGATTTGTACCATTATTAATTGGGGCACCAGATATGGCGTTTCCAC
GATTAAATAATTTAAGTTTTTGATTTTTACCTCCTGCATTAATTTTATTATTAACATCAAGT
TATTTTGATCCAGGAGTAGGTACCGGTTGAACAGTTTATCCACCATTATCTGGTGCAATCG
CACATGCAGGACCTTCTGTAGATTTTGCAATTTTTAGTTTACATATTGCAGGAGCATCATC
TATTATGGGTGCTATTAATTTTATTACAACCATTGTTAATATGCGGCATCCAGGATGCGAA
TGAAATGGTTTAAATTTATTTGTTTGGTCTGTATTTATTACTGCAATTTTATTATTATTATC
ACTACCAGTTTTAGCAGGAGCAATTACTATGTTATTAACAGATCGAAATTTAAATACAGT
GTTTTTTGAGGCTACAGGGGGTGGAGATCCTGTTTTATATCAACATTTATTT

>NC_026516

ATTTATGTATATTCTGGTAAGGGTTTGGAGGGGCGTAATAGGGTTTAGGTTGAGTATAGT
AATTCGTTTAGAGTTGGGATCGGGAGGACAGTGATTAGGGGATGAGTATTTGTATAATTT
GGTTGTAACTAGGCATGGGGTGATAATGTTGTTCTTTTTGGTAATACCTATGTTTATGGGG
GGATTTGGGAATTGAATGATACCTATAATGTTAGGATTAGATGACATAGCTTTACCTCGG

TTAAATAACTTGAGGTTTTTAATAGTACCTGTAGCTTTAGTGTTATTTAGGGTATCTATGG
TGATTAAGGGAGGTAGGGCTGGATGGACTATGTACCCCCCTTTGATATTAAGTGAATACA
GATCTTCTGTTTCGGTTGATATGATGATTTTGAGTTTACATGTGGCAGGTTTGTCGTCATT
GTTAGGGTCAATTAATATTGTAGTTACTGGGGTGATTGGAAGGAAGATGGGTGGAAGAGT
GGATCAAATTCCTTTGTTAGTTTGAGCACTTATAATTACAGCAGTTTTAGTGTTGTTAACT
ATTCCTGTATTAGCTGCTGCTTTGACAATAATGTTGTTGGATCGTAATTTTAGTACGAGTT
ATTTTGATCCTGCAGGAGGAGGAAGCCCGCTATTATATCAACATTTATTT

>NC_026666

GACTATTTATTTTGTTTTTGGGATATGATCTGGAATATTAGGTCTTTCCATAAGTATTTTAA
TTCGGAGAGAGCTTTCTAGCCCTGGTAGTTTAATTAGAGATGATCATATTTTTAATGTTGT
GGTTACCTCACATGCGTTAATTATAATTTTTTTTATGGTGATGCCTATTTTAATTGGAGGG
TTTGGGAATTGGTTGATCCCTTTAATGCTAGGTAGCCCAGACATAGCCTTCCCTCGGATAA
ATAATTTAAGATTTTGATTATTGCCTCCATCTTTATTTATACTATTAATAAGAAGAATAGT
TGAGTCTGGAGTTGGGGCAGGATGGACCTTGTACCCTCCTCTTTCTGATAAATTAAGTCAT
TCTGGGGGTTCGGTAGATTTGTCAATTTTTTCTTTACATTTGGCCGGGGCTTCTTCTATTTT
GGGGGCTATTAATTTTATTACAACAATTTCTAATATACGAAGATTTAATTTAAAATGATTA
AAAATTACATTATTTTCTTGATCAGTTTTTATTACTGCTTTTTTGCTTCTTTTATCTTTACCT
GTATTAGCTGGGGCTATTACTATATTGTTAATAGACCGAAATATAAATTCTTCTTTTTTTA
ACCCAAGAGGTGGGGGGGATCCTGTATTATTTCAACATTTATTT

>NC_026983

AATAAATTATTTATGGTATGGTTTGACTACAGGTTTTGTAGCTTCTGGTTTAAGTATGATT
ATTCGACTAGAATTAGGTACTACTGAATCAGTTTTAATGAATGATCATACATATAATGTA
GTAATTACTGCACATGGTTTATTAATATTATTTTTTGTTGTTACACCTGTTTTAATAGGTAG
TTATGGTAATTATTTTGTACCTGTTATATTGGGTTGTCCAGACATAGCTTTTCCTCGAATG
AATAATTTAAGATTTTGAATAGCTATTCCTGCATTTTTATTGCTTATTAGAAGATCTATAA
TAGAAGGTGGAGCTGGAACAGGTTGAACTGTTTATCCTCCTTTATCGAGTATTGAATTTCA
TAGAAGAATTTCAGTAGATTTAGCAATTTTTTCTTTGCATTTTTCTGGAATATCTTTAATTT
TAGGTTCTGTAAATTTTATTGCTACTATTACTAATATACGAGTATCTTCTATGTATCTTATG
CGAATTAGACTTTACACATGATCTATTATAATTACTGCTGTTTTATTGATTATTTCTTTACC
TGTATTTGCAGGAGCTATTACGATGCTTTTAGTTGATCGAAATTTTAATTGTTCTTTTTTTG
ATCCTATAGGTGGTGGAAGTTTAATTTTATATCAACATTTATTT

>NC_026985

TATGCTTTATATCCTCTTTGGCCTTTGATCTGCTATGGTAGGGACCGGCCTAAGACTACTT
ATTCGAATTGAGCTAAGTGTAGCAACTAGCTGAATGGGAGACGATCAGCTATATAACGTG
ATTGTTACCGCTCACGCCTTTGTAATGATTTTCTTCTTTGTAATGCCTTTTATGGTAGGTGG

ATTTGGAAACAGTCTTCTACCTCTAATGATTGGAGCGCCTGACATGGCTTTTCCTCGTCTA
AATAATATGAGATTCTGATTACTACCTCCTTCTCTTACCCTTCTACTTGTTTCAGCATTAGT
TGAAAGAGGGGCAGGAACTGGATGGACTGTATACCCTCCTCTATCTGGCATTGTTGCCCA
TGCAGGTGGTAGGGTAGATTTTGCTATTTTCTCTCTTCACCTTGCAGGTGCCTCTTCTATTT
TAGGTGCTATTAACTTTATTGCCACTACGCTAAACATGCGAGGAGCAGGTGTTACTTTCG
AGCGACTTCCTCTTTTCGTTTGGGCTGCTATTATTACGGTAGTTCTTCTACTTCTATCTTTA
CCAGTACTTGCTGGTGGGATTACTATGCTACTGACAGATCGAAACCTAAATACTAGGTTT
TTCGATCCTGCAGGTGGTGGTGACCCTATTCTATACCAGCACCTTTTC

>NC_027000

GACTCTCTATTTCATCTTCGGTGCCATTGCTGGAGTGATGGGAACATGCTTCTCAGTACTG
ATTCGTATGGAATTAGCACGACCCGGCGATCAGCTCCTTGGTGGAAATCATCAACTTTAT
AATGTTCTAATAACGGCTCACGCCTTTTTAATGATCTTTTTTATGGTCATGCCGGCCATGA
TAGGTGGGTTTGGGAATTGGTTTGTTCCTATTTTAATAGGGGCACCTGACATGGCATTTCC
ACGCTTAAATAATATTTCATTTTGGTTGTTGCCACCAAGTCTCTTGCTCCTATTAAGCTCA
GCCTTAGTAGAAGTGGGGAGCGGGACTGGGTGGACGGTCTATCCGCCCCTAAGTGGTATT
ACCAGTCATTCTGGAGGAGCAGTGGATTTAGCCATTTTTAGTCTTCATCTATCGGGTATTT
CATCCATTTTAGGTTCTATCAATTTTATAACAACTATCTTCAACATGCGTGGACCCGGAAT
GACTATGCACAGATTACCCCTATTTGTGTGGTCCGTTCTAGTCACAGCATTCCTACTTTTA
TTATCACTTCCGGTACTGGCAGGGGCCATTACCATGTTATTAACCGATCGAAACTTTAATA
CAACCTTTTTTGATCCCGCTGGAGGGGGAGACCCCATATTATATCAGCATCTCTTT

>NC_027069

TACCCTTTATCTTGTATTTGGTGCCTGAGCCGGAATAGTGGGAACCGCCTTAAGCCTTCTT
ATTCGGGCCGAACTAAGCCAACCCGGGTCGCTTCTAGGTGATGACCAAATTTATAATGTT
ATCGTTACTGCCCACGCCTTCGTAATAATTTTCTTTATAGTAATGCCCATTCTCATTGGAG
GGTTTGGAAACTGACTTGTACCCTTAATGATTGGGGCCCCAGACATAGCATTCCCCCGAA
TAAATAACATAAGCTTCTGACTTCTACCCCCATCATTCCTGCTACTATTAGCCTCTTCTGG
TGTTGAAGCCGGGGCCGGGACAGGGTGAACAGTTTATCCCCCCCTTGCAGGGAATCTAGC
CCACGCAGGAGCATCAGTAGACCTAACAATTTTCTCCCTTCACTTAGCAGGTGTTTCATCA
ATTCTAGGGGCCATTAATTTTATTACCACAACCATTAATATGAAACCCCCAGCCATCTCCC
AATATCAAACACCTCTATTTGTTTGATCCGTACTTGTAACCGCCGTACTACTTCTTCTATC
ACTACCTGTTTTAGCCGCCGGGATTACAATACTCTTAACAGATCGAAACCTTAACACCAC
ATTCTTTGACCCCGCAGGAGGGGGGACCCAATTCTTTACCAACACTTATTC

>NC_027406

GACTCTATATTTCATCTTCGGTGCCATTGCTGGAGTGATGGGCACATGCTTCTCAGTACTG
ATTCGTATGGAATTAGCACGACCCGGCGATCAAATTCTTGGTGGAAATCATCAACTTTAT

AATGTTTTAATAACGGCTCACGCTTTTTTAATGATCTTTTTTATGGTTATGCCGGCGATGA
TAGGTGGATCTGGTAATTGGTTTGTTCCGATTCTGATAGGTGCACCTGACATGGCATTTCC
GCGATTAAATAATATTTCATTCTGGTTGTTGCCACCAAGTCTCTTGCTCCTATTAAGCTCA
GCCTTAGTAGAAGTGGGTAGCGGCACTGGGTGGACGGTCTATCCGCCCTTAAGCGGTATT
ACCAGCCATTCTGGAGGAGCAGTTGATTCAGCAATTTCTAGTCTTCATCTATCTGGTGTTT
CGTCCATTTTAGGTTCTATCAATTTTATAACAACTATCTCCAACATGCGTGGACCTGGAAT
GACTATGCATAGATCACCCCTTTTTGTGTGGTCCGTTCTAGTGACAGCATTCCCACTTTTA
TTATCACTTCCGGTACTGGCAGGGGCAATTACCATGTTATTAACCGATCGAAACTTTAATA
CAACCTTTTCTGATCCCGCTGGAGGGGGAGACCCCATATTATACCAGCATCTCTTT

>NC_027832

AACTCTATACTTCATTTTAGGAATTTGAGCCGGAATAATTGGGGCTGGAATAAGTCTTCTT
ATTCGAATCGAATTAAGACAACCTGGGTCATTCCTGGGAAGTGACCAACTTTATAACACA
ATTGTAACAGCACATGCATTCTTAATAATTTTTTTTCTAGTAATACCAGTATTTATTGGTG
GTTTTGGTAATTGACTATTACCACTTATATTGGGGACTCCAGATATAGCATTTCCACGCCT
AAATAATATAAGATTCTGACTATTACCCCCCTCACTCATCTTACTAGTCTCTTCTGCAGCA
GCGGAAAAAGGTGCAGGAACAGGATGAACAGTTTACCCACCACTAGCAAGTAACATTGC
GCATGCTGGACCATCAGTAGATCTGGCAATTTTCTCACTACACTTAGCAGGGGCATCATC
AATTCTAGGTGCAATCAATTTTATTACTACAGTAATTAATATACGATGATCAGGCCTACG
ACTAGAACGAATTCCCCTATTTGTATGAGCAGTAGTAATTACCGTAGTTCTACTACTTCTA
TCATTACCAGTACTAGCCGGTGCTATCACAATACTATTAACAGATCGAAACCTTAATACA
TCATTCTTTGATCCAGCAGGAGGGGGAGACCCAATTTTATATCAACACCTATTT

>NC_028002

TACTTTGTACATTTTATTTGGTATATGATCTGGGTTAGTTGGGACTGCTCTTAGTTTGCTTA
TTCGAGCTGAATTAGGACAACCAGGAGCTTTACTTGGTGATGATCAACTCTATAATGTGA
TTGTTACAGCTCATGCCTTTGTAATAATTTTTTTTCTTGTTATACCTATAATGATTGGTGGG
TTTGGGAATTGATTAGTTCCTTTAATACTTGGAGCTCCTGATATAGCTTTTCCTCGTTTAAA
TAATATAAGCTTTTGACTTTTGCCTCCAGCTTTATTATTATTACTTTCTTCAGCAGCTGTTG
AAAGTGGAGTAGGAACTGGATGAACAGTTTATCCTCCTTTATCAGGTAATTTAGCTCATG
CCGGTGGATCTGTAGATTTAGCAATTTTTTCGCTTCATTTAGCTGGTGCTTCTTCAATTTTA
GGAGCTGTAAATTTTATTACTACTATTATTAATATACGATGACGAGGAATACAATTTGAG
CGATTGCCTTTATTTGTATGATCTGTAAAAATTACTGCTATTTTATTACTTCTTTCTTTACC
TGTATTGGCTGGTGCTATTACTATACTTTTAACTGATCGAAATTTTAATACTGCTTTTTTTG
ATCCAGCAGGAGGTGGAGATCCTATTTTATATCAACATTTATTT

>NC_028054

TACATTATACATAATATTTGGTATATTTGGAGCCTTTATCGGAACTTCACTAAGAACAATA
ATACGATTAGAGTTATCCCAAACAGGAACTTTACTAGAAAATGACCATTTATATAATGTT
ATAGTAACTGCACACGCTTTAATAATGATATTCTTTTTCGTCATGCCAATTATAATTGGCG
GATTTGGAAACTGATTTATACCTCTATGCATCGGAGCCCCAGACATGGCATTTCCACGAC
TAAATAACATAAGATTTTGATTACTACCACCTTCCTTATTCTTACTACTATCTTCTAGCTTC
GTAGAAAACGGAGTAGGAACGGGATGAACTTTATACCCTCCCCTATCTGGTATACAAACA
CACTCCAGAAGAGGGGTAGATTTAGCAATATTTAGACTACATCTCGCAGGAATTTCATCA
ATTCTAAGATCAATTAACTTTATAACAACTATAATAAACATGCGAACTAGAGCAATAACA
ATATATCGTATACCTCTATTTATTTGAGCAATCTTTTTCACAGCACTTCTATTAATCTTATC
GCTTCCTGTACTCGCAGGTGGAATTACAATGCTATTAACAGATCGAAACTTCAATACAAC
TTTCTTTGACCCNGCAGGAGGAGGAGATCCAATTCTATTTCAACACTTATTT

>NC_028096

GACTCTCTATTTCATCTTCGGTGCCATTGCTGGAGTGATGGGCACATGCTTCTCAGTACTG
ATTCGTATGGAATTAGCACGACCCGGCGATCAAATTCTTGGTGGGAATCATCAACTTTAT
AATGTTTTAATAACGGCTCACGCTTTTTTAATGATCTTTTTTATGGTTATGCCGGCGATGA
TAGGTGGATTTGGTAATTGGTTTGTTCCGATTCTGATAGGTGCACCTGACATGGCATTTCC
ACGATTAAATAATATTTCATTCTGGTTGTTGCCACCAAGTCTCTTACTCTTATTAAGCTCA
GCCTTAGTAGAAGTGGGTAGCGGCACTGGGTGGACGGTCTATCCGCCCTTAAGTGGTATT
ACCAGCCATTCTGGAGGAGCAGTTGATTTAGCAATTTTTAGTCTTCATCTATCTGGTGTTT
CATCCATTTTAGGTTCTATCAATTTTATAACAACTATCTTCAACATGCGTGGACCTGGAAT
GACTATGCATAGATTACCCCTATTTGTGTGGTCCGTTCTAGTGACAGCATTCCTACTTTTA
TTATCACTTCCGGTACTGGCAGGGGCAATTACCATGTTATTAACCGATCGAAACTTTAATA
CAACCTTTTTTGATCCCGCTGGAGGGGGGGACCCCATCTTATACCAGCATCTCTTT

>NC_028626

TCTATACTATTTATGGTTTTCATTTTTATTTGGTAGTTATGGTTTTTTATTATCTGTTATTTT
ACGTACAGAATTATATTCTTCTTCTTTAAGAATAATTGCACAAGAAAATGTTAACTTATAT
AATATGATATTTACATTACATGGAATTATTATGATATTCTTTAATATAATGCCAGGATTAT
TTGGAGGATTCGGTAATTACTTCCTACCAATTTTATGTGGTTCTCCAGAACTTGCATATCC
AAGAATTAATAGTATATCTTTATTATTACAACCAATAGCTTTTATATTAGTAATTTTATCT
ACAGCAGCAGAATTTGGAGGAGGTACTGGATGGACTTTATATCCACCATTAAGTACATCA
CTTATGTCTTTATCTCCTGTTGCAGTAGATGTTATCATTGTTGGTCTTTTAGTATCTGGTAT
TGCTAGTATTATGTCTTCTTTAAATTTTATTACTACTGTAATGCATCTAAGATCTAAAGGTT
TAACACTTGGTATATTAAGTGTATCTACATGGTCATTAATAATTACATCTGTAATGCTATT
ATTAACATTACCTGTTTTAACAGGTGGTGTTTTAATGTTATTATCAGATTTACATTTTAATA
CATTATTTTTTGATCCTACATTTGCTGGAGATCCTATTTTATATCAACATCTATTT

>NC_029039

AACTCTGTATTTCCTCTTCGGTGCGCTTTCGGGACTGATTGGAACATGCTTTTCAGTTTTG
ATTCGTATGGAATTATCAACCCCAGGCGATCAACTGCTAGGCGGGAACCATCAACTATAT
AATGTTTTTCTTACGGCGCACGCTATAGTCATGATCTTTTTCATGGTGATGCCGGCTATGA
TAGGTGGTTCTGGAAATTGGCTTGTTCCGATTCTGATTGGGTCCCCCGAAATGTCCTTTCC
ACGCTTAAATAATATATCGTTCTGGTTGTTGCCTCCGAGTCTGTTGCTCCTATTGTGCCCA
GCAGTAGTGGAATTGGGTAGCGGCACTGGTTGGACGGTCTATCCGCCCCTAAGTGGGATC
ACCAGCCATTCGGGAGGCGCGGTCGATTTAGCAATTTCTAGTCTGCATCTTAGCGGGTTA
TCTTCCATTTTAGGTGCGATCAATTTTATTTCAACGATATCGAACATGCGTGGCCCTGGAA
TGACGATGCATAGATCACCCCTCTTTGTGTGGTCCGTTCCTGTGACTGCATGCCCCATAGT
GTTGGCTCTTCCCGTACTGGCTGGGGGACTAACAATGTTATTAACGGATCGAAACTTCAA
TACTACGTTTTCTGATCCAGCTGGCGGTGGAGACCCCATTTTATACCAGCATCTCTTT

>NC_029130

GACTCTCTATTTTGTTTTCGGCGCCGTTGCTGGAGTGATGGGCACATGCTTCTCAGTACTA
ATTCGTATGGAATTAGCACAACCTGGCGATGGGTTAATAACGGCCGGTGGGAATCATCAA
CTTTATAATGTGTTAATAACGGCTCACGCCTTTCTAATGATCTTTTTTATGGTTATGCCGGC
GATGATAGGTGGATTTGGGAATTGGTTCGTCCCGATTCTTTGCGGTGCACCTGACATGGC
ATTTCCACGATTGAATAATATTTCATTCTGGTTGTTGCCACCTTCGCTGTTGCTCCTATGTT
GCTCAGCCTTGGTAGAAGTGGGTAGCGGCACTGGGTGGACGGTCTATCCGCCCCTAAGCG
GTATTACCAGTCATTCAGGAGGAGCCGTTGATCTAGCGATTTTTAGTCTTCATTTATCCGG
GATTTCATCCATAAGCGGTTCTATTAATTTTATCACTACTATCTACAACATGCGCGGGCCT
GGAATTACTATGCATAGATTACCCCTATTTGTGTGGTCTGTTCTAGTGACAGCATTTCTAC
TCTTATTATCACTTCCGGTACTTGCAGGGGCAATTACCATGTTATTAACCGATCGAAACTT
TCATACAACCTTTTTCGATCCCGCAGGAGGGGGAGACCCGATATTATACCAACATCTCTTT

>NC_029220

CACTCTCTACATCTTCTTCGGTATTTGAGCAGCAATAGTCGGCACAGGATTAAGCATAATC
ATCCGATTAGAACTAACTCAACCTGGAGCCCTTTTAGGAGATGACCAAATCTACAACGTA
GTAGTAACAGCCCATGCTTTAGTAATAATCTTCTTCATGGCTATACCAATTATAATCGGAG
GATTTGGAAACTGACTCCTGCCTTTAATAATCGGGGCCCCCGACATAGCTTTTCCTCGTTT
AAATAATATGAGTTTTTGGCTCCTCCCCCCATCATTCTTATTGCTGGTAGCCTCAGCAGGG
GTTGAAAGCGGAGTAGGAACAGGATGAACCCTGTACCCACCTTTATCAAACAACCTAGCC
CACGCTGGAGGAAGTGTAGACTTAGCTATCTTCTCACTACATTTAGCAGGAGCATCATCT
ATTCTTGGGGCAGCCAACTTTATTACCACTTCAATCAATATACGAGCCCCAGGTATAACTC
TAGACCGACTGCCTTTATTCGTGTGATCAGTGATCATCACTGCAGTTCTACTCCTACTATC

ACTACCCGTCCTAGCCGGAGGAATCACTATATTACTAACAGACCGAAACCTAAACACCTC
ATTCTTTGACCCGGCCGGTGGAGGAGACCCAGTATTATTCCAACATTTATTT

>NC_029355

GACTCTCTATTGCATTTTTGGTGCCATTGCCGGAGTCATGGGTACATGCTTCTCAGTACTA
ATTCGTATGGAATTAGCACAGCCTGGCAATCAAATTCTTGGCGGAAATCATCAACTTTAT
AATGTGTTAATAACAGCTCACGCTTTTTTAATGATTTTTTTTATGGTTATGCCTGCAATGAT
AGGTGGATTTGGTAATTGGTTCGTCCCGATTCTTATAGGTGCACCTGATATGGCATTTCCA
CGATTAAATAACATTAGTTTTTGGTTGTTACCACCGTCACTGTTACTTCTCTTAAGTTCTGC
TTTGGTAGAAGTGGGCGCTGGTACCGGATGGACGGTCTATCCGCCGTTAAGTGGTATAAC
CAGTCATTCTGGAGGATCTGTGGATTTAGCCATTTTCAGTCTTCATTTATCGGGTGTTTCCT
CTATTTTAGGTTCTATCAATTTTATCACTACTATTTTTAACATGCGTGGGCCAGGAATGAC
CATGCATAGATTACCCCTATTCGTATGGTCCGTATTAGTGACAGCATTCCTACTTTTATTA
TCTCTTCCAGTATTGGCAGGTGCAATTACCATGTTATTAACTGATAGGAACTTTAATACAA
CCTTTTTTGATCCTGCAGGAGGAGGAGATCCAATATTATACCAGCATCTCTTT

>NC_029396

AGTGTTATATTTAATATATGCACTATTCGCAGGGTTAATAGGAACTGCCTTTTCTGTTTTA
ATTAGATTAGAACTGTCAGGACCTGGTGTTCAGTATATTGCTGATAACCAATTATATAAC
AGTATTATTACTGCTCACGCTATAATAATGATCTTCTTCATGCGTATGCCAGCTTTAATTG
GAGGTTTTGGTAACTTCTTACTTCCATTAGGATTAGGAGGACCTGATATGGGGTTCCCTAG
ACTTAATAATATTAGTTACTTATTATTAATACCTAGTATTGTTTTATTCTTATTTGCAGGTG
GAATTGAAAATGGTGTAGGTACAGGTTGAACTCTTTACCCTCCATTATCAGGAATACAAA
GTCATAGTGGTCCAAGTGTGGATTTAGCTATCTTTGGATTACACCTTTCTGGTATTTCAAG
TTTACTTGGAGCTATGAATTTTATGACAACTACATTTAATATGAGAAGTCCTGGAATAAG
ATTACACAAATTAATACTATTTGCATGAGCAGTTGTTATTACAGCTGTTTTATTATTATTA
TCATTACCTGTTTTAGCCGGTGGAATTACTATGGTTCTTACAGATAGAAACTTTAATACTT
CATTCTTTGAAGTAGCAGGTGGTGGTGATCCTATATTATACCAACATCTGTTT

>NC_029766

TGTTATGTATTTTATTGTAAGGTTTTGGGGGGGTTTATTAGGGTTTAGTTTAAGGGGGGTG
ATTCGGTTAGAGTTGGGGTGTCCTGGGAGATGGATGGGGGCTGAGGGGGTTTATAATATG
ATAGTTACTTCTCATGCAGTGTTGATAGTGTTTTTTTTGGTTATGCCTGTGTTTATGGGTGG
GTTTGGTAATTGATTAACCCCTATTTTGTTGGGGGTGAGAGATATGATTTATCCTCGTTTG
AATAATTTTAGGTTTTTGTTAGTGCCTGTGGCTTTAAGGTTATTTTTAGGCTCTATGTTTAT
TGAGGGCCCACAAGCAGGGTGGACATATTACCCGCCCCTTTCTTTAGGTGAGTTTAGGGG
ATCAGTGAGTGTGGATGCTTCTATTTTAAGTTTACATTTTGTGGGTTTATCTTCTATTTTAG
GTTCTATTAATATTATTTCTACTTTTTTGGGGGCAGTATTTAGCTCTTCGATTAAAATAGA

ACAGGTGGCTTTATTGGTGTGGGCTTTAGTGGTAGCAGCTGGTATGATTTTACTTACGGTGCCGGTGTTGGCGGGGGCGTTAGTTATGTTATTGTTGGATCGTAATTTTAATTCTAGGTTTTTTGATCCGTCAGGTGGGGGGAGGTTGGTGTTGTATCAACACCTGTTT

>NC_029886

AACATTATATCTAATTTTTGGATTCTTATCAGGAATAATTGGAACAACATTTTCAGTAATAATCCGAATGGAATTAGCATCATCAGGAAATCAAATCCTAAATGGAAATCACCAATTATATAACGTAATAGTAACAGCACATGCGTTCTTAATGATTTTCTTTATGGTAATGCCTATTTTAATAGGTGGATTTGGTAACTGGTTCGTTCCAATCTTAATTGGTGCTCCAGATATGGCATTCCCAAGATTAAATAATATTAGTTTCTGGTTATTACCACCATCATTAATTCTATTATTAACATCTGCTTTTGTTGAAGTTGGTGCAGGTACTGGATGGACAGTTTATCCACCATTATCATCAATTCAAGCACATTCAGGTGGGGCAGTTGATTTAGCAATTTTCAGCCTACATTTATCAGGTATTTCATCATTACTAGGATCAATTAACTTTATTACAACAGTGATAAATATGAGAGCTCCTGGAATGGGATTTGCTCAATTACCATTATTTGTTTGGTCTGTATTCTTTACTGCTTTCTTATTGTTATTATCATTACCAGTTCTAGCTGGAGGAATTACAATGTTATTAACTGATAGAAACTTCAATACTTCATTCTTTGATCCTGCTGGAGGTGGTGATCCAATCTTATATCAGCACCTATTT

>NC_030255

TACTTTATATTTTATTTTTGGTGCTTGAGCAGGAATAGTAGGAACATCTTTAAGGTTAATTATTCGAGCTGAATTAGGACAACCAGGAAGTTTAATTGGAGATGATCAGATTTATAATGTTATTGTAACTGCTCATGCTTTTGTTATAATTTTTTTTATAGTTATACCTATTATAATTGGTGGTTTTGGTAACTGACTTGTTCCTCTTATATTAGGAGCACCTGATATAGCTTTTCCTCGAATAAATAATATAAGGTTTTGACTTCTACCTCCATCTTTAACTCTCCTTCTTATAAGAGGTATAGTTGAAAGAGGTGTTGGTACAGGTTGAACAGTTTATCCTCCTTTAGCTGCAGGTATTGCTCATGCAGGAGCATCTGTTGATTTAGGTATTTTTTCTCTACATCTCGCTGGGGTGTCTTCTATTTTAGGTGCAGTTAATTTTATAACAACAGTTATTAATATACGACCCTCTGGTATAACAATAGATCGTATACCTTTATTTGTATGATCTGTTTTTATTACAGCTATTTTACTTCTTCTATCTTTACCAGTATTAGCAGGAGCTATTACTATATTATTAACAGACCGAAATTTAAACACTTCTTTTTTTGATCCGGCTGGAGGTGGAGATCCTATTTTATATCAACATTTATTC

>NC_030337

AACTCTTTACCTGGTTTTTGGTGCTTTTTCAGGAATGGTGGGTACAGCTTTTAGTATGCTTATCCGTCTTGAATTATCTGCTCCCGGCACTATGCTAGGAGACGACCAACTCTACAACGTTTTGGTCACAGCTCATGCTTTTGTTATGATTTTCTTCTTAGTCATGCCTGTTATGATTGGCGGGTTTGGAAATTGGTTGGTCCCCCTTTACATTGGTGCCCCTGATATGGCTTTTCCTCGGCTTAACAATATTAGTTTTTGATTGCTACCTCCTGCCCTTATTTTATTATTGGGGTCTTCTTTAATTGAACAAGGAGTAGGAACAGGATGGACAGTTTACCCTCCCTTATCTGGAATCCAAGCCCATTCCGGTGGGGCTGTAGATATGGGAATTTTTAGCCTCCATTTAGCAGGTGCTTCTTCTATCT

TAGGAGCTATGAATTTTATTACAACTATATTCAACATGCGAGCCCCTGGAGTTACTTGGG
ACAAACTCCCTCTTTTTGTTTGATCTGTCCTCATTACTGCTTTTCTTCTACTTCTATCTCTTC
CTGTATTAGCTGGAGCTATTACTATGCTTCTAACAGACCGGAATTTTAACACTTCTTTCTT
TGACCCTGCTGGGGGAGGAGATCCCATCCTTTTCCAACATCTATTC

>NC_030635

AACATTATATTTAATATTTGCCTTATTTTCTGGATTATTAGGTACAGCATTTTCTGTATTAA
TAAGATTAGAATTAAGTGGACCAGGAGTTCAATTTATAGCAAATAATCAATTATATAACA
GTATAATAACAGCTCACGCTATATTAATGATATTCTTCATGGTTATGCCTGCTTTAATAGG
AGGGTTTGGTAACTTCTTAATGCCTTTAATGGTAGGAGGTCCTGATATGGCATTTCCAAGA
TTAAATAATATTAGTTTCTGATTATTACCACCTAGTTTATTATTATTAATATTCTCAGCTTG
TATAGAAGGTGGTGTAGGTACAGGGTGAACTCTTTATCCTCCATTATCAGGATTACAAAG
TCATAGTGGACCTAGTGTAGATTTAGCAATATTTGCTTTACACTTATCAGGGGTAAGTAGT
TTATTAGGTGCTATTAACTTTATAACTACAATAGCTAATATGAGAACACCTGGTATAAAA
TTACATAAATTAACTTTATTTGGATGAGCAGTAGGTATTACAGCTATATTATTATTATTAT
CATTACCTGTATTAGCTGGTGGAATTACTATGATATTAACAGATAGAAACTTTAATACATC
ATTCTTTGAAGTAGCTGGTGGTGGAGATCCTATATTATTCCAACATCTTTTC

>NC_030753

GACTCTATATTTCATCTTCGGTGCCATTGCTGGAGTGATGGGCACATGCTTCTCAGTACTG
ATTCGTATGGAATTAGCACGACCCGGCGATCAAATTCTTGGTGGGAATCATCAACTTTAT
AATGTTTTAATAACGGCTCACGCTCCTTTAATGATATTTTTTATGGTTATGCCGGCGATGA
TAGGTGGATTTGGTAATTGGTTTGTCCCGATTCTGATAGGTGCACCTGACATGGCATTTCC
ACGATTAAATAATATTTCATTCTGGTTGTTGCCACCAAGTCTCTTGCTCCTATTAAGCTCA
GCCTTAGTAGAAGTGGGTAGCGGCACTGGGTGGACGGTCTATCCGCCCTTAAGTGGTATT
ACCAGCCATTCTGGAGGAGCAGTTGATTTAGCAATTTTTAGTCTTCATCTATCTGGTGTTT
CATCCATTTTAGGTTCTATCAATTTTATAACAACTATCTTCAACATGCGTGGACCTGGAAT
GACTATGCATAGATTACCCCTATTTGTGTGGTCCGTTCTAGTGACAGCATTCCTACTTTTA
TTATCACTTCCGGTACTGGCAGGGGCAATTACCATGTTATTAACCGATCGAAACTTTAATA
CAACCTTTTTTGATCCCGCTGGAGGGGGAGACCCCATATTATACCAGCATCTCTTT

>NC_030900

GACTCTACATTCAGTCTTCGGTGCCATGGCCGGAGTAATGGGGACATGCTTCTCGGTACT
AATCCGTATGGAATTAGCGCAACCTGGCAATCAGATTCCTGGTGGAAATCATCAACTTTA
TAATGTGTTAATAACGGCTCATGCCTCTCTAATGATATTTTTTATGGTTATGCCGGCGATG
ATAGGTGGATTTGGGAATTGGTTCGTTCCGATTCTTATAGGTGCACCTGACATGGCATTTC
CGCGATTAAATAATATTTCCTTTTGGTTGTTGCCGCCTTCGCTGCTGCTTCTACTAAGCTCA
GCTTTGGTAGAGGTGGGTAGCGGCACTGGGTGGACGGTCTATCCGCCCTTGAGTGGTATA

ACCAGTCATTCTGGAGGAGCTGTTGATTTAGCCATTTCTAGTCTCCACTTATCAGGTGTTT
CATCCATTTCAGGCTCTATCAATTTTATCACTACCATCTTCAACATGCGTGGACCTGGAAT
GACCATGCATAGATTACCCCTCTTCGTGTGGTCTGTTTCAGTGACAGCATTCTCACTCTTA
CTATCCCTCCCAGTACTGGCAGGTGCAATCACCATGTTATTAACCGATCGAAACTTTAATA
CCACCTTCTTTGATCCCGCCGGAGGAGGGGACCCTATCTTATACCAGCATCCCTCT

>NC_031407

AACCCTATATTTTATTTTCGGCATTTGAGCCGCCACAGTGGGAAGATCACTTAGTATAATT
ATTCGTTCCGAACTTAGAGAACCCGGCTCACTGTTTGCCGAAGAGCAACTTTACAATGTC
ACAGTAACCAGACATGCCTTTATTATAATTTTTTTTTTGTTATACCTATTCTGATCGGAGG
CTTTGGAAACTGATTAGTCCCACTTATAATTGGGGCGCCGGATATAGCTTTCCCACGAAT
AAATAATTTAAGATTTTGACTTCTGCCCCCCTCTTTTCTATTAATCTCAACAAGCACAATA
GCTGAACAGGGGGCAGGAACGGGATGAACTGTATACCCGCCTTTATCCAATTATTTTGCT
CATAGTGGCCCTGCCGTAGATTTAACCATTTTCTCTCTCCATATTGCTGGCGTCTCTTCAAT
TTTGGGAGCAATCAATTTTATTTCCACAATCATTAACATGCGAACACCTGCCATAAGAAT
AGAAAACATACCATTATTTGTGTGATCGGTTTTAATTACAGCCGTTTTACTGCTTTTGGCC
CTTCCTGTACTAGCTGGTGCTATTACCATACTTTTACTTGACCGAAATTTTAATACTTCTTT
TTTCGATCCAGCAGGGGGAGGGGACCCTATTCTATACCAACACCTATTT

>NC_031417

TTCCCTTTATTTAGTTTTTGCTATGTTTGCTGGTGTAATAGGGACAACTTTTTCTGTTTTAA
TGAGGATGGAATTAAGTGCCACTGGTGATCAATTTTTTGATGGTAGTTATCAACATTACA
ATGTTGTTGTAACAGCTCATGGTTTAATTATGGTATTTTTTTTAGTTATGCCGGCACTAATT
GGTGGTTTTGGTAATTGGTTTATTCCTTTATTATTAGGAGCTCCAGATATGGCTTTTCCTAG
ATTAAATAATTTAAGTTTTTGATTAATGCCTCCATCTTTTTTATTATTATTATTATCTTCATT
AGTTGAAACAGGGTCAGGTACTGGTTGAACTGTTTATCCACCATTATCTAGTGTAGTTGC
ACATTCAACTCCTTCCGTAGATTTAGCTATATTTAGTTTACATTTAGCAGGTATAGCATCA
ATTGCAGGATCTATTAATTATATTACAACTATTTTTAATATGAGAGTTGCTAAATATGATA
TGTTTAGATTACCTTTATTTATTTGAGGTATGCTTATTACTGCTTATTTATTAGTTTTTACTT
TACCTGTTTTAGCTGGTGCTATCACTATGCTATTAACTGATAGAAATTTTAATACTTCTTTT
TTTGATCCCGCTGGAGGTGGAGATCCTATACTATATCAACATTTATTT

>NC_031438

TATAATATATAGTGTAGTAGGGGTATGAGCTGGATTTGTAGGTTTAGGTTTAAGAATTCT
AATTCGTATTCAGTTATCAGACCCATACTTTAATATAATCCCTTTTGAAGTATATAATTAC
GTTATCACCAGGCATGGTATTATAATGATATTTTTCTTCTTAATGCCAGTTTTAATTGGTG
GGTTTGGAAATATACTATTACCTATACTTTTAAATTTAAATGATTTAAACTTACCTCGGTT
AAATGCTTTAAGAGCATGGTTATTAATGCCATCTATGGTTTTAGTATTTGCTAGTATATGA

TTCGGTAGAGGTACTGGATGAACTTTTTATCCCCCGTTATCTGGTGCAAGATTTAGCCCTA
GCGTAGGGACTGACTTTTTAATGTTTTCTTTACATTTATCTGGTATTTCTAGGATATTTAGT
TCATTAAATTTTATTTGTACTATTATTAGTGCATGAGGAGTTTCAGTAAATATAAAGGACA
CAGCTATAGTGGTATGAGCTTATCTGTTTACTTCTATTTTACTTATTTTATCTTTACCAGTT
TTAGCAGCTGGTATAACAATGTTACTTTTTGATCGTAACTTTAATTCTTCTTTTTTTGATCC
AGTAGGTGGAGGTGATCCAGTACTTTTTCAACATTTATTC

>NC_031504

AAGTTTGTACTTAATCTTTGGTGCCTTCTCCGGTGTTCTAGGAACTTGTATGTCTATGCTA
ATTAGAATGGAATTAGCACATCCAGGAAATCAAATTTTAGCTGGAAATCATCAATTATAT
AATGTACTTATAACAGCCCATGCTTTTTTAATGATCTTTTTTATGGTAATGCCAAGTTTAA
TTGGAGGTTTTGGTAATTGGTTTGTACCAATAATGATTGGTGCACCTGACATGGCCTTTCC
AAGATTAAATAATATAAGTTTTTGGTTATTACCCCCTTCACTTCTTTTATTACTTAGCTCCG
CTTTAGTAGAAGTAGGTGCAGGTACTGGATGGACAGTTTATCCACCTTTAAGTAGTATCA
CAGCCCATTCTGGGGGATCAGTGGATTTAGCTATCTTCAGTCTTCATTTATCAGGATTAAG
TAGTTTATTAGGTGCAATTAACTTTATTACTACAATATTGAATATGCGAGGTCCTGGTATG
AGCATGCATCGTATTCCTTTATTTGTATGGTCAGTTTTTATAACAGCTATCTTACTCCTTCT
TTCTCTTCCAGTATTAGCTGGTGCTATCACAATGCTTCTAACAGATCGTAACTTTAATACT
AGTTTTTTTGATCCCGCTGGAGGAGGAGATCCAATTCTTTTTCAACATTTATTC

>NC_031872

AAGATTATATTTTTTTCTAGGGGTTTGATCTGGAATGGTAGGAACAGCCTATAGAGTAATT
ATTCGATTAGAGTTAATACATCCTGGAAGATGGGGTGGGGATGCACATTTTTTTAATAAA
GTTATTACTGCTCATGCATTAATTATAATTTTTTTTTTAGTAATACCAGTGTTGATTGGAGG
GTTTGGAAATTGATTAATTCCTTTAATGTTAAATGTCCCAGATATAGCATTTGCACGATTA
AATAATTTGAGGTTTTGAGTTTTACCCCCTGCATTATTTTTATTAGTAGTGAGATTTGGAA
CTGACGGGGGTGTAGGGGCTGGGTGAACAATTTATCCCCCTTTAGCAGGATTAACTGGTC
ATGGAGGGGGTTCAATAGATTTAGCTATTTTAGCTCTTCATACAGCTGGGGCCGGATCAA
TTTTAGGTTCAATTAATTTTTTAGTAACTCTTTTATGTATGCGAGGAATTGCAGTGGGAAT
TGATCAAATAACTTTTTTTAATGTTTCAATTTTAGTAACTACATTTTTACTTTTAGTATCTG
TTCCTGTATTAGCTGGGGCTATTACAATATTGTTATTAGATCGAAATTTTAATACTTCGTTT
TATGAGGTTGGAGGTGGGGGAGACCCAGTTTTATTTCAACATATTTTT

>NC_031873

AACATTGTATTTTTTATTGGGTTTATGGTCTGGTATGGTTGGAACAGCAATAAGGTTTATT
ATTCGTATAGAATTATCTCATCCTGGTATGTGAATTGGTAGTGGTCATTTTTATAATAAAG
TTGTTACGGCTCATGCATTAATTATAATTTTTTTTATGGTGATACCTGTTCTTATTGGAGGT
TTTGGTAATTGATTTTTACCTTTAATATTAAATGTGCCTGATATGGCATTTGCTCGATTAA

ATAATTTGAGATTTTGGGTTGTTCCTCCTGCATTATTATTATTAGTTATTAGTTTTGATACT
GATGGTGGTGTTGGGGCAGGTTGAACAATTTATCCTCCGCTAGCTGGGTTAACTGGCCAT
GGTGGGGCTTCTGTTGATTTAGCTATTTTATCTTTACATATGGCTGGGGCTGGGTCTATTTT
AGGCTCAATTAATTTTTTAACAACTATATTACATATACGAAGATTTGGAATTGGGGTAGA
TCAATTACAATTTTTTAATACAGCTATTTTAGTTACTACTTTTTTATTAGTTGTTTCTGTTCC
TGTTTTAGCTGGGGCTATTACTATACTATTAATAGACCGTAATTTTAATACTTCATTTTATG
AGGTTGGAGGTGGTGGGGACCCTATTTTATTTCAACATTTATTT

>NC_032361

CACTCTCTATTTTATCTTAGGTACATGATCAGGACTCTTAGGCACCTCCATAAGTCTACTA
ATTCGAGCAGAATTAGGGCAACCTGGAGCCCTTCTTGGTAGAGATCAACTATATAACACT
ATCGTTACCGCCCATGCTTTCCTAATAATTTTTTTCCTTGTAATACCAGTAATAATCGGAG
GATTCGGCAACTGACTTGTTCCTCTTATACTAGGGGCCCCAGATATGGCATTTCCCCGTCT
AAACAACATAAGATTCTGACTGCTCCCCCCTTCTCTAACTCTCCTCCTAGCAAGGGCTGCA
GTAGAAAAAGGTGTTGGAACAGGATGAACTGTATACCCCCCTTTAGCCGGCAACATTGCT
CATGCAGGACCGTCAGTAGACTTAGCTATCTTCTCTCTTCACTTAGCAGGAGTCTCCTCAA
TTCTAGGAGCCCTTAACTTTATTACAACAGTTATTAACATACGATCCAAAGGCCTACGACT
AGAACGAGTCCCTTTATTTGTGTGATCAGTTATAATTACTGCTATTCTTCTCCTACTCTCTC
TTCCTGTTTTAGCAGGAGCCATCACTATACTTCTTACAGATCGTAATCTTAACACTGCCTT
CTTTGATCCTGCAGGAGGAGGAGACCCCGTACTATATCAGCACTTGTTC

>NC_033379

AAGCCTTTACCTAATTTTTGGTATTTGATGTGGGTTGGTTGGAACTGGCTTGAGAGTTTTA
ATTCGGAGGGAGTTAGGTCAGCCTGGAAGATTTATAGGGGATGATCATTTGTATAATGTT
GTAGTGACTGCACATGCTTTTATTATAATTTTTTTTTAGTAATGCCTATGATAATTGGGG
GGTTTGGGAATTGGTTGGTTCCTTTAATATTAGGGGCGCCTGATATAGCTTTTCCTCGACT
GAATAATATAAGATTTTGGTTGTTACCGCCTTCTTTGATTTTGTTATTGTCTTCTGCTGCTG
TAGAATATGGGGTTGGAACTGGGTGGACTGTTTATCCGCCACTATCTAATTATGATATTCA
TTGGGGGGTTGCAGTTGATTTGGCTATTTTTTCTTTACATTTGGCGGGTGTGTCTTCTATTT
TGGGAGCAGTGAATTTTATTACTACTGTAATTAATATGCGATGAAGGGGGATGACTCTTG
AACGGGTTCCTTTGTTTGTATGATCCGTAAAAATTACGGCTATTTTATTATTACTATCTCTT
CCGGTATTAGCGGGGGCGGTTACAATGTTATTGACTGATCGAAATTTTAATACTTCTTTTT
TTGATCCGGCTGGGGGAGGGGATCCTATTTTATTTCAGCATCTTTTT

>NC_033540

AACTTTATATTTTATTTTTGGGATTTGAGCAGGAATAGTAGGAACATCATTAAGATTATTA
ATTCGAGCTGAATTAGGTAACCCAGGATCATTAATTGGGGATGATCAAATTTATAATACT
ATTGTAACAGCTCATGCATTTATTATAATTTTTTTTATAGTAATACCTATTATAATTGGAG

GATTTGGAAATTGATTAGTACCATTAATATTAGGAGCTCCTGATATAGCTTTCCCACGTAT
AAATAATATAAGATTTTGATTATTACCTCCTTCATTAACTTTATTAATTTCAAGAAGAATT
GTAGAAAATGGAGCAGGAACAGGATGAACAGTATACCCCCCACTTTCATCTAATATTGCT
CATGGTGGAAGATCTGTAGATTTAGCAATTTTCTCACTTCATTTAGCTGGTATTTCATCTA
TTTTAGGAGCAATTAATTTTATTACAACTATTATTAATATACGAATTAATGGATTATCTTT
TGATCAAATACCTTTATTTGTTTGAGCAGTAGGAATTACAGCATTATTATTATTATTATCT
TTACCTGTATTAGCTGGAGCTATCACAATACTTTTAACAGATCGAAATTTAAATACATCAT
TTTTTGATCCTGCTGGAGGAGGAGATCCTATTTTATACCAACATTTATTT

>NC_033969

TTTGCTATACTTGATTTTTGCCTTCGTCGGAGGTTTAATCGGTACTTCTCTTAGTATGTTGA
TTCGTTACGAGTTGGCTTTGCCTGGTCGTGGTTTGTTAGACGGTAATGGACAACTATACAA
TGTTATCATTACTGGTCATGGAATTATCATGCTATTGTTTATGGTAATGCCAGCCCTGTTC
GGTGGTTTTGGTAACTGGTTGCTACCAATTATGATTGGTGCTCCAGATATGGCATTTCCCC
GCCTAAACAACATTAGTTTCTGGTTGAACCCACCTGCTTTGGCTTTATTGTTGTTGTCTACT
TTGGTAGAGCAAGGTCCAGGTACTGGCTGGACCGCTTATCCCCCACTAAGTGTACAACAC
AGTGGCTCTAGCGTAGATTTGGCTATTTTGAGTTTGCATTTGAACGGCTTGAGTTCTATCT
TAGGTTCTGTAAACATGCTAGTAACTGTAGCCGGTTTGCGTGCTCCTGGAATGAAATTGTT
GCACATTCCATTGTTTGTATGGTCTATTAGCTTTACAGCTGTATTGGTTATTTTGGCTGTAC
CTGTATTGGCCGCTGCTTTGGTTATGTTGTTGACTGATCGTAATTTGAACACTGCTTACTTC
TGTGAGTCTGGTGATTTGATTTTGTATCAGCATCTTTTC

>NC_034000

TATTTACTATGTATGGTTTGCCTTTCTTTTCTCAATTGTAGGTACATTACTATCAGTATTAA
TTAGACTTGAACTAAGTTCCTCTGGATTACGTGTTGTAGCATTAGAAAATCAAAATTTCTA
CAATTTAGCATTCACACTACATGGTGCTATTATGATTTTCTTTGTAGTTATGCCAGGTCTA
TATGGTGGATATGGTAACTATTTCTTACCAATTTATCTAGGAGCATCTGAGGTAGCTTTCC
CTAGAGTAAATTGTGTTTCATTATTACTAGTTCCAATTGCATGGGTTATAGTAAGTACTTC
ACTAATTTCTGAATTTGGTTCAGGTGTTGGTTGGACACTATACCCTCCATTAAGTACATCT
TTAATGTCATTATCTCCAACCTCAGTAGATTTAATTGTATTTGGTTTAGCTTTATCTGGTAT
TTCTAGCTTCTTATCTTCTATTAATTTCTTAACTACAATTGCTGTATTAGGTGTTACTAATG
GTTCAAAACCATGGTGTCTATTTACTTGGGCTATTGTATTTACAGCTATTATGCTACTTGG
AACACTTCCAATTCTTACAGGTGGATTACTAATGCTGGTATTAGACTTACATCTAAATACT
CAATTCTACGATGCCTCTTTTAATGGTGATCCAGTACTATATCAACATCTATTC

>NC_034947

TAGTCTATATTTTATTTTTGGTGCTTGAGCTGGTCTTATTGGCACTCTTTTAAGAGTGTTAA
TCCGACAAGAACTTCGTAATCCAGGTGCAGGTTTAATTAGAGAAAAAAGATATAATGTTA

CTATTACATCACATGGTTTAATTATAATTTTTTTTTTTGTAATACCAGTGTTGATAGGGGGT
TTCGGAAATTGATTACTGCCGTTAATACTTGGTTGTGCTGATATAGCTTTTCCTCGTTTAA
ATAATTTTTCTTTTTGATTACTTCCCCCCAGTTTTAGTTTATTAGTTCTTAGAAGACTCGCA
GAAAGAGGAGTTGGAGGAGGGTGAACACTTTATCCCCCCATAAGTGATTTAATAGGTCA
ACCAGGAATAAGAGTTGATTTGGCTATTTTCAGATTACATATTGCGGGAGCTTCCTCCATC
GGAGGTTCTATTAACTTTTTATGTACTATTATTAATTTACGAACTCCTGGAATAACATGAC
AAGAATTACCTCTTTTTATTTGAAGAGTGTTTTTTACAGCTATTTTATTAGTTCTTTCTTTA
CCAGTTTTTGCAGGTGGGATTACAATACTTCTAACAGATCGTAATTTCAATACTAGATTTT
TTACTCCAGGAGGAGGAGGAGATCCAGTTTTATTTGTACATTTATTT

>NC_035256

AACTATTTACTTTATTTTTGGTATGTGAAGAGGCCTAGTTGGGTCAGGTTTAAGTGTTTTA
ATTCGACTTGAACTATCAACACCAGGAATTTTTATTGGAGACGATCAGATTTATAACGCA
GTGGTAACCGCTCACGCTTTTGTTATAATTTTCTTTATGGTCATACCAGTTATGATTGGAG
GTTTCGGGAACTGACTAATTCCACTTATAATTAATTCTGTGGATATGGCTTTCCCTCGAAT
AAACAACATAAGATTTTGACTTCTGCCTCCTTCATTATTTATATTGTTAGTATCTTTCTTCC
TTGAAAAAGGAGTTGGTACTGGGTGAACAGTTTACCCACCACTATCGGGTAATGTTTCTC
ATGCTGGACCTGCTGTAGATTGTGCAATTTTCTCTCTACATTTGGCCGGTGCTTCATCTATT
ATAAGTTCAATCAATTTCATTACAACTATTTCATTTTTCCGAGGAGACAAAGTAAAATGA
GAAAGAGTAAGTTTATTTGTTTGAGCAATTTTAATTACAGTCTTTTTATTATTGCTTAGCCT
GCCTGTTCTAGCTGGGGGAATTACAATACTATTAACGGATCGTAATTTAAATACTTCTTTC
TTTGATCCTGCTGGTGGAGGAGACCCTATTCTTTATCAACACCTATTC

>NC_035351

AATTTTATACTTAATTTTTGGTGCTATATCAGGGTTATTAGGAACTGTTGTTAGTGTAATT
ATGCGTATGGAATTAACGTACCCTGGGCCTCAATTTCTTGAGGGGAATTATCAATTATAC
AATGTTTTAATAACGGCACATGCATTACTAATGATATTTTTTTTTGTGATGCCTATATTAA
TTGGTGGATGAGGTAACTGATTTGTACCTATAATGATTGGCGCTGTTGATATGGCTTTTCC
TAGATTGAACAATATTAGCTTTTGGTTATTACCCCCAGCATTAATGTTACTATTACTTTCG
GCTCTAGTAGAAGTTGGGGCTGGTACTGGTTGAACTATATATCCTCCCTTAAGTTCTATAC
AAAGTCATTCAGGAGCTGCTGTTGACTTGGGAATTTTTTCACTTCACGTTTCTGGTGCTGC
TTCAATTTTAGGTGCTATAAATTTTATTGTAACTATTTTTAATATGAGAAGTCCAGGTCAA
AGTTTACATCGTATGCCTTTATTTGTATGATCAGTGTTAGTTACAGCATTTTTATTATTATT
ATCGCTTCCTGTTTTTGCTGGAGGTATTACAATGTTATTGACTGATCGAAACTTTAATTCA
ATGTTTTTTGATGCTAAAGGAGGAGGAGATCCCATATTGTTTCAGCATTTATTT

>NC_035616

GACTCTATATTTCATCTTCGGTGCCATTGCTGGAGTGATGGGCACATGCTTCTCAGTACTG
ATTCGTATGGAATTAGCACGACCCGGCGATCAAATTCTTGGGGGGAATCATCAACTTTAT
AATGTTTTAATAACGGCTCACGCTTTTTTAATGATCTTTTTTATGGTTATGCCGGCGATGA
TAGGTGGATCTGGTAATTGGTCTGTTCCGATTCTGATAGGTGCACCTGACATGGCATTTCC
ACGATTAAATAATATTTCATTCTGGTTGTTGCCACCAAGTCTCTTGCTCCTATTAAGCCCA
GCCTTAGTAGAAGTGGGTAGCGGAACTGGGTGGACAGTCTATCCGCCCTTAAGTGGTATT
ACCAGCCATTCTGGAGGAGCAGTTGATTCAGCAATTTCTAGTCTTCATCTATCTGGTGTTT
CCTCAATTTTAGGTTCTATCAATTTTATAACAACTATCTCCAACATGCGTGGACCTGGAAT
GACTATGCATAGATCACCCCTATTTGTGTGGTCCGTTCCAGTGACAGCATTCCCACTTTTA
TTATCACTTCCGGTACTGGCAGGGGCAATTACCATGTTATTAACCGATCGAAACTTTAATA
CAACCTTTTCTGATCCCGCAGGAGGGGGAGACCCCATATTATACCAGCATCTCTTT

>NC_035722

TTACCTAATTTTTGCAATTTTTAGCGGAATTATTGGTACCGCACTTTCAATGATAATTAGA
GCAGAGCTTGCTCTACCGGGTACACAGGTACTATCTGGTAATTATCAATTATATAACGTT
GTAATTACAGCGCATGCTTTATTCATGATTTTTTTCATGGTAATGCCCGCCCTAATGGGTG
GATTTGGAAATATTCTGGTGCCCGTAATGATTGGAGCTCCAGATATGGCATTTCCACGAC
TGAATAATATTAGCTTTTGGCTGTTGCCGGTAAGTTTACTATTATTATTAAGTTCAGCATT
GGTTGAGACTGGTGCTGGTACAGGATGGACCATTTACCCCCCTTTATCATCACTAGTAGG
AATGCCAGGTGCATCTGTAGACTTAGCCATTTTTTCATTACATATTTCGGGATTAAGTAGT
TTACTAGGGTCTATAAACTTTATTACAACTATTTTTAATATGCGAGCTCCGGGAATGAAAA
TGCATAGGGTACCCCTATTTGCTTGGAGTGTGCTTATTACTAGTTTCCTACTACTGCTAAG
TCTACCAGTATTAGCCGCTGGAATTACAATGCTACTAACGGATCGAAACTTTAATACAAG
TTTCTTTGACCCCGCTGGAGGTGGTGATCCGGTTTTATTTCAGCATATCTTT

>NC_035740

TACTTTATATCTTATTTTTGGGGCATTTTCCGCTATGATAGGAACTGCCTTTAGTATGATTA
TAAGGCTAGAATTATCTGGGCCAGGATCAATGTTAGGTGATGACCAACTTTATAACGTAG
TTGTCACAGCACACGCATTAATAATGATATTTTTCTTTGTTATGCCTGTTTTAATAGGAGG
GTTTGGAAACTGGTTAGTTCCCCTTTATATAGGAGCTCCGGACATGGCCTTCCCTAGGTTA
AACAATATTAGTTTTTGGTTATTACCTCCAGCTTTATTATTACTATTAGGATCATCCTTAGT
TGAACAAGGAGCAGGAACAGGATGAACTATCTACCCTCCCCTTAGTTCTATACAAGCTCA
CTCAGGAGGGTCCGTTGATATGGCTATATTTAGCCTCCATTTAGCAGGAGCCTCCTCTATT
ATGGGAGCTATCAATTTTATTACAACAATTTTAAATATGAGAGCACCAGGAATGACTATG
GATAAATTACCTTTATTTGTTTGATCAGTTTTAGTTACCGCAATTCTTCTATTACTATCATT
ACCTGTTTTAGCTGGTGCAATTACTATGTTATTAACAGATAGAAACTTTAATACTTCTTTC
TTTGACCCAGCTGGAGGGGGAGACCCAATATTATTTCAACATTTATTT

>NC_035979

TACTCTATATTTAATTTTCGGTGCCATTGCTGGAGTAATGGGTACATGCTTTTCAGTACCA
ATTCGTATGGAATTAGCACAACCCGGCAATCAAATTCTTGGTGGAAATCATCAACTTTAT
AATGTGTTAATAACAGCTCACGCTTTTTAATGATCTTCTTTATGGTTATGCCGGCGATGA
TAGGTGGTTTTGGTAATTGGTTCGTTCCCATTCTTATAGGAAGTCCGGATATGGCATTTCC
TAGATTAAATAATATTCCATTTTGGCTTTTGCCACCGTCATTGTTACTTCTTCCGAGCTCAG
CCTTAGTAGAGGTGGGTTGCGGTTCGGGGTGGACGGTCTATCCACCCTTAAGTGGTATAA
CCAGTCATTCCGGAGGATCTGTTGATTTAGCTATTTCTAGCCCTCATTTATCAGGTGTTTC
CTCTATTTTAGGTTCTATTAATTTTATAACAACTATCTTCAATATGAGGGGCCCCGGATTG
ACCATGCATAGATTACCTCTATTTGTGTGGTCTGTTTCAGTCACAGCTTTCCTACTTTTATT
ATCCCTTCCAGTATTGGCAGGTGCAATTACCATGTTATTAACTGATAGAAACTTTAATACA
ACCTTTTTCGATCCTGCCGGTGGCGGGGATCCCATTTTATACCAGCATCTTTTC

>NC_036051

TACTTTATATATGATATTTGGTATTTGAGCAGGTTTAGTGGGGACTGCTTTTAGATTTTTA
ATTCGGTCTGAGTTAGCTCAACCGGGTAGCTTATTAAATGATCCTCATTTATATAATAGTA
TAATTACGGCTCATGGATTAATTATGATTTTTTTTTTCGTAATGCCGGTCTTAATTGGGGG
TTTCGGTAAATGGTTAATTCCTTTATATTTGACTAGCCCAGATATGGCTTTTCCTCGGTTA
AAAAATATGAGTTTTTGGTTGTTGCCCCCTTCCTTTTTCCTTCTTTTAGGTTCTTTTGTTGTT
GAGAGGGGTGTTGGTACAGGTTGAACTCTTTATCCACCCTTGTCTGCTAATATAGCTCAA
AGGGGACCTAGAGTGGATATGGCTATTTTTTCTTTACATTTGGCTGGGGCTAGTTCTATTT
TAGGTTCTATAAATTTTATTACTACTATGGTTAATGCTAAGATCCAAGTTTCTTGGGGTCA
ATTGCCTTTGTTTTTGTGGGCAGTGATGGTTACTGCTTATATGTTAGTTCTTTCTTTACCCG
TTTTAGCTGGTGGGTTAACTATGTTATTAACTGATCGGAATTTTAATACTACTTTTTTTGAT
CCTGGTGGGGGTGGTGATCCTATATTATGGCAACATTTGTTT

>NC_036679

GACTATTTACTTGTATATAGGGCTTTGATCTGGGGTATTTGGTCTTAGATTAAGCCACTGT
ATACGGATTGAACTTAGCCATCCTGGAGAATGGCTTCAAGTTGGGTATATATACCATAGG
ATTATAACTATACATGCTTTTATGATAATTTTTTTTTTGTTATGCCCACAAGAATTGGAG
GATTGGGAAACTGGTTTATTCCATTAATGATTAAGATTAAAGACTTATCTATACCTCGATT
AAATAACTTAAGAGTGTGACTAGCATTAGGGTCTTTATTTCTTATGTGTATGGCTTTTATA
AGGAGAGGAGGGTTAGGTTGTGGTTGAACTATGTATCCACCCCTAAGGAATAGGGAGTTT
ATAGATGGATTACCTGTTGATTTGGCAGTATTCTCTCTCCACATAGCAGGGATATCATCAA
TTGCTGGAAGAATTAATTTTTTGGTGACTATTTTTAATATGCGTATAGGAGCTCTCTTTTTT
ATGAGGTTGAACCCTATGCTAATTTGGACTCTGTTTGGAACATCAATTTTATTGGTTACAT

CAGTTCCTGTGTTGGCTGCAGGTTTAACCTTATTATTATTAGATCGGCATTTTAGGACTAG
GTTTTATTACCCAGAGGGTGGTGGTGATCCAATCCTCTGGCAGCATTTA

>NC_037187

TACACTATATTTCTTATTTAGCATATGAGCAGGATTAATCGGAGCCATACTCAGCTTTTTA
ATTCGATTAGAACTCTCATCCCCCAGACCTACCTTCTTTAATACAGAGCTATTTAACATTA
CCATTACCGCCCACGCTCTTATCATAATCTTCTTTCTTGTAATACCAGCAATAATAGGAGG
ATTCGGGAACTGACTCATCCCTATAATACTCGGCATCCCAGATATAGCCTTCCCACGATTA
AACAACTTAAGATTTTGACTCCTACCGCCAGCCTTTATCCTCCTTACTCTTAGCATCCTTTA
TTTAAACGGCCCTAACACAGGCTGAACTATCTATCCCCCTCTCTCTAGCATCTCTGCAACA
TCATCTCCTAGTATTGATCTAGCCATTCTCTCTCTTCATATTGCTGGTGCCTCTTCCATCAT
AAGCTCAATCAACTTCATTACAACCATCCTTAACATGCACTCCCACAAGTCTTCCCTGTCC
CAAGTTCCACTCTTCCCTTGATCCATCATGATTACTGCCTTTCTAATCCTCCTCGCTATACC
CGTCCTCGCAGGAGCCATCACGATACTCCTTACAGATCGTAACTTCAATACTTCATTTTTC
GACCCTACTGGTGGAGGAGACCCTATCTTATTCCAACACTTATTC

>NC_037304

GACTCTCTATTTCATTTTCGGTGCCATTGCTGGAGTGATGGGCACATGCTTCTCAGTACTG
ATTCGTATGGAATTAGCACGACCCGGCGATCAAATTCTTGGTGGGAATCATCAACTTTAT
AATGTTTTAATAACAGCTCATGCTTTTTTAATGATCTTTTTTATGGTTATGCCGGCGATGAT
AGGTGGATTTGGTAATTGGTTTGTTCCGATTCTGATAGGTGCACCTGACATGGCATTTCCA
CGATTAAATAATATTTCATTCTGGTTGTTGCCACCAAGTCTCTTGCTCCTATTAAGCTCAG
CCTTAGTAGAAGTAGGTAGCGGCACTGGGTGGACGGTCTATCCGCCCTTAAGTGGTATTA
CCAGTCATTCTGGAGGAGCAGTTGATTTAGCAATTTTTAGTCTTCATCTATCTGGTGTTTC
ATCCATTTTAGGTTCTATCAATTTTATAACAACTATCTTCAACATGCGTGGACCTGGAATG
ACTATGCATAGATTACCCCTATTTGTGTGGTCCGTTCTAGTGACAGCATTCCTACTTTTATT
ATCACTCCCGGTACTGGCAGGGGCAATTACCATGTTATTAACCGATCGAAACTTTAATAC
AACCTTTTTTGATCCCGCTGGAGGGGGAGACCCAATTTTATACCAGCATCTCTTT

>NC_037431

TACTTTGTATTTATTATTAGCTTTGTGGTCTGGTTTAATTGGGTTGGCTTTAAGACTTTTAA
TCCGAGCTGAGTTAGGGCAACCTGGAAGGTTGTTAGGAGATGACCAGTTATACAATGTGA
TTGTTACAGCTCATGCTTTTATAATAATTTTTTTTTTAGTAATACCTATGATAATTGGTGGT
TTTGGTAATTGGCTTATTCCTTTAATAATTGGCGCTCCTGATATGGCTTTTCCTCGACTAAA
TAATTTAAGTTTTTGGCTTCTTGTACCAGCTTTGTTTTTGTTATTAAGTTCTTCGTTGGTTG
AAAGGGGTGTGGGGACTGGGTGAACAGTGTATCCTCCTTTATCTGGAAATGTTGCTCATT
CCGGGGCTTCGGTTGATTTAGCTATTTTTTCTTTACATCTTGCTGGTGCTTCTTCAATTTTA
GGTGCTATTAATTTTATTTCTACTGTTGGAAATATGCGGTCTCCTGGTTTGGTAGCTGAGC

GAATTCCTTTATTTGTTTGAGCTGTTACGGTAACAGCTATTTTGTTAGTTGCTGCTTTACCT
GTTTTAGCTGGTGCCATTACTATACTTCTTACTGATCGTAATTTAAATACTTCGTTTTTTGA
CCCTACGGGAGGAGGTGATCCTATTTTGTATATGCATTTGTTT

>NC_037463

GACTCTCTATTTCATCTTCGGTGCCATTGCTGGAGTGATGGGCACATGCTTCTCAGTACTG
ATTCGTATGGAATTAGCACGTCCCGGCGATCAAATTCTTGGTGGGAATCATCAACTTTAT
AATGTTTTAATAACGGCTCACGCTTTTTTAATGATCTTTTTTATGGTTATGCCGGCGATGA
TAGGTGGATCTGGTAATTGGTCTGTTCCGATTCTGATAGGTGCACCTGACATGGCATTTCC
ACGATTAAATAATATTTCATTCTGGTTGTTGCCACCAAGTCTCTTGCTCCTATTAAGCTCA
GCCTTAGTAGAAGTGGGTAGCGGCACTGGGTGGACGGTCTATCCGCCCTTAAGTGGTATT
ACCAGCCATTCTGGAGGAGCAGTTGATTCAGCAATTTCTAGTCTTCATCTATCTGGTGTTT
CATCCATTTTAGGTTCTATCAATTTTATAACAACAATCTCCAACATGCGTGGACCTGGAAT
GACTATGCATAGATCACCCCTATTTGTGTGGTCCGTTCTAGTGACAGCATTCCCACTTTTA
TTATCACTTCCGGTACTGGCGGGGGCAATTACCATGTTATTAACCGATCGAAACTTTAATA
CAACCTTTTCTGATCCCGCTGGAGGGGGAGACCCCATATTATACCAGCATCTCTTT

>NC_037468

GACTCCATATTTAATCTTCGGTGCCATTGCTGGAGTGATGGGCACATGCTTCTCCGTACTG
ATTCGTATGGAATTAGCACGACCCGGCGATCAAATTCTTTGTGGGAATCATCAACTTTAT
AATGTTTTAATAACGGCTCACGCTTTTTTAATGATCTTTTTTTTTGTTATGCCGGCGATGAT
AGGTGGATCTGGTAATTGGTCTGTTCCGATTCTTATAGGTGCACCTGACATGGCATTTCCA
CGATTAAATAATATTTCATTCCGGTTGTTGCCACCAAGTCTCTTGCTCCTATTAAGCCCAG
CCTTAGTAGAAGCGGGTAGCGGCACTGGGTGGACGGTCTATCCGCCCTTAAGTGGTATTA
CCAGCCATGGAGGAGGAGCAGTTGATTTAGCAATTTCTAGTCTTCATCTATCAGGTGTTTC
ATCCATTTTAGGTTCTATTAATTTTATAACAACTATCTCCAACATGCGCGGACCTGGAATG
ACTATGCATAGATCACCCCTCTTTGTGTGGTCCGTTCTAGTGACAGCATTCCTACTTTTATT
ATCACTTCCGGTACTGGCGGGGGCAATTACCATGTTATTAACCGATCGCAACTTTAATAC
AACATTTTCTGATCCCGCTGGAGGGGGAGACCCCATATTATACCAGCATCTCTTT

>NC_037610

AACCATATATTTAATTTTTGGTGTTTGATCAGCCATAGTTGGGACAGCATTTAGTGTTCTT
ATCCGACTTGAATTAGGTCAACCAGGATCTTTCATTGGAGACGATCAAATTTATAATGTC
ATAGTTACAGCCCATGCTTTTATTATAATTTTTTTTATAGTTATACCAATCATAATTGGAG
GGTTTGGAAATTGACTTGTACCTTTAATAATTGGAGCCCCAGATATAGCCTTCCCCCGAAT
AAATAACATAAGATTCTGACTTCTCCCTCCTTCACTTATTCTTTTATTATCAGGAGGACTG
GTGGAAAGAGGGGCTGGGACAGGATGAACAGTTTACCCCCCTCTTTCAGCAGGAATTGC
GCATGCCGGAGCTTCCGTTGATTTATCAATTTTTAGACTTCATCTAGCAGGGGCTTCTTCA

ATTCTAGGGGCAGTAAATTTTATTACAACAATTATTAATATACGCTCGGCTGGTATATCGT
GAGATCGAACACCTTTATTTGTTTGATCAGTATTATTAACAGCCATTCTTCTTCTACTATCT
CTCCCAGTCTTAGCTGGGGCAATCACTATACTTTTAACAGATCGTAATTTAAATACATCTT
TCTTTGATCCTGCTGGAGGAGGAGACCCTATTTTATACCAACATTTATTT

>NC_037891

TACCTTATATTTAATTTTTGGTGCTTTTTCTGGTATATTAGGCGGATGCATGTCAATGTTAA
TTCGTATGGAATTAGCCCAACCTGGAAATCAATTATTGTTAGGAAATCATCAAATTTATA
ATGTTTTAATTACAGCACATGCTTTTTTAATGATTTTTTTTATGGTAATGCCTGTTATGATT
GGAGGGTTTGGAAATTGGTTAGTTCCTATTATGATAGGTAGTCCAGATATGGCCTTTCCTC
GTTTAAATAATATATCATTTTGATTATTACCACCATCTCTTTGTTTACTTTTAGCATCAGCA
ATTGTGGAAGTAGGTGTAGGTACAGGATGAACTGTATACCCTCCACTTAGTTCAATTCAA
AGCCATTCTGGTGGTGCTGTAGACTTAGCGATTTTCAGTTTACATATTTCGGGTGCATCTT
CAATTTTAGGTGCTATTAATTTTATTTCTACGATTTTAAATATGCGTAATCCAGGACAAAG
TATGTATCGTATGCCTTTGTTTGTATGATCAATTTTTATTACTGCATTTTTATTATTATTAG
CTGTTCCTGTTTTAGCTGGTGCTATTACAATGCTTTTGACTGATCGTAATTTTAATACAGC
ATTTTTTGATCCTGCAGGAGGAGGAGATCCTGTATTATATCAGCATCTTTTT

>NC_037936

TACTCTTTATTTAATTTTCTCAGTTTTTGCTGGTATGATAGGTACTGCATTTTCTGTATTAA
TTAGATTAGAATTATCAGCACCTGGAGTGCAAGTTTTACAAGGAGACCATCAATTATTTA
ATGTGATCATTACTGCACATGCCTTTATAATGATCTTCTTTATGGTTATGCCTGCTTTAGTA
GGAGGTTTTGGTAACTATTTATTACCAGTTCAAGTGGGAGCCCCTGATATGGCTTTCCCGA
GATTAAATAATATCTCATTCTGGTTATTACCTCCTTCATTAATTTTATTATTATTAAGTGCT
TTAGTAGAAAATGGAGCTGGTACAGGATGGACAGTTTATCCTCCTTTAGCTGGTATCCAA
TCACACTCTGGTGCATCTGTTGATTTAGCTATCTTCAGTTTACACTTAGCTGGGGTATCTTC
TTTATTAGGGGCTATCAATTTCATTACTACTGTACTTAATATGAGAACTAATGGAATGAGT
TTACACAAATTACCATTATTTGTATGGGCAATATTTATTACTGCAATACTATTATTATTAT
CATTACCAGTATTAGCCGGTGCAATTACAATGTTATTAACAGATAGAAACTTTAATACTA
GTTTCTATGATCCAGCCGGAGGTGGTGATCCAATCTTATATCAACACTTATTC

>NC_037988

AACTTTGTATTTAATTTTTGGCGCAATGTCTGGTGTGGCTGGAACTGCTTTATCTTTATTTA
TTCGAATAACATTATCACAACCTGATAACAATTTTTTATCGTACAATCATCAATTATACAA
TGTTATAGTGACTGGTCACGCATTCATAATGATTTTTTTTATGGTTATGCCTACGTTAATTG
GTGGTTTTGGAAATTGGTTTGTACCGTTGATGATTGGAGCGCCTGACATGGCTTTTCCTAG
AATGAATAATATTAGTTTTTGGCTTTTACCACCATCATTACTTTTATTAATTTCTTCAATAC
TTGCAGAAGCAGGGGCTGGAACTGGTTGGACTGTTTATCCGCCACTATCAAGCGGTAGCT

CGCACTCAGGCGGTGCCGTGGATTTAGCAATTTTTAGCTTACACTTGTCGGGTGCGTCATC
AATTTTAGGAGCGATCAATTTTATATGCACTATAGTTAATATGCGTACAAAAAGTTTATTT
TTTCATCAATTACCTTTATTTGTTTGGTCTGTATTAATTACAGCATTTTTATTGTTATTATC
ATTACCAGTTTTAGCGGGCGCTATCACAATGCTTTTGACTGATCGTAATTTTAATACTACA
TTTTTTGATCCGGCAGGTGGTGGGGATCCAGTATTATATCAACATTTATTT

>NC_037990

AACCTTATATATTATTTTTGGTGCTATTGCTGGTGTAGCTGGTACAGCTTTATCTCTATATA
TTAGACTTACATTAGCACAACCAAATGGAGGTTTCTTAGAATATAATCACCATTTCTATAA
TGTAATTGTAACAGGTCATGCATTTATTATGATTTTTTTCATGGTAATGCCTGTTTTAATTG
GAGGATTTGGAAACTGGTTCGTTCCTTTAATGATTGGAGCACCAGATATGGCATTCCCAA
GAATGAATAACATTAGTTTTTGGTTATTACCTCCTTCTCTTCTATTATTAGTTGAGTCTGTA
CTTTGTGAAGCAGGTGTAGGTACCGGTTGGACAGTTTACCCTCCATTATCTGGTATCATTG
CACACTCTGGTGGTTCTGTAGATCTTGCTATCTTCAGTCTTCACTTATCAGGAGCTGCTTCT
ATTTTAGGTTCAATTAATTTTATTTGTACAGTTGCAAATATGAGAACTAATAGTTTACCAT
TCCATAAAATTCCTTTATTCGTTTGGTCTGTTTTCATTACTGCTATTCTATTATTATTATCAT
TACCTGTATTAGCTGGAGCTATTACTATGTTATTAACTGATAGAAATTTCAACACTACTTT
CTTTGATCCTGCAGGTGGTGGTGATCCTGTATTATACCAGCATTTATTC

>NC_039455

TACTCTTTACCTTATCTTTGCTGTATTCTCAGGACTACTAGGTACAGGATTCTCAATCCTA
ATTCGTGCTGAACTAGCTGCGCCTGGTGTTCAAGTTCTGGGAGGAAATCATCAACTATTT
AATGTGATTATTACAGCTCACGCATTCCTAATGGTATTCTTCCTAGTAATGCCAGCTCTAA
TGGGTGGATTCGGTAACTACTTCACTCCTGTAATGGTAGGTGCTGTAGATATGGCTTTCCC
ACGACTAAATAACATTAGTTTCTGGCTACTAGTACCTAGCCTAATTCTACTACTAGCAAGT
GCTTTCGTAGAACAAGGGGCTGGATCAGGATGGACAGTTTATCCACCACTGGCAGGACTA
CAAAGTCACTCAGGAGGTTCAATTGATCTAGCTATCTTTAGTCTTCACCTATCAGGTATTT
CATCTATGCTAGGTGCTATGAACTTTATGGTAACTATCCTAAATATGCGAGCTTCAGGTAT
GACTATGTACAAAATGCCACTATTCGTATGGTCAGTATTTGTAACATCAATTCTACTAATC
ACTACACTACCTGTACTAGCAGGTGCTATTACAATGCTACTAACTGACCGTAACTTTAATA
CATCATTCTATGATCCAGCAGGAGGTGGTGATCCAGTACTATATCAACATCTATTC

>NC_039746

GACTCCACATTCAATCTTCGGTGCCATTGCTGGAGTAATGGGCACATGCTTCTCAGTATCA
ATTCGTATGGAATTAGCACAACCCGGCGATCAAATTCTTGGTGGGAATCATCAACCTCAT
AATGTGTCAATAACGGCTCACGCTTCTCCAATGATCCCTTTTATGGTTATGCCGGCGGTGA
TAGGTGGATCTGGTAATTGGTCCGTTCCGATTCCTATAGGTGCACCTGACATGGCATTTCC
ACGATTGAATAATATTCCATCCCGGTTGTTGCCACCTTCGCTGTTGCTCCCATTAAGCCCA

GCCTCGGTAGAAGTGGGTAGCGGCACTGGGTGGACGGTCTATCCGCCCCTAAGTGGTATT
ACCAGTCATTCCGGAGGAGCTGCTGATCCAGCGATTTCTAGTCCTCATCTATCAGGTGTTT
CATCCATTTCAGGTTCTATCAATCTCATAACTACTATCCCCAACATGCGCGGGCCTGGAAT
GACTATGCATAGATCACCCCTATTTGTGCGGTCCGTTCCAGTGACAGCATTCCTACTCTTA
TCATCACTTCCGGTACCGGCAGGGGCAATTACCATGTTATCAACTGATCGAAGCTTTAAT
ACAACCTTTTTCGATCCTGCTGGAGGGGGAGACCCGATATTATACCAGCATCTCTTT
