## Supplementary material for "VTAM: A robust pipeline for validating metabarcoding data using internal controls": SuppInfo2.pdf

### SuppInfo2: Detailed protocol of the bioinformatics analyses of benchmarking.

VTAM was tested on two published datasets of metabarcoding : The fish dataset (Corse et al., 2017) contained 35 samples of excrement for *Zingel asper*, a fresh-water species, and 5 samples of excrement from *Pomatoschistus microps*, from brackish water. The bat dataset (Galan et al., 2018) had 357 bat fecal pellet samples from different bat species. Both datasets included negative controls (6 for fish and 19 for the bat dataset) and mock samples (2 for the fish and 24 for bat) with known composition. All samples in both studies had three PCR replicates. The fish samples were amplified by two primer sets (markers: MFZR and ZFZR), amplifying the first 158-182 bases of the COI gene (Meusnier et al., 2008; Zeale, Butlin, Barker, Lees, & Jones, 2011), while from the bat samples a 133-bp minibarcode of the COI (Gillet et al.,2015) was amplified. The detailed description of samples and laboratory protocols are found in the original studies (Corse et al., 2017; Galan et al., 2018) and the full raw datasets are available in Dryad (https://datadryad.org/stash/dataset/doi:10.5061/dryad.f40v5, https://datadryad.org/stash/dataset/doi:10.5061/dryad.kv02g).

**User input files are found here:**
http://net.imbe.fr/~emeglecz/SuppInfo_VTAM/user_input_files/

**Final ASV tables are found here:** http://net.imbe.fr/~emeglecz/SuppInfo_VTAM/ASV_tables/

**Custom scripts to obtain strictly equivalent input files for the VTAM and DADA2 based pipelines are found here:** http://net.imbe.fr/~emeglecz/SuppInfo_VTAM/scripts/

### VTAM

#### Fish dataset

The fish dataset was run separately for the two markers and then the results are pooled in one ASV table.

Move to the vtam directory and activate the Conda environment

```
cd vtam-0.1.12
conda activate vtam
```

#### merge, demultiplex (sortreads) , filter by default parametes and taxassign for MFZR and ZFZR

```
vtam merge --fastqinfo fish/user_input/fastqinfo_fish.tsv --fastqdir
fish/fastq --fastainfo fish/fastainfo_fish.tsv --fastadir fish/merged -
v --log fish/vtam.log

vtam sortreads --fastainfo fish/fastainfo_fish.tsv --fastadir
fish/merged --outdir fish/sorted -v --log fish/vtam.log

vtam filter --db fish/db.sqlite --readinfo fish/sorted/readinfo.tsv --
readdir fish/sorted --asvtable fish/asvtable_default.tsv -v --log
fish/vtam.log --lfn_variant_replicate

vtam taxassign --db fish/db.sqlite --variants fish/asvtable_default.tsv
--output fish/asvtable_default_taxa.tsv --taxonomy vtam_db/taxonomy.tsv
--blastdbdir vtam_db/coi_blast_db --blastdbname coi_blast_db_20200420 -
v --log fish/vtam.log
```

Create **known_occurrences_fish.tsv** including keep and delete occurrences of both markers.

#### Optimize

##### OptimizePCRerror and OptimizeLFNbiosampleReplicate

```
vtam optimize --db fish/db.sqlite --readinfo fish/sorted/readinfo.tsv -
-readdir fish/sorted --known_occurrences
fish/user_input/known_occurrences_fish.tsv --outdir fish/optimize -v --
log fish/vtam.log --until OptimizePCRerror
```

```
vtam optimize --db fish/db.sqlite --readinfo fish/sorted/readinfo.tsv -
-readdir fish/sorted --known_occurrences
fish/user_input/known_occurrences_fish.tsv --outdir fish/optimize -v --
log fish/vtam.log --until OptimizeLFNbiosampleReplicate
```

We have chosen the same values for mfzr and zfzr markers for the following parameters, and created a parameter file for OptimizeLFNreadCountAndLFNvariant: **params_optimize_fish.yml**

```
lfn_biosample_replicate_cutoff: 0.003
pcr_error_var_prop: 0.1
```

##### OptimizeLFNreadCountAndLFNvariant

```
vtam optimize --db fish/db.sqlite --readinfo fish/sorted/readinfo.tsv -
-readdir fish/sorted --known_occurrences
fish/user_input/known_occurrences_fish.tsv --outdir fish/optimize -v --
log fish/vtam.log --until OptimizeLFNreadCountAndLFNvariant --params
fish/user_input/params_optimize_fish.yml
```

We have chosen the same values for mfzr and zfzr markers for all parameters, and created a parameter file for filter: **params_filter_fish.yml**

```
lfn_biosample_replicate_cutoff: 0.003
pcr_error_var_prop: 0.1
lfn_variant_replicate_cutoff: 0.006
lfn_read_count_cutoff: 10
```

#### Filter with optimized parameters

```
vtam filter --db fish/db.sqlite --readinfo fish/sorted/readinfo.tsv --
readdir fish/sorted --asvtable fish/asvtable_optimized.tsv -v --log
```

```
fish/vtam.log --lfn_variant_replicate --params
fish/user_input/params_filter_fish.yml
```

##### pool

Pool sequences of the two markers if identical in their overlapping region

```
vtam pool --db fish/db.sqlite --runmarker
fish/user_input/pool_run_marker_fish.tsv --output
fish/asvtable_pooled_mfzr_zfzr_fish.tsv --log fish/vtam.log -v

vtam taxassign --db fish/db.sqlite --variants
fish/asvtable_pooled_mfzr_zfzr_fish.tsv --output
fish/asvtable_vtam_pooled_mfzr_zfzr_fish_taxa.tsv --taxonomy
vtam_db/taxonomy.tsv --blastdbdir vtam_db/coi_blast_db --blastdbname
coi_blast_db_20200420 -v --log fish/vtam.log
```

#### Bat dataset

The bat dataset is already demultiplexed, there is one fastq file pair for each sample-replicate.

##### Trim primers

Primers were trimmed from the unmerged fastq files using cutadapt v2.10 (Martin, 2011) run by a custom perl script (**cutadapt_galan.pl**). Reads were kept only if a primer was found both on the fw and reverse reads. Trimmed reads were discarded if shorter than 100 bp After primer trimming reads are shortened to 120 bp, to avoid part of the primer at the 3' of the sequence.

Files were renamed to obtain homogeneous nomenclature by a custom perl script: **reformat_galan_tsv.pl** (These files were also the input of the DADA based pipeline)

##### merge read pairs

Paired end reads were merged by vsearch –fastq_mergepairs (Rognes et al., 2016) option ran from **merge_galan.pl** custom perl script.

#### filter and taxassign

```
vtam filter --db bat/db.sqlite --readinfo
bat/user_input/fastainfo_bat.tsv --readdir bat/fasta_merged --asvtable
bat/asvtable_default.tsv -v --log bat/vtam.log

vtam taxassign --db bat/db.sqlite --variants bat/asvtable_default.tsv
--output bat/asvtable_default_taxa.tsv --taxonomy vtam_db/taxonomy.tsv
--blastdbdir vtam_db/coi_blast_db --blastdbname coi_blast_db_20200420 -
v --log bat/vtam.log
```

#### optimize

##### OptimizePCRerror and OptimizeLFNbiosampleReplicate

```
vtam optimize --db bat/db.sqlite --readinfo
bat/user_input/fastainfo_bat.tsv --readdir bat/fasta_merged --
known_occurrences bat/user_input/known_occurrences_bat.tsv --outdir
bat/optimize -v --log bat/vtam.log --until OptimizePCRerror

vtam optimize --db bat/db.sqlite --readinfo
bat/user_input/fastainfo_bat.tsv --readdir bat/fasta_merged --
known_occurrences bat/user_input/known_occurrences_bat.tsv --outdir
bat/optimize -v --log bat/vtam.log --until
OptimizeLFNbiosampleReplicate
```

**params_optimize_bat.yml**

```
lfn_biosample_replicate_cutoff: 0.002
pcr_error_var_prop: 0.2
```

##### OptimizeLFNreadCountAndLFNvariant

```
vtam optimize --db bat/db.sqlite --readinfo
bat/user_input/fastainfo_bat.tsv --readdir bat/fasta_merged --
known_occurrences bat/user_input/known_occurrences_bat.tsv --outdir
bat/optimize -v --log bat/vtam.log --until
OptimizeLFNreadCountAndLFNvariant --params
bat/user_input/params_optimize_bat.yml
```

**params_filter_bat.yml**

```
lfn_biosample_replicate_cutoff: 0.002
pcr_error_var_prop: 0.2
lfn_variant_cutoff:  0.006
lfn_read_count_cutoff: 60
```

##### Filter with optimized parameters

```
vtam filter --db bat/db.sqlite --readinfo
bat/user_input/fastainfo_bat.tsv --readdir bat/fasta_merged --asvtable
bat/asvtable_optimized_bat.tsv -v --log bat/vtam.log --params
bat/user_input/params_filter_bat.yml

vtam taxassign --db bat/db.sqlite --variants
bat/asvtable_optimized_bat.tsv --output
bat/asvtable_vtam_optimized_bat_taxa.tsv --taxonomy
vtam_db/taxonomy.tsv --blastdbdir vtam_db/coi_blast_db --blastdbname
coi_blast_db_20200420 -v --log bat/vtam.log
```

#### DADA based pipeline

This pipeline is composed of three main steps:

- DADA2 (Callahan et al., 2016) denoising algorithm was used to eliminate the majority of the artifacts
- LULU (Frøslev et al., 2017) to eliminate further false positive occurrences
- Equivalent of FilterMinReplicateNumber of VTAM, to pool occurrences over replicates was done by a custom perl script (**pool_replicates.pl**).

### Fish dataset

#### creating fastq file pairs per sample-replicate

The Fish dataset consists of fastq files that needs to be demultiplexed and trimmed from primers and tags to provide one fastq file pairs per sample-replicate as input to DADA2. To ensure to comparability between the DADA based pipeline and the VTAM filtering, we have used the demultiplexed fasta files produced by sortreads of VTAM, to recreate the input fastq file pairs for DADA2, using the **make_dada_fastq.pl** custom perl script.

##### trim adapters and tags from fastq and orient sequences

Fastq sequences have tags and primers, and the orientation is random (Forward reads can start by the forward or reverse primer). We used the **cutadapt_primers_for_dada.pl** custom perl script to trim fastq sequences. This script runs cutadapt2.10 for each file pairs (--discard-untrimmed; --no-indels; --minimum-length min_read_length; --length max_read_length; -e 0.1; minimum overlap length of the primer)

The script has been run twice for each marker to also identify read-pairs in the wrong orientation.

```
#### MFZR orientation_fw
my $dir = 'fish/fastq_demultiplexed_non_trimmed/';
my $motif = 'MFZR.+_fw.fastq';
my $fw = 'TCCACTAATCACAARGATATTGGTAC';
my $rev = 'WACTAATCAATTWCCAAATCCTCC';
my $min_read_length = 160;
my $max_read_length = 170;
my $outdir = $dir.'MFZR_orientation_fw/';

#### MFZR orientation_rev
my $dir = fish/fastq_demultiplexed_non_trimmed/';
my $motif = 'MFZR.+_fw.fastq';
my $fw = 'WACTAATCAATTWCCAAATCCTCC';
my $rev = 'TCCACTAATCACAARGATATTGGTAC';
my $min_read_length = 160;
my $max_read_length = 170;
```

```
my $outdir = $dir.'MFZR_orientation_rv/';

#### ZFZR orientation_fw
my $dir = fish/fastq_demultiplexed_non_trimmed/';
my $motif = 'ZFZR.+_fw.fastq';
my $fw = 'WACTAATCAATTWCCAAATCCTCC';
my $rev = 'AGATATTGGAACWTTATATTTTATTTTTGG';
my $min_read_length = 140;
my $max_read_length = 150;
my $outdir = $dir.'ZFZR_orientation_rev/';
```

**orient_fastq_filepairs.pl** custom perl script were run to pool and orient file pairs

```
#### MFZR
my $dir_fw =
'fish/fastq_demultiplexed_non_trimmed/MFZR_orientation_fw/';
my $dir_rev =
'fish/fastq_demultiplexed_non_trimmed/MFZR_orientation_rev/';
my $motif = '_fw.fastq';
my $outdir = 'fish/demultiplexed/MFZR/';

#### ZFZR
my $dir_fw =
'fish/fastq_demultiplexed_non_trimmed/ZFZR_orientation_fw/';
my $dir_rev =
'fish/fastq_demultiplexed_non_trimmed/ZFZR_orientation_rev/';
my $motif = '_fw.fastq';
my $outdir = 'fish/demultiplexed/ZFZR/';
```

#### DADA2

DADA2 was run separately for MFZR and ZFZR by the **dada2.R** script

**reformat_dada_asv_table_for_lulu.pl** was used to reformat the dada output table for LULU and produce BLAST hit table

##### LULU

LULU was run separately for MFZR and ZFZR by the **lulu.R** script

##### pool replicates

**pool_replicates.pl** Pool replicates from lulu output: accept occurrences if the ASV is present in at least two replicates of the sample

##### taxassign by VTAM

```
vtam taxassign --db fish/db.sqlite --variants
fish/lulu_output/MFZR/mfzr_dada_lulu_pooled_2repl.tsv --output
fish/lulu_output/MFZR/mfzr_dada_lulu_pooled_2repl_taxa.tsv --taxonomy
vtam_db/taxonomy.tsv --blastdbdir vtam_db/coi_blast_db --blastdbname
coi_blast_db_20200420 -v --log fish/vtam.log
```

##### pool_markers_dada.pl

Pool results from the two markers => output: **asvtable_dada_lulu_pooled_2repl_fishtaxa.tsv**

#### Bat dataset

The bat dataset is already demultiplexed, there is one fastq file pair for each sample-replicate.

##### Trim primers

Primers were trimmed from the unmerged fastq files using cutadapt v2.10 (Martin, 2011) run by a custom perl script (**cutadapt_galan.pl**). Reads are kept only if a primer was found both on the fw and reverse reads. Trimmed reads were discarded if shorter than 100 bp After primer trimming reads are shortened to 120 bp, to avoid part of the primer at the 3' of the sequence.

Files were renamed to obtain homogeneous nomenclature by a custom perl script: **reformat_galan_tsv.pl** (These files are the same as the input of VTAM based pipline)

##### DADA2

See **dada2.R** script

**reformat_dada_asv_table_for_lulu.pl** was used to reformat the dada output table for LULU and produce BLAST hit table

##### LULU

See **lulu.R** script

##### pool replicates

**pool_replicates.pl** pool replicates from lulu output: accept occurrences if the ASV is present in at least 2 replicates of the sample

##### Taxassign by VTAM

```
vtam taxassign --db galan/db.sqlite --variants
bat/lulu_output/dada_lulu_pooled_2repl_bat.tsv --output
bat/lulu_output/asvtable_dada_lulu_pooled_2repl_bat_taxa.tsv --taxonomy
vtam_db/taxonomy.tsv --blastdbdir vtam_db/coi_blast_db --blastdbname
coi_blast_db_20200420 -v --log galan/vtam.log
```
