## Supplementary material for "VTAM: A robust pipeline for validating metabarcoding data using internal controls": SuppInfo3.pdf

### SuppInfo3: Diversity estimations based on VTAM and DADA2 ASV tables

Unless specified otherwise, analyses were conducted using R software (R development Core Team., 2017).

ASV tables: The output ASV tables of the two pipelines (found in Supporting Information 2) were used for the diversity estimation.

Cluster tables: Within each ASV table, ASVs were clustered using 3% divergence threshold and the clusters were considered as a proxy for delineating invertebrate species. We used the number of ASVs of each cluster in each sample as semiquantitative estimate of abundance (details in Corse et al., 2017).

#### Diversity comparison between VTAM and DADA2

The ASVs tables were used to compare the richness of the ASVs per sample obtained with VTAM and DADA2.

The cluster tables were used to compare the richness of cluster per sample obtained with VTAM and DADA2.

The cluster tables were also used to compare the beta diversity between pair of samples obtained with VTAM and DADA2. Beta diversity was assessed using the Bray-Curtis pairwise dissimilarity index using the diversity function of the vegan R package (Oksanen et al., 2013). Pairwise sample comparisons were conducted among samples of the same type (control, biological samples) and same host/predator (e.g. *Zingel asper* faeces; *Pomatoschistus microps* faeces; Bats faeces).

We performed additional analyses to study the correlation between ASVs and Cluster richness per sample. The differences between slopes of the two pipelines were tested using the *lstrend* function of the *lsmeans* R package (Lenth, 2016) and the *pairs* R base function with the following linear model:

Cluster_Richness = ASV_richness * Pipelines.

Slopes between VTAM and DADA2 were significantly different (estimate= -0.0586 SE=0.010; DF= 892; t.ratio=-5.601; Pvalue< 0.0001).
